# The dietary ellagitannin metabolite urolithin A is produced by a molybdenum-dependent dehydroxylase encoded by prevalent human gut *Enterocloster* spp

**DOI:** 10.1101/2024.02.08.579493

**Authors:** Reilly Pidgeon, Sacha Mitchell, Michael Shamash, Layan Suleiman, Lharbi Dridi, Corinne F. Maurice, Bastien Castagner

**Affiliations:** Department of Pharmacology & Therapeutics, McGill University, 3655 Prom. Sir-William-Osler, Montreal, Quebec, H3G 1Y6, Canada; Department of Microbiology & Immunology, McGill University, 3775 University Street, Montreal, Quebec, H3A 2B4, Canada; McGill Centre for Microbiome Research, Montreal, Quebec, Canada

## Abstract

Urolithin A (uroA) is a polyphenol derived from the multi-step metabolism of dietary ellagitannins by the human gut microbiota that can affect host health by stimulating mitophagy. Most individuals harbor a microbiota capable of uroA production; however, the mechanisms underlying the dehydroxylation of its catechol-containing dietary precursor (uroC) are unknown. Here, we use a combination of untargeted bacterial transcriptomics, proteomics, and comparative genomics to uncover an inducible uroC dehydroxylase (*ucd*) operon in *Enterocloster* spp. We show that *Enterocloster* spp. are sensitive to iron chelation by uroC, and dehydroxylation to uroA rescues growth by disrupting the iron-binding catechol. Importantly, only microbiota samples actively transcribing *ucd* could produce uroA, establishing *ucd*-containing *Enterocloster* spp. as keystone urolithin metabolizers. Overall, this work identifies *Enterocloster* spp. and the *ucd* operon as main contributors to uroA production and establishes a multi-omics framework to further our mechanistic understanding of polyphenol metabolism by the human gut microbiota.

## Main

The human gut microbiota is a collection of trillions of microorganisms that colonize the gastrointestinal tract and play pivotal roles in host health and disease ^1^. Gut bacteria help maintain homeostasis by regulating host immune cell activity, gut barrier integrity, and nutrient availability ^2^. One of the main mediators of microbiota-host interactions are microbial metabolites. Gut bacteria possess an immense metabolic repertoire (nearly 1000-fold more protein coding sequences than the human genome ^3^) to perform four main classes of reactions: hydrolysis, conjugation, cleavage, and reduction ^4–7^. These ubiquitous reactions have been linked to microbiota-dependent metabolism of therapeutic drugs ^8–10^, host bile acids ^6,11,12^, and diet-derived compounds ^13–15^.

Diet is a strong modulator of the composition and function of the gut microbiota ^16–19^. Diet-derived polyphenols are a diverse class of plant secondary metabolites found in fruits, vegetables, and nuts (reviewed in ^20^) that are poorly absorbed by the host and reach the large intestine relatively intact ^7,21^. Ellagitannins are a large sub-group of polyphenols that belong to the family of hydrolysable tannins and are characterized by a central glucose (open-chain or pyranose forms) linked to diverse pyrogallol-like moieties ^20^. Camu camu, a berry rich in the ellagitannin castalagin, has been shown to impact anti-cancer immunity via the gut microbiome, and is currently in clinical trials (NCT05303493, NCT06049576) in combination with immune checkpoint inhibitors ^22,23^. Depending on microbiota composition, ellagitannins can be hydrolyzed and reduced by gut bacteria into bioactive metabolites (ellagic acid, urolithins, nasutins) according to different metabolic phenotypes characterized by the terminal metabolites observed in biological fluids ^24^ (Supplementary Fig. 1).

Urolithin A (uroA) is the most common terminal metabolite of ellagitannin metabolism and has reported pharmacological activities both within the gut environment and systemically following absorption ^25^. In the gut, uroA can attenuate colitis by increasing the expression of epithelial tight junction proteins ^26–28^ *via* the activation of aryl hydrocarbon receptor (AhR)-Nrf2 pathways ^29,30^. Additionally, uroA can enhance immunotherapy in colorectal cancer models by activating Pink1-dependent mitophagy pathways in T cells, improving anti-tumor CD8+ T cell immunity ^31^. Clinical trials in healthy individuals have demonstrated that uroA is safe, bioavailable, and can be detected in its aglycone, glucuronidated, and sulfated forms in plasma ^25,27^. Once absorbed by the host, uroA can trigger mitophagy in muscle cells, improving muscle function in animal models of ageing and Duchenne muscular dystrophy ^26,29,32,33^. Overall, uroA can enhance gut barrier integrity, modulate the immune system, and promote mitochondrial health in the host, thus showing promise as a postbiotic to treat age-related conditions ^34–36^.

While urolithin metabolism is prevalent in human populations, few gut bacteria have been reported to metabolize urolithins ^34–36^. Most known urolithin metabolizers belong to the *Eggerthellaceae* family (*Gordonibacter urolithinfaciens*, *Gordonibacter pamelaeae*, *Ellagibacter isourolithinifaciens*) and can perform multiple metabolic steps in the urolithin metabolism pathway, yielding either urolithin C (uroC) or isourolithin A (isouroA) from ellagic acid ^37^. Recently, certain members of the *Enterocloster* spp. (*Lachnospiraceae* family) were reported to dehydroxylate uroC to uroA and isouroA to urolithin B (uroB) both *in vitro* and *in vivo* ^38,39^. These findings shed light on the minimal bacterial community required for the complete metabolism of ellagic acid to uroA; however, the genes and enzymes responsible for these dehydroxylation reactions remain unknown (Fig. 1A).

**Figure 1.**
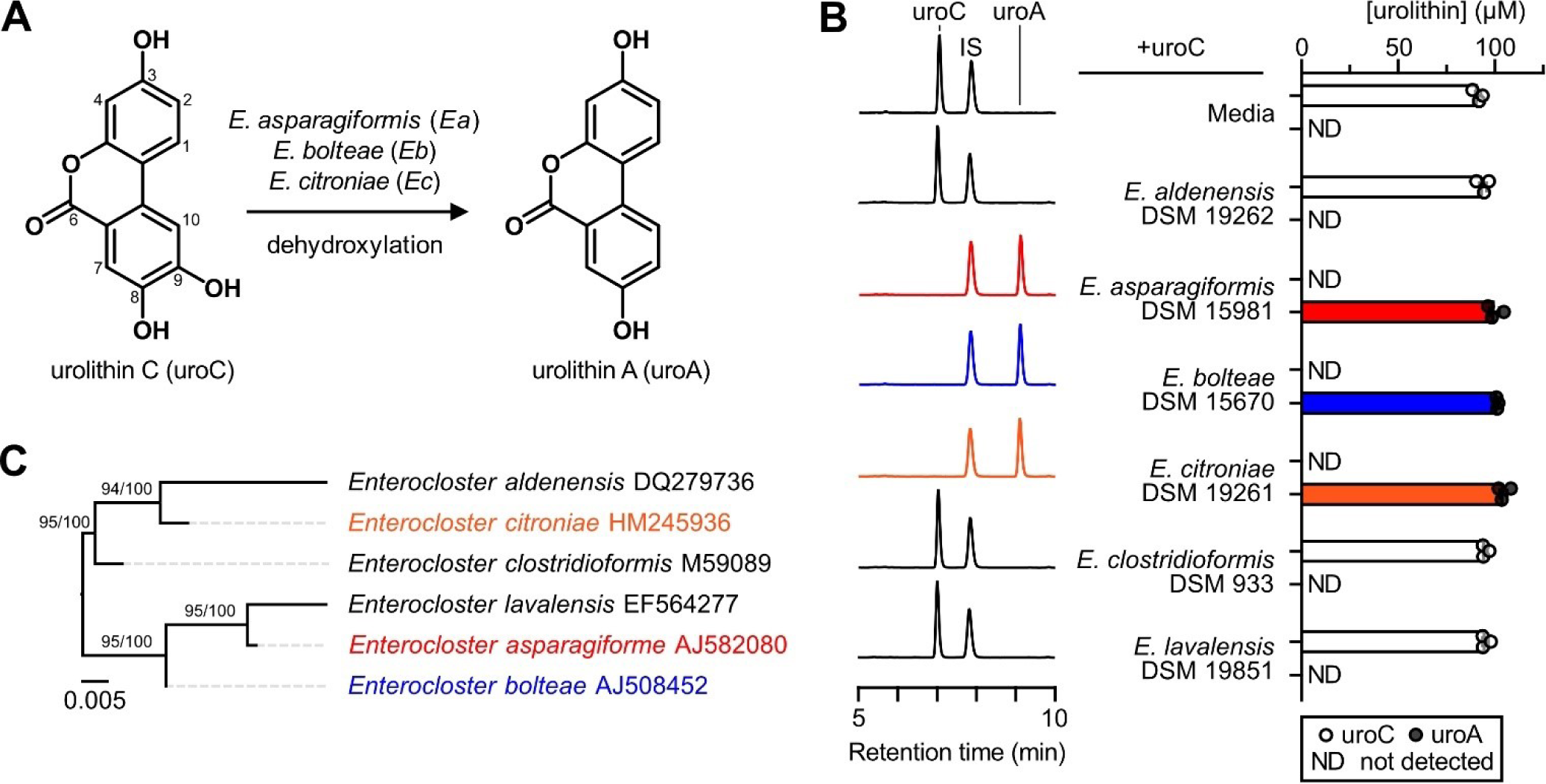
Urolithin C metabolism by *Enterocloster* spp. is not predicted by phylogeny. **A)** Reaction scheme of uroC dehydroxylation by gut resident *Enterocloster* spp. via unknown enzymes. **B)** LC-MS screen of *Enterocloster* spp. type strains for dehydroxylation activity. UroC (100 μM) was added to cultures (in mABB+H media) at the start of growth and urolithins were extracted after 24 h anaerobic incubation, then analyzed by LC-MS. Left: Representative chromatograms (λ = 305 nm) for each experimental group (from one representative biological replicate). The same scale was used for each chromatogram. Right: Quantification of urolithin peak areas relative to a salicylic acid internal standard (IS) (n = 3 biological replicates). Data are represented as mean ± SEM. **C)** Phylogenetic tree of tested *Enterocloster* spp. type strain 16S rRNA sequences constructed using the Genome-to-Genome Distance Calculator (GGDC) Phylogeny Server ^40^. Maximum likelihood (ML) tree inferred under the GTR+GAMMA model and rooted by midpoint-rooting. The branches are scaled in terms of the expected number of substitutions per site. The numbers above the branches are support values when larger than 60% from ML (left) and maximum parsimony (right) bootstrapping. The GenBank accession numbers are provided to the right of each taxon. Source data and statistical details are provided as a Source data file.

Here, we use a multi-omics enzyme identification framework to uncover uroC dehydroxylase (*ucd*) genes and enzymes in *Enterocloster* spp. and their relative, *Lachnoclostridium pacaense*. We find that the UcdCFO enzyme complex specifically dehydroxylates 9-hydroxy urolithins and that both metabolizing species and *ucd* genes are prevalent and actively transcribed in human feces during *ex vivo* metabolism. We further demonstrate that *Enterocloster* spp. growth is delayed by uroC and that dehydroxylation may be a mechanism to inactivate its iron-binding catechol moiety. Our study sheds light on the genetic and chemical basis underlying the complex reciprocal interactions between urolithins and the gut microbiota.

## Results

### A subset of *Enterocloster* species converts urolithin C to urolithin A *in vitro*

Members of the *Enterocloster* spp. have previously been shown to dehydroxylate uroC *in vitro* ^38^ and *in vivo* ^39^ under anaerobic conditions (Fig. 1A, full metabolic pathway in Supplementary Fig. 1). To determine the prevalence of uroC metabolism within this genus, we incubated all available *Enterocloster* spp. type strains (Methods) with uroC and quantified urolithin concentrations by liquid chromatography-mass spectrometry (LC-MS). Of the tested bacteria, only *E. asparagiformis*, *E. bolteae*, and *E. citroniae* dehydroxylated uroC to produce uroA (Fig. 1B). Interestingly, uroC metabolism was not predicted by phylogeny, as uroC-metabolizing species did not cluster based on 16S rRNA genes, genomes, or proteomes (Fig. 1C, Supplementary Fig. 2A,B, respectively), suggesting gain or loss of metabolic gene clusters throughout the evolution of *Enterocloster* spp. Based on these results, we chose to perform more in-depth analysis on *E. asparagiformis* and *E. bolteae* to identify the metabolic gene clusters involved in uroC dehydroxylation.

### A putative urolithin C dehydroxylase metabolic gene cluster is upregulated upon urolithin C treatment

To understand when uroC metabolism machinery was being expressed, we first sought to characterize the kinetics of uroC dehydroxylation in rich media (mABB+H). Therefore, a simultaneous growth and metabolism experiment was designed, whereby uroC was spiked into *E. asparagiformis* and *E. boltea*e cultures during the exponential phase of growth and metabolites were measured by LC-MS (Fig. 2A). Treatment with uroC during the exponential phase did not affect growth of either bacterium compared to the DMSO control (Fig. 2B). In both bacteria, quantitative conversion of uroC to uroA occurred within 4 h post-spike-in (Fig. 2C,D), demonstrating that metabolism in rich media is fast and robust.

**Figure 2.**
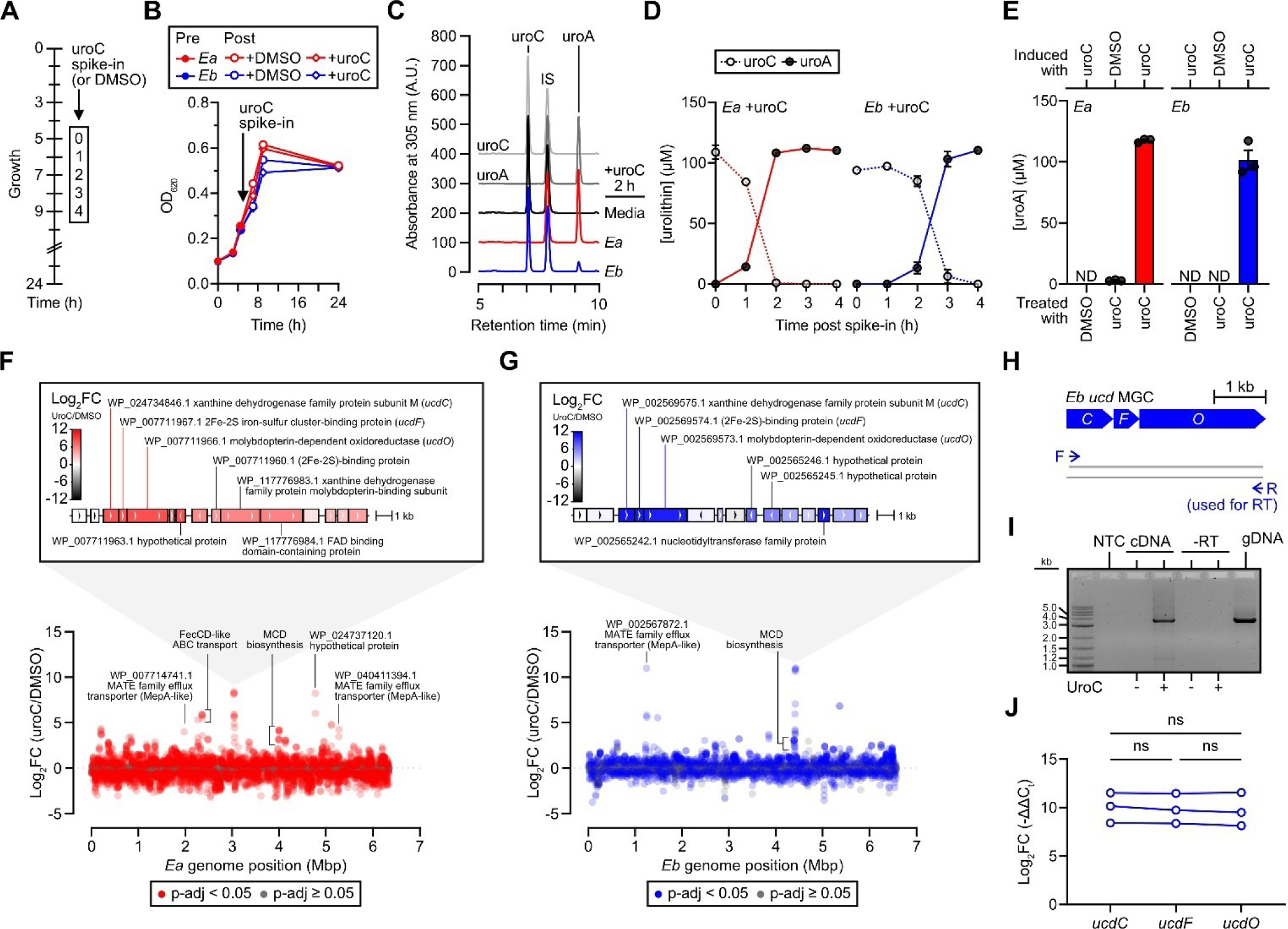
Urolithin C treatment upregulates a putative dehydroxylase operon. **A)** Experimental design of uroC (100 μM) spike-in experiments during the exponential phase of growth. For each biological replicate in this design, growth (B), metabolism (C,D), and RNA-seq (F-G) results are matched. **B)** Growth curve (optical density (OD) at 620 nm) of DMSO or uroC-spiked *E. asparagiformis* (*Ea*) and *E. bolteae* (*Eb*) type strain cultures according to the design in (A). 200 μL of culture were sampled at each timepoint and OD_620_ was measured in a 96-well plate (n = 4 biological replicates). The same sampled culture was then frozen and extracted for analysis by LC-MS (C,D). **C)** Representative chromatograms (λ = 305 nm) of cultures sampled 2 h post-spike-in (from one representative biological replicate). The same scale was used for each chromatogram. **D)** Quantification of urolithin concentrations from peak areas relative to a salicylic acid internal standard (IS) over 4 h in uroC-spiked *Ea* and *Eb* type strain cultures (n = 4 biological replicates). **E)** Quantification of urolithin A concentrations in DMSO- or uroC-treated *Ea* and *Eb* cell suspensions. Cell suspensions were prepared from *Ea* and *Eb* cells grown with either DMSO or 50 μM uroC. The cells were washed and resuspended in PBS to halt the production of new enzymes, then treated with DMSO or 100 μM uroC (n = 3 biological replicates). **F,G)** Manhattan plots of genes altered by uroC treatment in *Ea* (F) and *Eb* (G) based on DESeq2 analysis (n = 4 biological replicates). Data points are colored according to their adjusted p-value (based on the Benjamini-Hochberg-corrected Wald statistic). Grey, p-adj ≥ 0.05. Red or blue, p-adj < 0.05 for *Ea* and *Eb*, respectively. The genomic organization around the differentially expressed genes (generated from the NCBI Sequence Viewer) is depicted above Manhattan plots, which show the most highly and differentially expressed genes by RNA-seq. Genes are colored according to their log_2_FC values. NCBI accessions for select proteins encoded by highlighted genes are provided. **H)** Primer design for RT-PCR (I) experiment targeting the *Eb ucd* gene cluster. The same reverse primer was used for both the reverse transcription step and the subsequent PCR reaction. **I)** 1% agarose gel image of RT-PCR amplicons using primers (H) that span the full-length *Eb ucd* gene cluster (from one biological replicate). NTC, no template control. **J)** RT-qPCR expression of each gene in the *Eb ucd* operon. Growing *Eb* cultures were treated with DMSO or uroC (100 μM) for 2 h before RNA isolation and reverse transcription (n = 3 biological replicates). Gene expression profiles of each target gene in the *Eb ucd* gene cluster displayed as log_2_FC (equivalent to - ΔΔCt, where ΔΔC_t_ = ΔC_t uroC_ - ΔC_t DMSO_) with lines connecting paired biological replicates; repeated-measures one-way ANOVA with Tukey’s multiple comparisons test; ns, not significant. Data are represented as mean ± SEM (behind symbols) in (B,D,E). FC, fold change (uroC/DMSO); Source data and statistical details are provided as a Source data file.

We next sought to determine whether uroC metabolism is inducible or constitutive. To test for inducibility, both bacteria were treated with DMSO or uroC during exponential growth, then washed and resuspended in PBS, yielding cell suspensions unable to synthesize new proteins. Metabolism of uroC to uroA was inducible as only cells originating from bacteria grown with uroC were capable of uroA production (Fig. 2E). Consequently, we performed RNA-sequencing to compare gene expression in DMSO and uroC-treated cultures of *E. asparagiformis* and *E. bolteae*. Since uroA was detected in both bacterial cultures as soon as 2 h post spike-in (Fig. 2C), this timepoint was selected to isolate mRNA.

RNA sequencing of uroC-induced cultures revealed a distinct gene cluster, which we term uroC dehydroxylase (*ucd*), that was highly and differentially expressed (log_2_FC > 8) in both *E. asparagiformis* (Fig. 2F) and *E. bolteae* (Fig. 2G). In both bacteria, these clusters contained adjacent genes that were expressed to similar log_2_FC values: a xanthine dehydrogenase family protein subunit M, a (2Fe-2S)-binding protein, and a molybdopterin-dependent oxidoreductase (Fig. 2F,G). These genes will hereafter be referred to as *ucdC* (for coenzyme), *ucdF* (for ferredoxin), and *ucdO* (for oxidoreductase), respectively. Interestingly, we also observed an upregulation of genes involved in efflux (MepA-like multidrug and toxin extrusion (MATE) transporters) and iron transport (FecCD-like) (Fig. 2F,G), suggesting a link between uroC metabolism, iron uptake, and efflux.

### The *ucd* metabolic gene cluster is organized in an operon

We next sought to characterize the *ucd* metabolic gene cluster in *E. bolteae* since this bacterium is considered a core species of the gut microbiome ^8^. Based on the proximity, sense, and expression levels of each of the three genes by RNA-seq (Fig. 2G), we hypothesized that all three genes in the cluster were organized in an operon. We designed a gene-specific RT- PCR assay that would enable the detection of full-length polycistronic *ucdCFO* genes using cDNA from DMSO- or uroC-treated *E. bolteae* as a template (Fig. 2H). An amplicon of the expected size (∼3.6 kb) was detected only in cDNA derived from uroC-treated *E. bolteae*, validating the inducibility of these genes (Fig. 2I). Long-read sequencing of the obtained amplicon yielded a sequence corresponding to the *E. bolteae ucdCFO* metabolic gene cluster with 100% identity (Supplementary Sequence 1). Using an independent set of *E. bolteae* cultures, we then performed RT-qPCR on DMSO- or uroC-treated *E. bolteae* with all three genes in the putative operon as targets. Similar to our RNA-seq results, all three genes were highly induced (mean log_2_FC ≥ 9.7 for all three *ucd* genes) relative to DMSO controls and were expressed at the same level (Fig. 2J). These results indicate that the *ucdCFO* genes are transcribed as a single polycistronic mRNA and therefore form a uroC-inducible operon.

### The *ucd* operon is induced by 9-hydroxy urolithins

Next, we aimed to determine the substrate scope of the *ucd* operon. Multiple urolithins possess pyrogallol, catechol, and phenol structural motifs that are dehydroxylated at various positions by gut bacteria (Supplementary Fig. 1). Interestingly, *E. bolteae* only metabolizes the 9-position hydroxyl group of urolithins (Supplementary Fig. 3A-D) and does not require adjacent hydroxyl groups since isouroA is dehydroxylated to uroB (Supplementary Fig. 3D) ^38^. Since dehydroxylation of urolithins in *E. bolteae* is position specific, we hypothesized that the *ucd* operon would be induced by other 9-hydroxy urolithins (uroM6 and isouroA). Therefore, we performed RT-qPCR on DMSO-, uroM6-, uroC-, or isouroA-treated *E. bolteae* cultures using the *ucdO* gene as a target. Each urolithin significantly induced the expression of the *ucd* operon to a similar extent (Supplementary Fig. 3E-F). In addition, *E. bolteae* cell suspensions induced with uroC were capable of dehydroxylating uroM6 (Supplementary Fig. 3G) and isouroA (Supplementary Fig. 3H), indicating that the same proteins induced by uroC can metabolize structurally similar 9-hydroxy urolithins. Thus, it is likely that the same metabolic enzymes, encoded by the *E. bolteae ucd* operon, are acting on 9-hydroxy urolithins.

### Presence of *ucd* operon homologs in genomes predicts urolithin C metabolism by gut bacteria

We wondered whether novel metabolizers of uroC could be discovered based on nucleotide sequence homology to the *ucd* operon. Homology searches using the *E. bolteae ucd* operon sequence confirmed that only uroC-metabolizing *Enterocloster* spp. (*E. asparagiformis*, *E. bolteae*, and *E. citroniae*) possessed homologs of the *ucdCFO* genes with a similar organization (Supplementary Fig. 4). In addition, the gut bacterium *Lachnoclostridium pacaense* ^41^ was identified as another hit (Supplementary Fig. 4). The type strain of this bacterium (CCUG 71489T = Marseille-P3100) was closely related to *Enterocloster* spp. Based on 16S rRNA, whole genome, and whole proteome phylogenies (Fig. 3A, Supplementary Fig. 2A,B, respectively). *L. pacaense* possessed genomic sequences with high homology (86.5% nucleotide identity) and identical functional annotations to the *E. bolteae ucd* operon sequence (Fig. 3B). When incubated with uroC, *L. pacaense* CCUG 71489T quantitatively produced uroA (Fig. 3C,D). We searched for homologs of the *E. bolteae ucd* in the genomes of urolithin- and catechol-metabolizing bacteria belonging to the *Eggerthellaceae* but could not identify any hits. Notably, *Eggerthellaceae* lack 9-hydroxy urolithin dehydroxylase activity ^35,37^, which correlates with an absence of *ucd*-like operons in their genomes (Fig. 3B). These comparative genomics data indicate that the presence of a *ucd* operon in genomes predicts uroC metabolism by gut bacteria.

**Figure 3.**
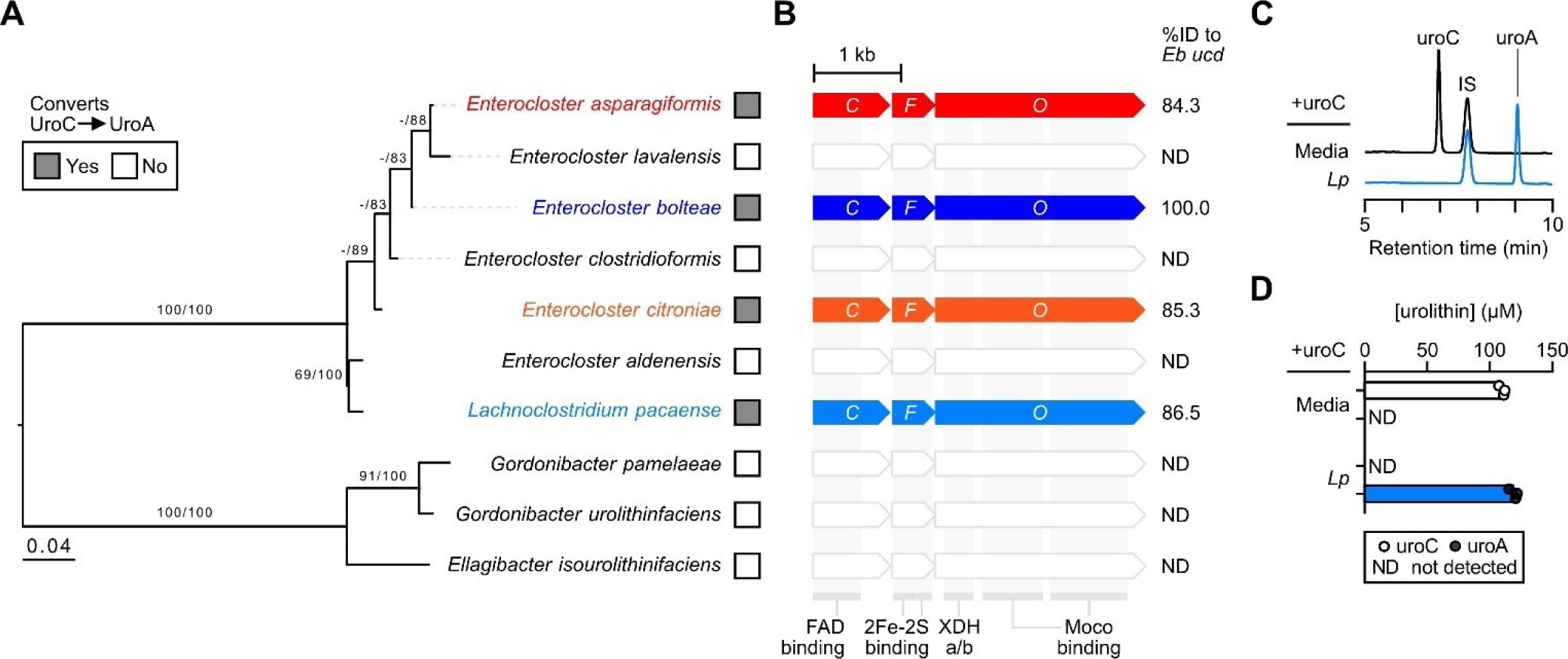
Urolithin C metabolism correlates with *ucd* operon prevalence in gut bacteria. **A)** Phylogenetic tree of *Enterocloster* spp., *Lachnoclostridium pacaense* (*Lp*), and catechol- metabolizing *Eggerthellaceae* type strain 16S rRNA sequences constructed using the Genome-to-Genome Distance Calculator (GGDC) Phylogeny Server ^40^. Maximum likelihood (ML) tree inferred under the GTR+GAMMA model and rooted by midpoint-rooting. The branches are scaled in terms of the expected number of substitutions per site. The numbers above the branches are support values when larger than 60% from ML (left) and maximum parsimony (right) bootstrapping. Bacteria that convert uroC to uroA are labeled with grey squares. **B)** NBCI Multiple Sequence Aligner viewer hits for BLASTn searches using the *E. bolteae* DSM 15670 *ucd* operon nucleotide sequence as a query against the NCBI refseq_genomes database. Only hits with ≥ 90 % query coverage and species-level taxonomic resolution are displayed with % identity to the query sequence. Domain annotations for each gene are denoted below according to InterPro annotations for corresponding proteins. ND, Not detected; Moco, Molybdenum cofactor. **C,D)** *In vitro* metabolism of uroC by *Lp*. UroC (100 μM) was added to cultures (in mABB+H media) at the start of growth and urolithins were extracted after 24 h anaerobic incubation, then analyzed by LC-MS. **C)** Representative chromatograms (λ = 305 nm) for each experimental group (from one representative biological replicate). The same scale was used for each chromatogram. **D)** Quantification of urolithin peak areas relative to a salicylic acid internal standard (IS) (n = 3 biological replicates). Data are represented as mean ± SEM. Source data and statistical details are provided as a Source data file.

### A molybdopterin cofactor biosynthetic gene cluster is upregulated upon urolithin C treatment

In addition to the three genes in the *ucd* operon, we observed a significant increase (log2FC ≥ 2.6) in 9 molybdopterin cytosine dinucleotide (MCD) biosynthesis genes upon uroC treatment (Fig. 2F,G, Supplementary Fig. 5A). These 9 genes, which recapitulate the function of 10 genes in *E. coli* ^42^, are involved in molybdenum cofactor biosynthesis (*moaAC*, *mogA*, *moeA*), molybdate ion transport (*modABCE*), cytosine addition to the molybdenum cofactor (*mocA*), and MCD cofactor insertion into the active site (*xdhC*) (Supplementary Fig. 5B) ^42^. All 9 genes cluster in the genomes of *E. asparagiformis* and *E. bolteae* and are organized into 2 adjacent operons (Supplementary Fig. 5C) that are induced upon uroC treatment. Based on sequence homology to *E. coli* oxidoreductases and MCD biosynthetic machinery, proteins encoded by the *ucd* operon belong to the xanthine dehydrogenase family ^43^. These findings imply that uroC dehydroxylation is MCD-dependent, which differs from the bis-molybdopterin guanine dinucleotide requirement of catechol dehydroxylases in *Eggerthellaceae* ^10,14^.

### The UcdCFO complex enables anaerobic electron transport from NADH to uroC

Since oxidoreductases utilise a variety of cofactors and coenzymes for catalytic activity, we sought to determine the redox coenzymes and conditions necessary for uroC dehydroxylation. Therefore, we performed metabolism assays using crude lysates from uroC-induced *E. bolteae*. As crude lysates alone did not metabolize uroC, various redox coenzymes (NADPH, NADH, and FAD) were added to lysates to promote uroC dehydroxylation (Fig. 4A). Only NADH-treated lysates yielded quantitative dehydroxylation of uroC to uroA compared to the no cofactor control (Fig. 4A). Interestingly, the addition of free FAD partially inhibited uroC dehydroxylation in NADH-treated lysates (Fig. 4A), likely by decreasing the free NADH pool. NADPH, which differs from NADH by a phosphate group on the 2’-OH group of the adenosine moiety, was unable to promote uroC dehydroxylation, indicating some specificity in the redox cofactors necessary for dehydroxylation. Aerobic incubation of crude lysates supplemented with NADH completely inhibited uroC dehydroxylation (Fig. 4B), indicating that the active enzyme complex requires a strictly anaerobic environment for dehydroxylation, as has been demonstrated for various metalloenzymes ^44^.

**Figure 4.**
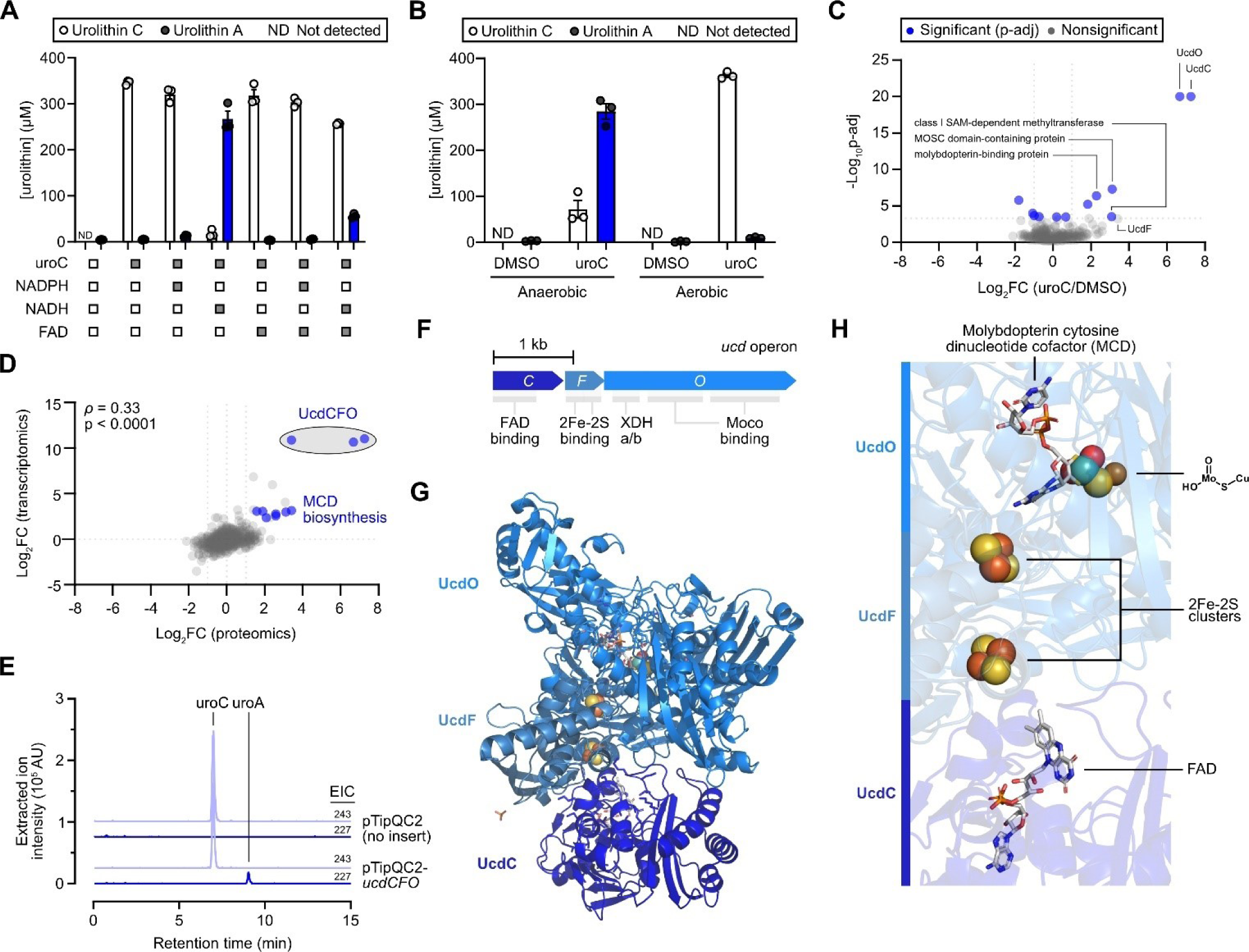
The UcdCFO complex enables anaerobic electron transport from NADH to uroC. **A)** Quantification of urolithin peak concentrations in crude uroC-induced *Eb* lysates re-treated with DMSO or uroC (350 μM) and various coenzymes (n = 3 biological replicates). NADPH, nicotinamide adenine dinucleotide phosphate; NADH, nicotinamide adenine dinucleotide; FAD, flavin adenine dinucleotide. **B)** Quantification of urolithin peak concentrations in crude uroC-induced *Eb* lysates re-treated with DMSO or uroC (350 μM) and NADH in anaerobic or aerobic environments (n = 3 biological replicates). Data are represented as mean ± SEM for (A,B). **C)** Volcano plot of untargeted proteomics analysis on DMSO or uroC-treated *Eb* (n = 3 biological replicates). Data points are colored according to their significance (Fisher’s exact test with Benjamini-Hochberg correction for multiple comparisons). Grey, p-adj ≥ cutoff p-value (0.00048). Blue, p-adj < cutoff p-value (0.00048). **D)** Scatter plot showing the correlation between gene and protein expression (log_2_FC values) induced in uroC-treated *Eb* using the datasets in Fig. 2G and Fig. 5C, respectively. The non-parametric Spearman rank correlation test was used for statistical analysis. **E)** LC-MS extracted ion chromatograms (EIC) of uroC ([M-H]^-^ = 243) and uroA ([M-H]^-^ = 227) from a representative anaerobic uroC dehydroxylation assay using crude lysates of *R. erythropolis* harboring either pTipQC2 (no insert) or pTipQC2-*ucdCFO* plasmids. **F)** Domains of genes in the *ucd* operon based on InterPro annotations. **G)** Quaternary structure prediction of the proteins encoded by the *Eb ucd* operon. AlphaFold2 structures for each protein were superposed onto the X-ray crystal structure of PDB 1ZXI (carbon monoxide dehydrogenase from *Afipia carboxidovorans* OM5). **H)** Small molecule ligands from PDB 1ZXI in the superposed UcdCFO model form a complete electron transport chain from FAD to two 2Fe-2S clusters to a molybdopterin cytosine dinucleotide cofactor, which can then reduce uroC (terminal electron acceptor). Source data and statistical details are provided as a Source data file.

To confirm that *ucd* operon-encoded proteins were expressed in *E. bolteae* crude lysates, we performed untargeted proteomics and compared protein expression upon DMSO or uroC treatment. Indeed, all 3 proteins encoded by the *ucd* operon (UcdC, UcdF, and UcdO) were the most differentially expressed proteins in the uroC treatment group (Fig. 4C). In addition, proteins involved in MCD biosynthesis were also strongly increased upon uroC treatment (Fig. 4C,D, Supplementary Fig. 5A), pointing to the coordination between MCD biosynthesis and active UcdCFO oxidoreductase assembly. These multi-omics datasets implicate all three *ucdCFO* genes and MCD biosynthesis genes in the metabolism of uroC to uroA, as demonstrated by the strong positive correlation between transcript and protein differential expression (Fig. 4D).

To validate the function of the *E. bolteae ucd* operon, we attempted heterologous expression of *E. bolteae* UcdCFO in *E. coli*; however, all expression and activity assays were unsuccessful despite the inclusion of *mocA* and *xdhC* genes involved in MCD maturation in our expression plasmids. This lack of activity likely resulted from the choice of heterologous host and from the complex assembly of active molybdoenzymes ^43^. We therefore attempted to express UcdCFO in the phenol-degrading soil bacterium *Rhodococcus erythropolis* using a thiostrepton-inducible expression system ^45^ (pTipQC2-*ucdCFO*, Supplementary Fig. 6A,B), previously used to express the anaerobic *E. lenta* Cgr2 protein ^44^. Despite the poor yield of soluble protein (Supplementary Fig. 6C,D), we were able to observe uroM6 and uroC dehydroxylation at the 9-position in crude lysates of *R. erythropolis* transformed with pTipQC2-*ucdCFO*, but not in the no insert control (pTipQC2) (Fig. 4E, Supplementary Fig. 6E-G), thus confirming that the *ucd* operon confers 9-hydroxy urolithin dehydroxylase activity.

To gain an understanding of the structural organization of proteins encoded by the *ucd* operon, we performed modeling using AlphaFold2 ^46,47^. Structures of each protein encoded by the *E. bolteae ucd* operon (Fig. 4F) were superposed onto published X-ray crystal structures of xanthine dehydrogenase family enzymes with similar folds ^48^, yet from different taxonomic domains: *Afipia carboxidovorans* carbon monoxide dehydrogenase ^49^ and *Bos taurus* xanthine dehydrogenase ^50^. The 3 proteins encoded by the *ucd* operon formed subunits in an oxidoreductase complex with a similar quaternary structure to the published crystal structures (Fig. 4G, Supplementary Fig. 7A,B). The predicted quaternary structure of the UcdCFO enzyme complex supported a complete electron transport chain whereby electrons would flow from reduced FAD to two 2Fe-2S clusters, then to the MCD cofactor, and finally to uroC as the terminal electron acceptor (Fig. 4H, Supplementary Fig. 7C,D). This model supports our findings in crude lysates whereby NADH serves as an electron donor to reduce UcdC-bound FAD (Fig. 4A). Using homology modeling, we further identified the putative uroC binding site in UcdO, which overlaps with the salicylic acid ligand in the *Bos taurus* xanthine dehydrogenase structure (Supplementary Fig. 7E). This putative uroC binding site contains multiple tyrosine (Y375, Y538, Y624, Y632), tryptophan (W345), and phenylalanine (F458, F464) residues that could form π-π stacking interactions with uroC (Supplementary Fig. 7F), orienting it towards the molybdenum cofactor.

### Disruption of the urolithin C catechol moiety rescues growth delay in iron-limited media

To gain an understanding of why *Enterocloster* spp. and *L. pacaense* metabolize 9-hydroxy urolithins, we performed growth experiments in different media conditions. When uroC was added prior to growth in rich medium containing hemin (mABB+H), a concentration-dependent increase in lag time was observed for all uroC-metabolizing bacteria (Fig. 5A,B). As catechols are common structural motifs in iron-binding siderophores ^51^, we hypothesized that uroC was delaying growth by altering iron availability in the growth medium *via* its catechol moiety. Incubation of *Enterocloster* spp. and *L. pacaense* in medium lacking added iron (mABB) exacerbated the growth delay by uroC (Fig. 5C, Supplementary Fig. 8A,B); however, this growth delay was partially rescued upon supplementation of different iron sources (hemin, Fe(II)SO_4_, or Fe(III) pyrophosphate) (Fig. 5C, Supplementary Fig. 8C). To validate that iron chelation could extend the lag time of *Enterocloster* spp. and *L. pacaense*, we incubated all four uroC-metabolizers with 2,2’-bipyridyl (biP) in mABB media. As observed with uroC, biP delayed the growth of all tested bacteria, but supplementation of Fe(II)SO_4_, or Fe(III) pyrophosphate could partially rescue growth delay (Supplementary Fig. 8C). Interestingly, uroA, which lacks a catechol moiety, did not impact the growth of the tested bacteria in either mABB or mABB+H media compared to uroC (Fig. 5C, Supplementary Fig. 8A,B). To confirm that the catechol moiety of uroC was responsible for delaying growth, we synthesized a methylated analogue of uroC (8,9-di-O-methyl-uroC, Fig. 5D, Supplementary Fig. 9A-F), which is unable to bind iron ^52^, and tested its effect on growth in mABB. Like uroA, 8,9-di-O-methyl-uroC did not delay growth of uroC-metabolizing bacteria (Fig. 5E,F). These data demonstrate that both the catechol moiety of uroC and iron availability are essential to uroC-mediated lag phase extension.

**Figure 5.**
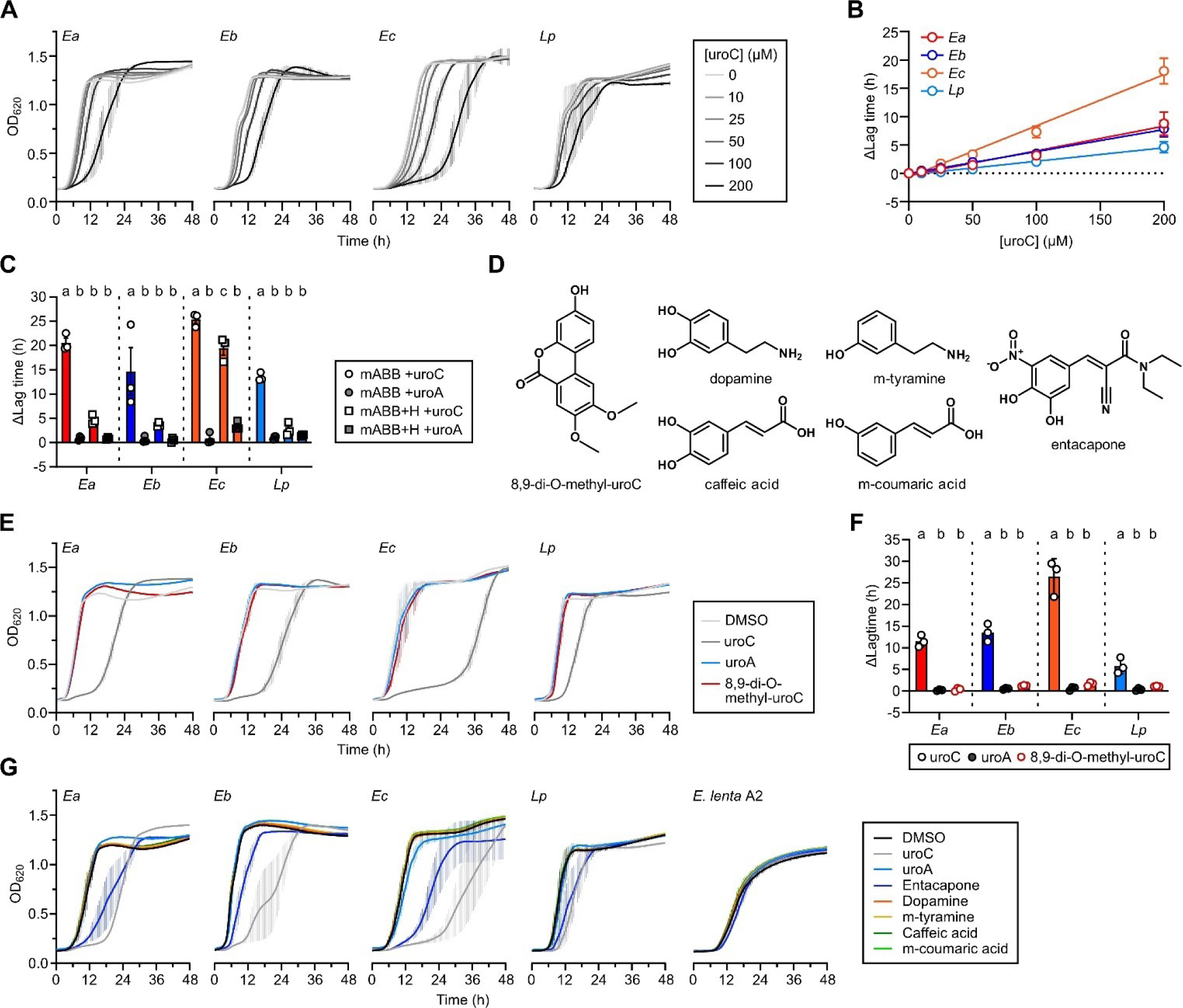
The catechol moiety of uroC delays Enterocloster spp. growth in an iron-dependent manner. **A)** Growth curves (optical density (OD) at 620 nm) of uroC-metabolizing *Enterocloster* spp. and *L. pacaense* treated with increasing concentrations of uroC in rich mABB+H media (7.7 μM hemin) (n = 3 biological replicates). Data are represented as mean ± SEM. **B)** Quantification of the difference in lag time compared to the DMSO control for growth curves in (A). Data are represented as mean ± SEM; lines were fitted using simple linear regression. **C)** Quantification of the difference in lag time of *Enterocloster* spp. and *L. pacaense* grown in mABB (no added iron) or mABB+H (7.7 μM hemin) compared to respective DMSO controls for growth curves in Supplementary Fig. 8A,B. Data are represented as mean ± SEM; repeated measures two-way ANOVA (matching by biological replicate) with Tukey’s multiple comparisons test. Significant differences between treatments for individual bacteria are denoted by a different lowercase letter above each plot. **D)** Structures of tested catechols and their derivatives. **E)** Growth curves (OD at 620 nm) of uroC-metabolizing *Enterocloster* spp. and *L. pacaense* grown in mABB media and treated with DMSO (vehicle), 100 μM of uroC, uroA, or 8,9-di-O-methyl-uroC (n = 3 biological replicates). Data are represented as mean ± SEM. **F)** Quantification of the difference in lag time compared to the DMSO control for growth curves in (E). Data are represented as mean ± SEM; repeated measures two-way ANOVA (matching by biological replicate) with Tukey’s multiple comparisons test. Significant differences between treatments for individual bacteria are denoted by a different lowercase letter above each plot. **G)** Growth curves (OD at 620 nm) of uroC-metabolizing *Enterocloster* spp., *L. pacaense*, and *E. lenta* A2 grown in mABB media and treated with DMSO (vehicle), 100 μM of uroC, uroA, entacapone, dopamine, m-tyramine, caffeic acid or m-coumaric acid. Data are represented as mean ± SEM.

Dehydroxylation of catechols by gut bacteria has been observed for diverse classes of compounds like neurotransmitters, therapeutic drugs, and diet-derived polyphenols ^14^. Although catechol dehydroxylation can promote growth in some species ^14^, we hypothesized that dehydroxylation could be a mechanism used by gut bacteria to inactivate catechol-containing compounds that affect their fitness. To determine whether diverse catechols can delay growth, uroC-metabolizing bacteria and the dopamine-metabolizing *Eggerthella lenta* A2 were incubated with catechol-containing compounds and their dehydroxylated counterparts: uroC (uroA), entacapone, dopamine (m-tyramine), caffeic acid (m-coumaric acid) (Fig. 5D). Surprisingly, neither dopamine nor caffeic acid (and their dehydroxylated counterparts) delayed the growth of the tested bacteria (Fig. 5G). On the other hand, both uroC and the nitrocatechol-containing Parkinson’s drug entacapone delayed the growth of *Enterocloster spp*. and *L. pacaense* but did not affect *E. lenta* A2 (Fig. 5G) ^8^. Thus, catechol-containing compounds show differential effects on the growth of gut bacteria, depending on their structure. These results prompted us to investigate the effect of uroC on a more diverse panel of gut bacteria including *E. aldenensis, E. clostridioformis*, and *E. lavalensis*, which do not metabolize uroC, along with *Gordonibacter* spp., which produce uroC from dietary ellagic acid ^35^. Treatment with uroC delayed the growth of *E. aldenensis*, *E. clostridioformis,* and *E. lavalensis* to varying extents (Supplementary Fig. 10A); however, there was no difference in growth between the DMSO-, uroC-, and uroA-treated cultures of *Enterococcus faecium* and *Gordonibacter* spp. (Supplementary Fig. 10B,C). Thus, all *Enterocloster* spp. tested showed sensitivity to uroC-mediated lag time extension, while other bacteria were insensitive to its effects on growth.

### UroC-metabolizing species and *ucd* genes are prevalent and correlate with uroC metabolism in human fecal samples

We next wondered whether uroC-metabolizing *Enterocloster* spp. and their *ucd* operons were prevalent and active in human fecal samples. We first utilized uniformly processed metagenomic data from the curatedMetagenomicData R package ^53^. After filtering for fecal samples (86 studies, n = 21,030 subjects), we counted the prevalence of at least one uroC-metabolizing species and at least one *ucd* gene homolog (Methods). The prevalence of both features was variable across studies (Fig. 6A for studies with >200 participants, Supplementary Fig. 11A for all studies). Combining all studies, the prevalence of at least one uroC-metabolizing species and at least one *ucdCFO* gene homolog was 9,343/21,030 (44.9%) and 4,356/21,030 (20.7%), respectively. *E. bolteae* was the most prevalent and abundant uroC-metabolizing species detected in gut metagenomes (Supplementary Fig. 11B,C) and correlated strongly with ucd abundance (Supplementary Fig. 11D). These findings suggest that uroC-metabolizing *Enterocloster* spp. and *ucd* operon genes are prevalent in human fecal metagenomic samples and reflect the variable urolithin metabolism profiles (metabotypes) in the general population^25,54^.

**Figure 6.**
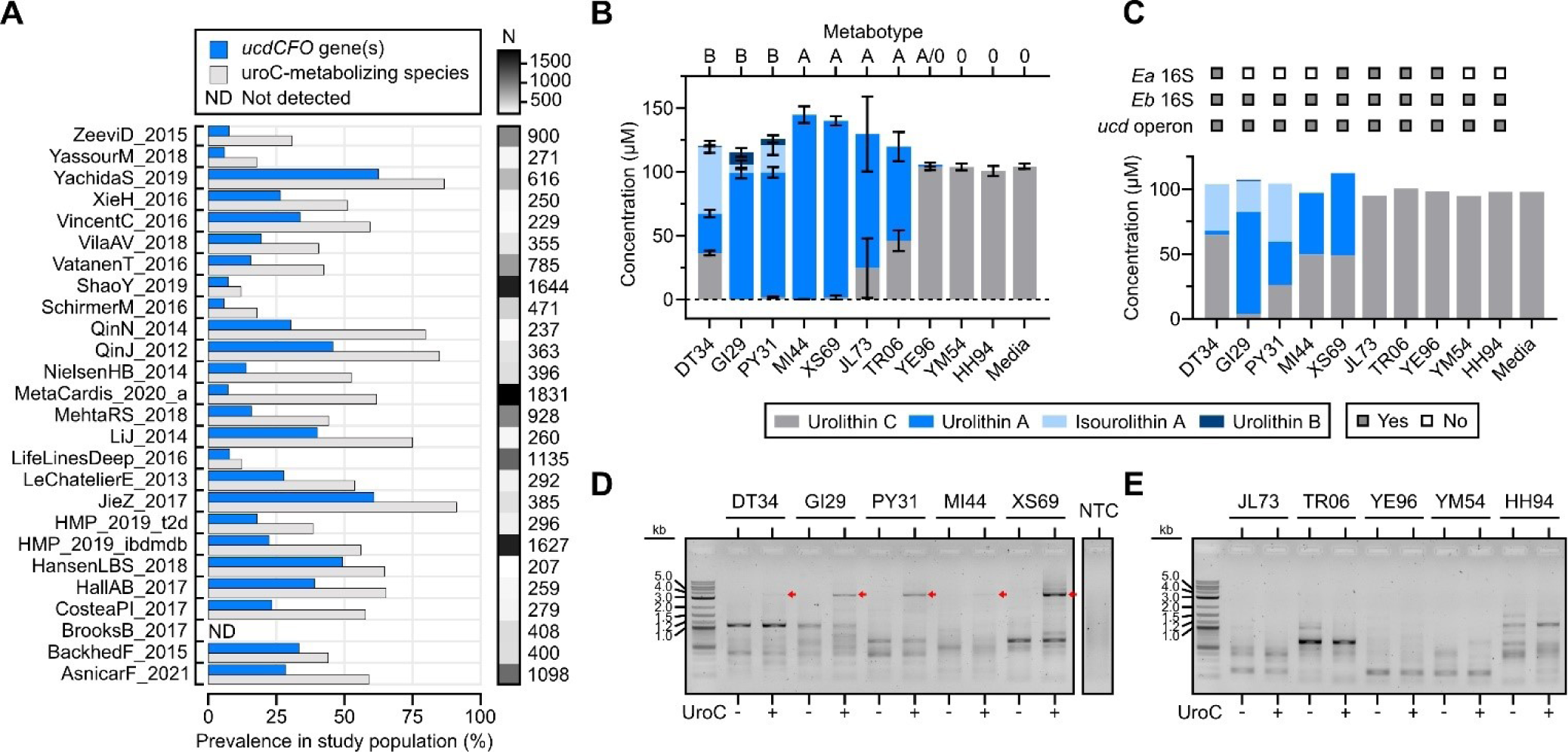
The *ucd* operon is prevalent in metagenomes and actively transcribed in urolithin C-metabolizing human fecal samples. **A)** Prevalence of *ucd* operon (at least one gene) and of a uroC-metabolizing species (at least one species) in fecal metagenomes from the CuratedMetagenomicData R package. Only studies with ≥200 participants are depicted. All 86 studies are available in Supplementary Fig. 11. **B)** Summary of urolithin concentrations in fecal slurries (n = 10 healthy donors) incubated with 100 μM uroC for 48 h. Data are represented as mean ± SEM (n = 3-6 experimental replicates). **C)** Summary of urolithin concentrations in fecal slurries (n = 10 healthy donors) incubated with 100 μM uroC for 48 h. Presence of uroC-metabolizing species and of the *ucd* operon is denoted above the graph if the bacterium or operon was detected in the uroC-treated fecal slurry (Supplementary Fig. 12B,F,G). Data are representative of 1 replicate where DNA and RNA was also extracted from fecal slurries. **D,E)** *ucd* gene-specific RT-PCR on fecal microbiota communities from 10 healthy donors using the primer set described in Fig. 2H. Samples are matched to the urolithin metabolism data in C) and 16S rRNA sequencing in Supplementary Fig. 12A-E. 1% agarose gel of amplicons derived from **D)** uroC-metabolizing and **E)** non-metabolizing fecal slurries. Bands corresponding to the *E. bolteae ucd* operon (∼3.6 kb) are labeled with red arrows. The no template control (NTC) is the same for (D,E). See Supplementary Fig. 12H,I for the no reverse transcriptase control PCR reactions on the same samples. Source data are provided as a Source data file.

Next, we performed *ex vivo* metabolism assays to determine whether *Enterocloster* spp. could metabolize uroC in the context of a complex community. Fecal slurries from 10 healthy individuals were first profiled according to their uroC metabotypes (Fig. 6B) ^24^. Individuals clustered into metabotypes A (only uroA produced), B (uroA and isouroA/uroB), and 0 (no terminal urolithin metabolites). While we observed metabotypes A and B in uroC-metabolizing fecal slurries, all slurries produced some amount of uroA from uroC. Stools JL73, TR06, and YE96 displayed variable metabolism patterns and did not metabolize uroC in some experiments, likely reflecting differences in activity between aliquots of feces (Fig. 6B). We then repeated metabolism assays using fecal slurries from all 10 healthy individuals and extracted urolithins, DNA, and RNA from each culture. In this experiment, only 5/10 fecal slurries metabolized uroC to uroA (Fig. 6C). We hypothesized that differences in metabotypes could be explained by microbial composition. Therefore, long-read V1-V9 16S rRNA sequencing was performed on fecal slurries. Both DMSO- and uroC-treated fecal slurries within individuals had similar microbial compositions and diversity metrics (Supplementary Fig. 12A-D) but showed differences in composition between individuals and metabolism status (Supplementary Fig. 12A,E). Surprisingly, all samples contained 16S rRNA sequences mapping to *E. bolteae*, and many non-metabolizing fecal slurries contained *E. asparagiformis* (Fig. 6C, Supplementary Fig. 12B). We then assayed genomic DNA from treated fecal slurries for the presence of the *ucd* operon by PCR and found that 10/10 individuals (19/20 conditions) yielded a detectable amplicon of the expected size (∼3.6 kb) (Supplementary Fig. 12F,G). These data indicate that the prevalence of uroC-metabolizing *Enterocloster* spp. 16S rRNA and *ucd* operon genes does not predict metabolism in fecal samples.

We then surmised that the *ucd* operon would be transcribed only in fecal slurries actively metabolizing uroC. Using a gene-specific reverse primer that binds to *ucdO* (Fig. 2H), the full-length *ucd* operon was reverse transcribed and amplified in RNA extracted from DMSO- and uroC-treated fecal slurries. An amplicon (∼3.6 kb) corresponding to the *ucd* operon was only detected in uroC-metabolizing fecal slurries (Fig. 6D) when treated with uroC and entirely absent from non-metabolizing slurries (Fig. 6E). This amplicon was absent in no reverse transcriptase controls, indicating no gDNA contamination (Supplementary Fig. 12H,I). These data demonstrate that *ucd* transcription correlates with uroC metabolism in complex fecal communities and that *E. bolteae* is keystone species involved in urolithin A production.

## Discussion

We identified genes and proteins that are essential for the metabolism of urolithins by gut resident *Lachnospiraceae* through a combination of transcriptomics, comparative genomics, and untargeted proteomics. Our study reveals a novel multi-subunit molybdoenzyme (urolithin C dehydroxylase, Ucd) that catalyzes the dehydroxylation of 9-hydroxy urolithins including uroM6, uroC, and isouroA. Importantly, prevalence analysis in published data and *ex vivo* transcriptomics established *E. bolteae* as a keystone urolithin-metabolizing member of the gut microbiota.

Catechol dehydroxylases are widespread in gut resident *Eggerthella lenta* and *Gordonibacter* spp. ^10,14,55^. These molybdoenzymes, which belong to the DMSO reductase superfamily ^56^, dehydroxylate substrates like catechol lignan (Cldh), dopamine (Dadh), DOPAC (Dodh), hydrocaffeic acid (Hcdh), and caffeic acid (Cadh), which can promote growth by using these substrates as alternative electron acceptors ^14^. A recent survey of reductases in gut bacteria established that most respiratory reductases contain N-terminal signal sequences and are translocated across the cytoplasmic membrane, while non-respiratory reductases, which lack signal sequences, remain in the cytoplasm ^57^. The UcdCFO enzyme complex we found in *Enterocloster* spp. differs from catechol dehydroxylases in *Eggerthellaceae* in important ways as it does not require a catechol structural motif for activity, belongs to the xanthine oxidase superfamily, and is composed of 3 subunits that each lack signal sequences. Based on the absence of signal sequences and the cytoplasmic localization of xanthine dehydrogenases, the *ucd* operon likely encodes for a non-respiratory reductase serving a different role than previously characterized catechol dehydroxylases ^57^.

In rich media conditions, uroC, but not uroA, extended the lag phase of growth in both uroC-metabolizing and non-metabolizing *Enterocloster* spp. This growth delay was not observed for other taxa, suggesting that *Enterocloster* spp. are especially sensitive to uroC-mediated iron chelation. In addition to the *ucd* operon, uroC-treated *E. bolteae* and *E. asparagiformis* upregulated gene clusters related to efflux (MepA-like MATE family efflux transporters) and iron/siderophore transport (FecCD-like ABC transporter). These responses are analogous to antimicrobial resistance mechanisms raised against entacapone and other non-antibiotic drugs ^8^. This suggests that non-respiratory uroC dehydroxylation could serve as an additional strategy that evolved in *E. asparagiformis*, *E. bolteae*, *E. citroniae*, and *L. pacaense* to overcome catechol-mediated iron chelation.

While uroA is the most common terminal metabolite following ellagitannin consumption in humans, its production varies widely ^36^. Interestingly, the ability of a fecal sample to produce uroA from uroC did not correlate with the presence of widespread uroC-metabolizing *Enterocloster* spp. or a *ucd* gene homologue, likely owing to poor viability or dead bacteria in fecal samples. However, the active transcription of the *ucd* operon correlated perfectly with metabotypes. These findings further emphasize the importance of functional assays such as transcriptomics and *ex vivo* metabolism to understand the metabolism of xenobiotics by the gut microbiota.

By identifying the genetic basis for metabolism of uroC, we found a novel metabolizing species that could not have been predicted based on phylogeny alone. Our data suggests that *ucd*-containing *Enterocloster* spp., and the closely related *L. pacaense*, are the main drivers of urolithin A production in the gut microbiota based on their prevalence in metagenomes and activity in fecal samples. However, we cannot conclude that they are solely responsible for this activity. Rare, strain-specific urolithin A production has been reported for *Bifidobacterium pseudocatenulatum* INIA P815 ^36^, *Streptococcus thermophilus* FUA329 ^58^, and *Enterococcus faecium* FUA027 ^58^, which may be a result of horizontal gene transfer since it is not shared by other members of the taxa. Thus, further enzyme discovery efforts are necessary to understand urolithin production in these bacteria.

In conclusion, our studies reveal the genetic and chemical basis for urolithin A production by gut bacteria and broaden our understanding of the molecular mechanisms underlying urolithin metabotypes in human populations. Since diet can modulate gut microbiota function and host health, elucidating the xenobiotic metabolism genes encoded by gut bacteria will be key to developing dietary interventions targeting the gut microbiota.

## Acknowledgements

This research was funded by the Canadian Institutes of Health Research (CIHR) grant PJT-437944 to B.C. and C.F.M. and a Weston Family Microbiome Initiative’s 2021 Transformational Research program to B.C. B.C. holds a tier II Canada Research Chair (CRC) in Therapeutic Chemistry. C.F.M. holds a tier II CRC in Gut Microbial Physiology and is a Azrieli Global Scholar in the Humans & Microbiome program. R.P. is supported by the CIHR Canada Graduate Scholarship-Doctoral (#493808 for 2023-2026) and by the Fonds de Recherche du Québec-Santé : Bourse de formation au doctorat (#316063 for 2022-2023). M.S. is supported by the CIHR Canada Graduate Scholarship to Honor Nelson Mandela (#DF2-187718 for 2023-2026), the Fonds de Recherche du Québec-Santé: Bourse de formation au doctorat (#311071 for 2022-2023).

RNA sample preparation and sequencing was performed by Génome Québec. Untargeted proteomics sample preparation and analysis was performed by the Research Institute of the McGill University Health Centre (RI-MUHC) Proteomics and Molecular Analysis Platform. Preliminary small-molecule MALDI-TOF mass spectrometry was performed by Mark Hancock at the McGill SPR-MS Facility. High resolution mass spectrometry was performed by the McGill Chemistry Characterization Facility. Molecular biology experiments were enabled by the McGill University Imaging and Molecular Biology Platform (IMBP). High-performance cloud computing was enabled by Calcul Québec (https://www.calculquebec.ca/) and the Digital Research Alliance of Canada (https://alliancecan.ca/). RNA-seq data analysis was performed using two instances of Galaxy: (https://usegalaxy.org/) and Compute Canada Genetics and Genomics Analysis Platform (GenAP) (https://www.genap.ca/).

## Contributions

**R.P.** designed the study, performed microbiology experiments, performed bioinformatics analyses, analyzed data, created figures, and wrote the initial manuscript with B.C. **S.M.** performed experiments on microbial communities, analyzed data, and created figures. **M.S.** performed Nanopore sequencing and processed raw sequencing reads. **L.S.** performed microbiology experiments. **L.D.** consulted on experimental design and methodology. **C.M.** supervised research, obtained ethical approval for the use of human fecal samples, and obtained research funding. **B.C.** designed the study, supervised the research, obtained research funding, and wrote the initial manuscript with R.P. All authors reviewed and edited the manuscript.

## Ethics declarations

The authors declare no competing interests.

## Online Methods

### Resources table

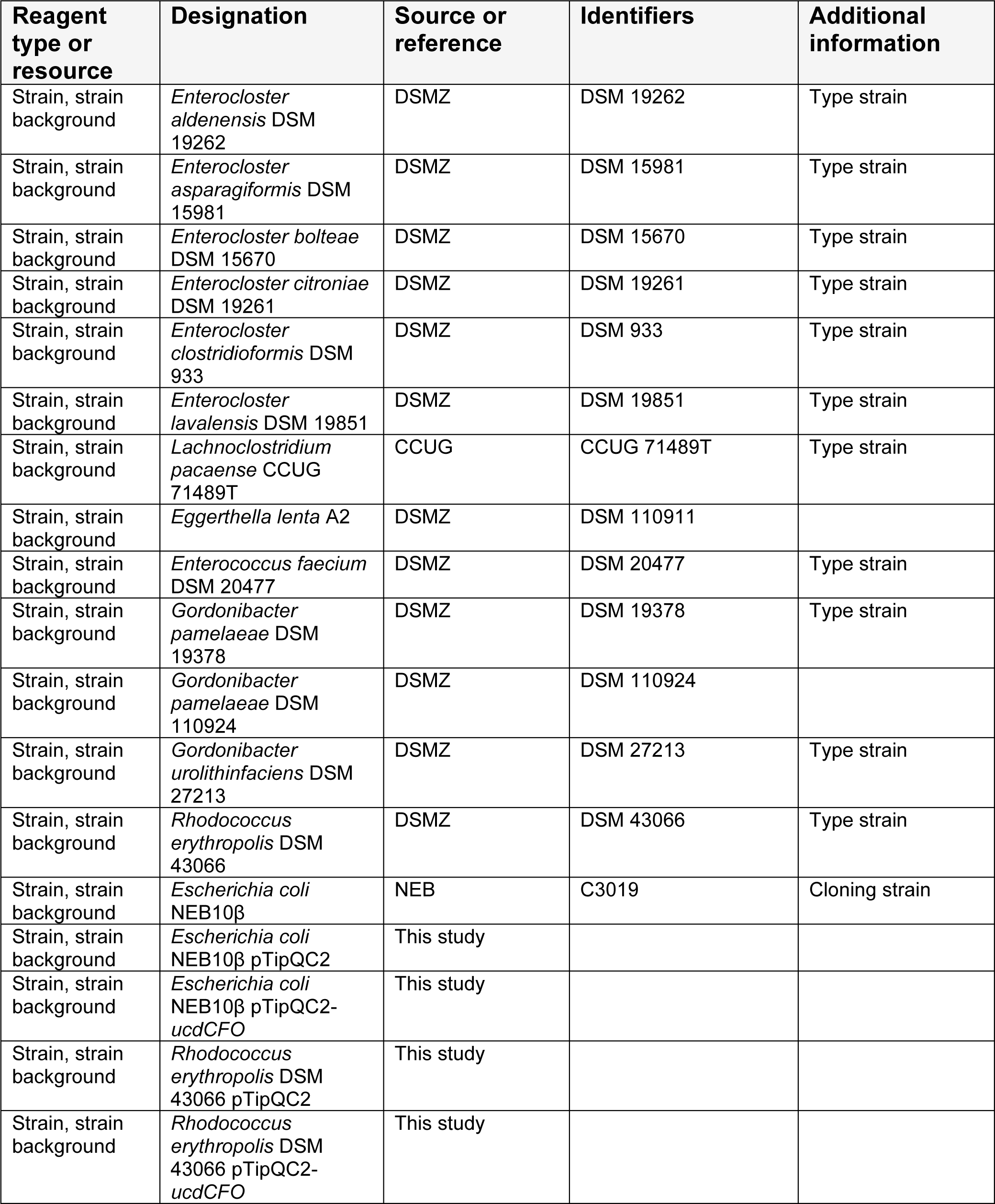
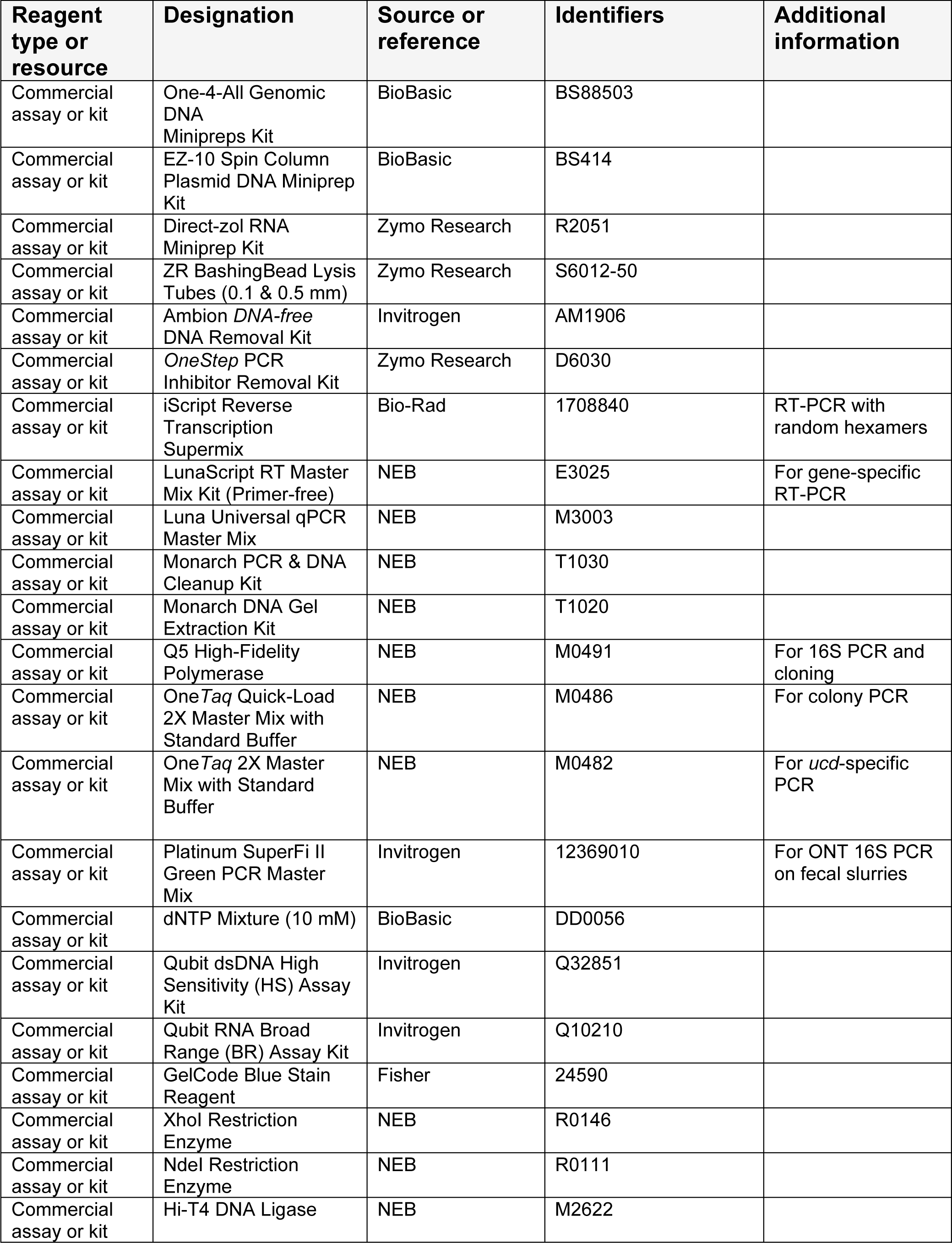
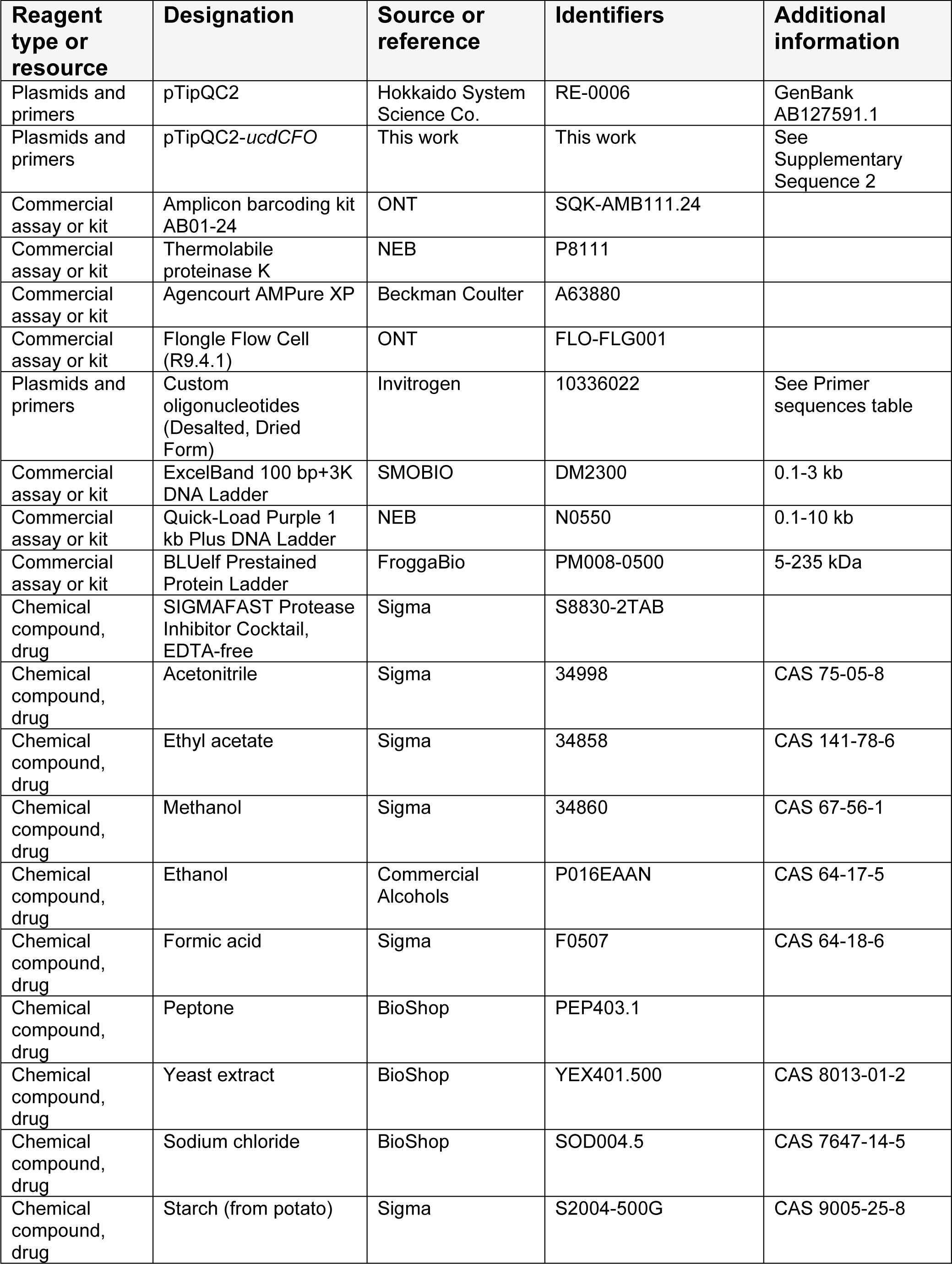
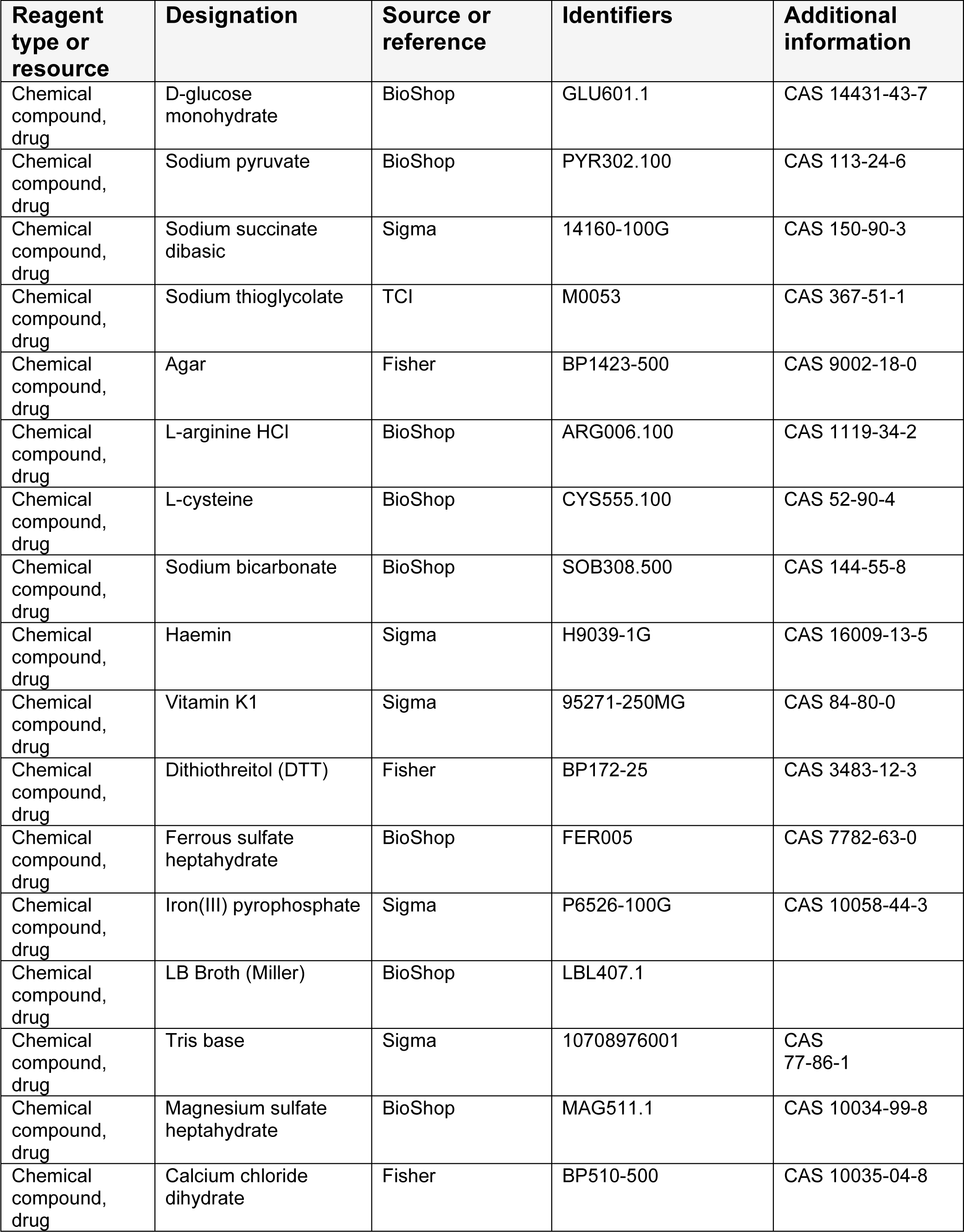
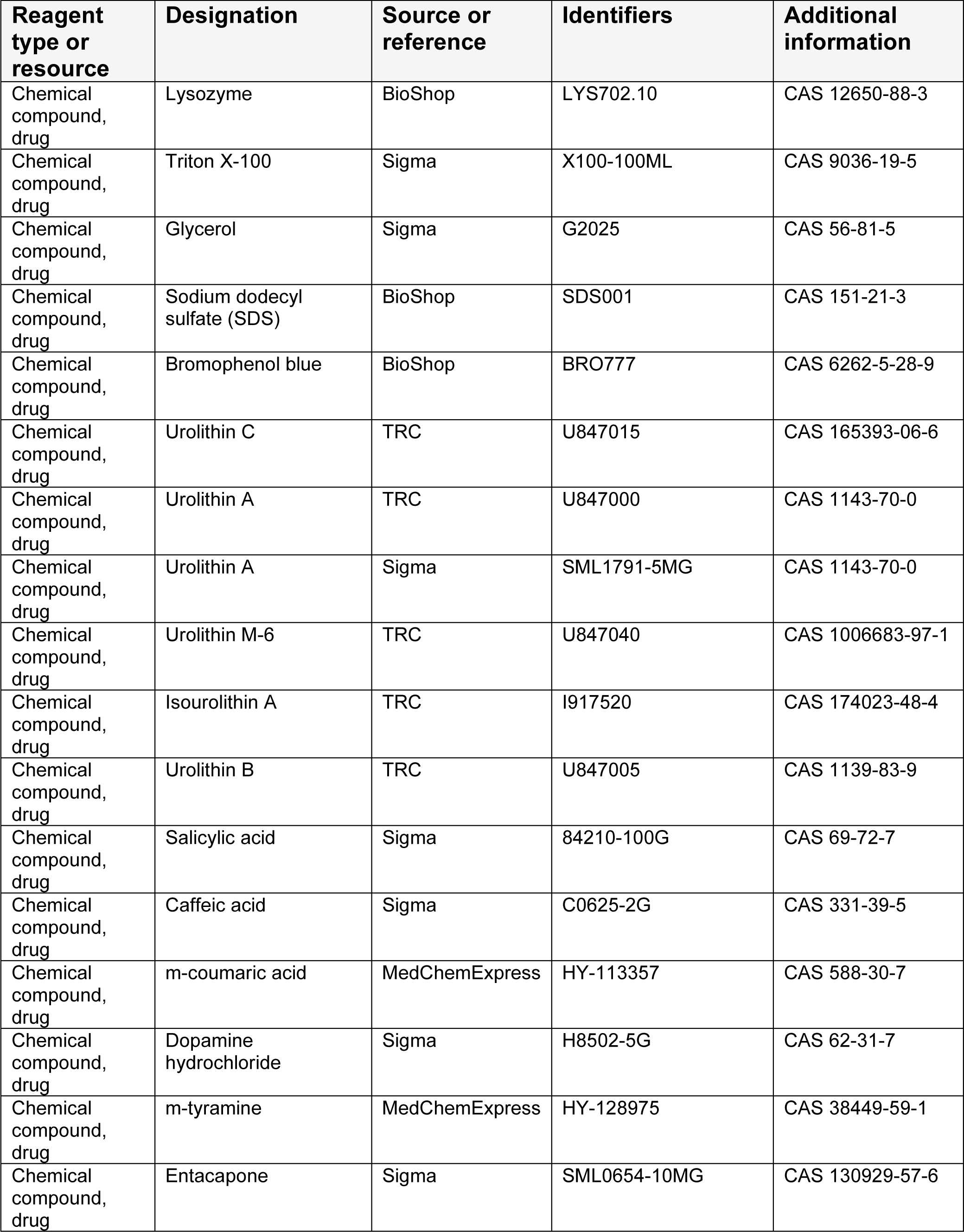
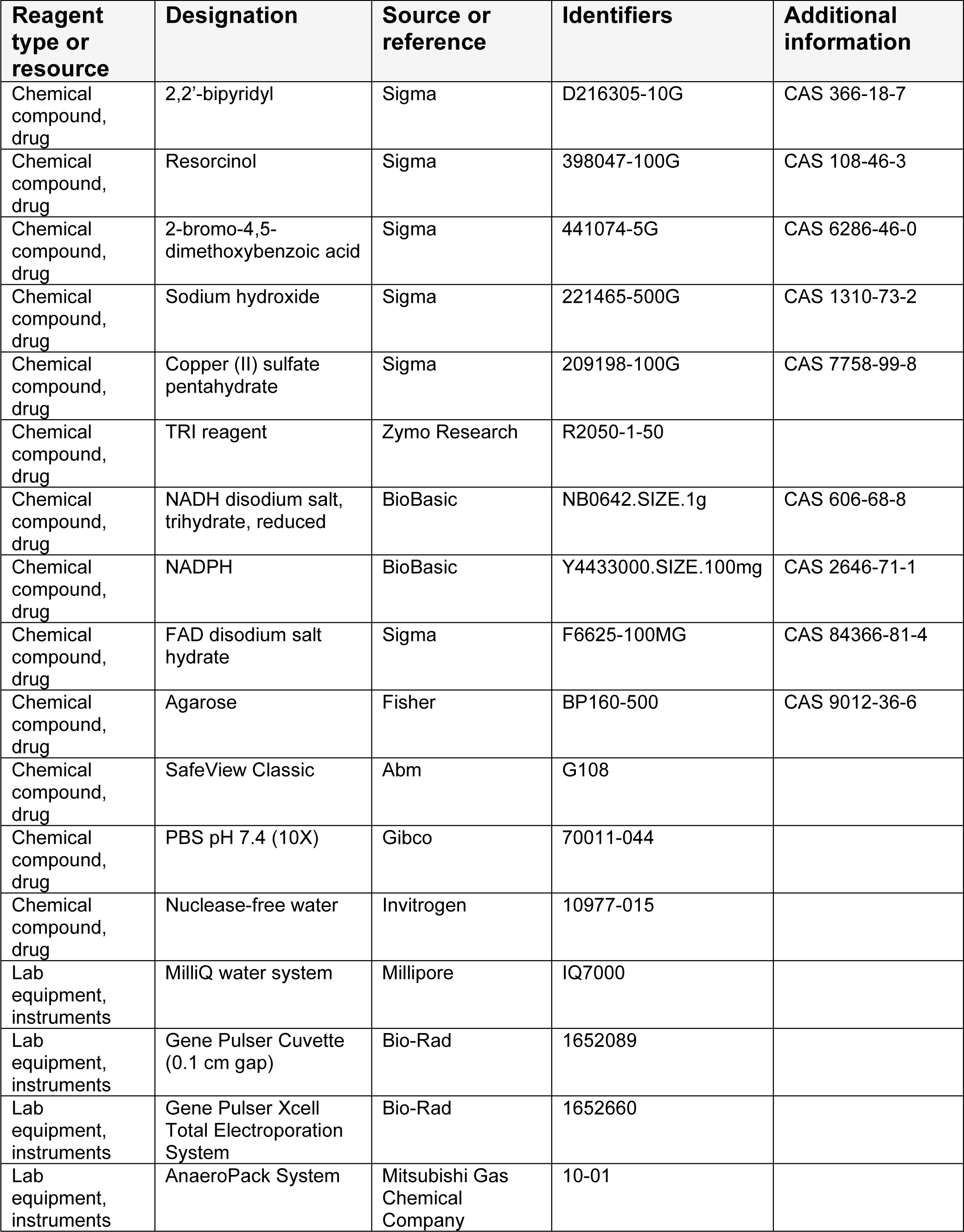
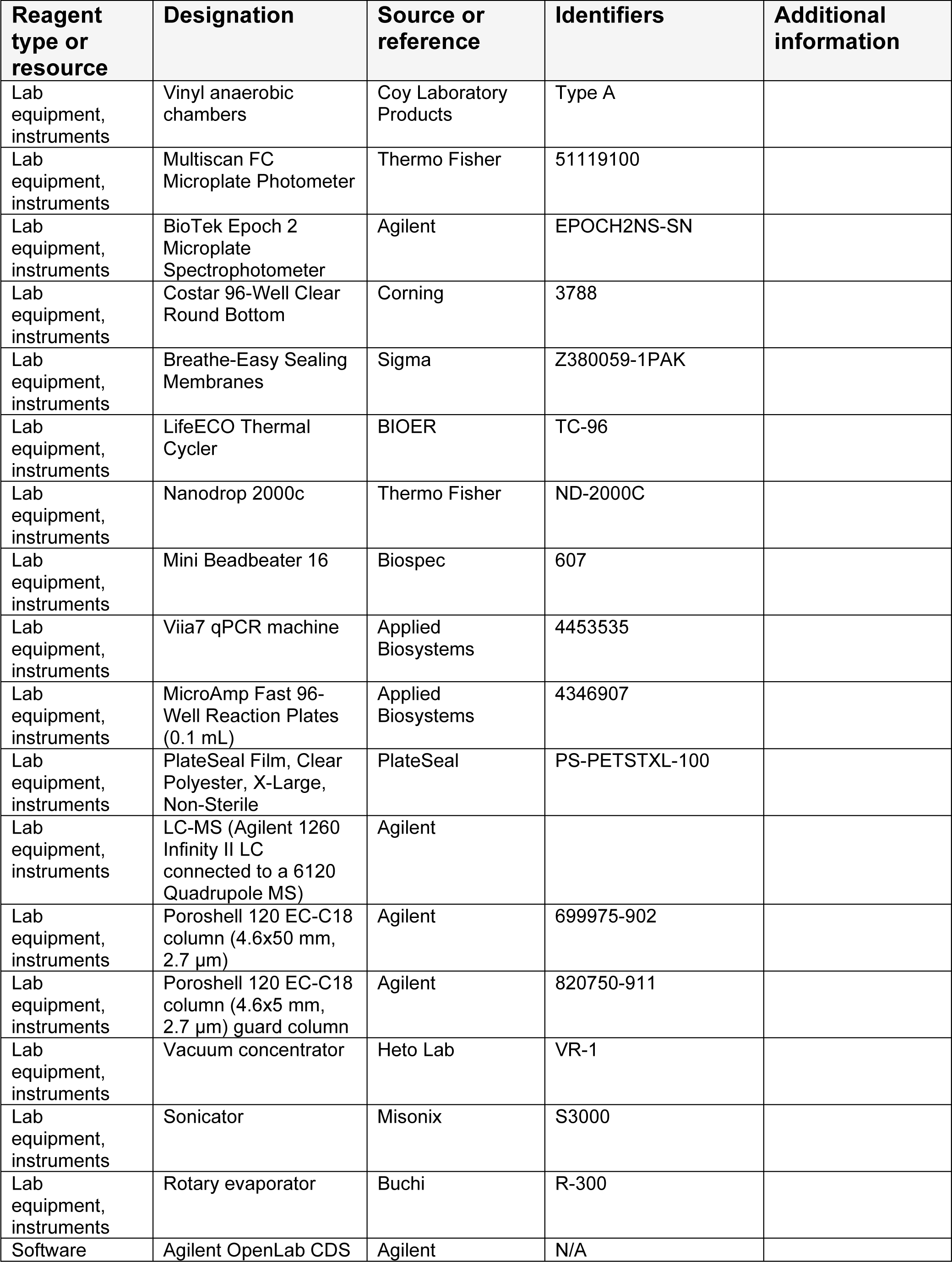
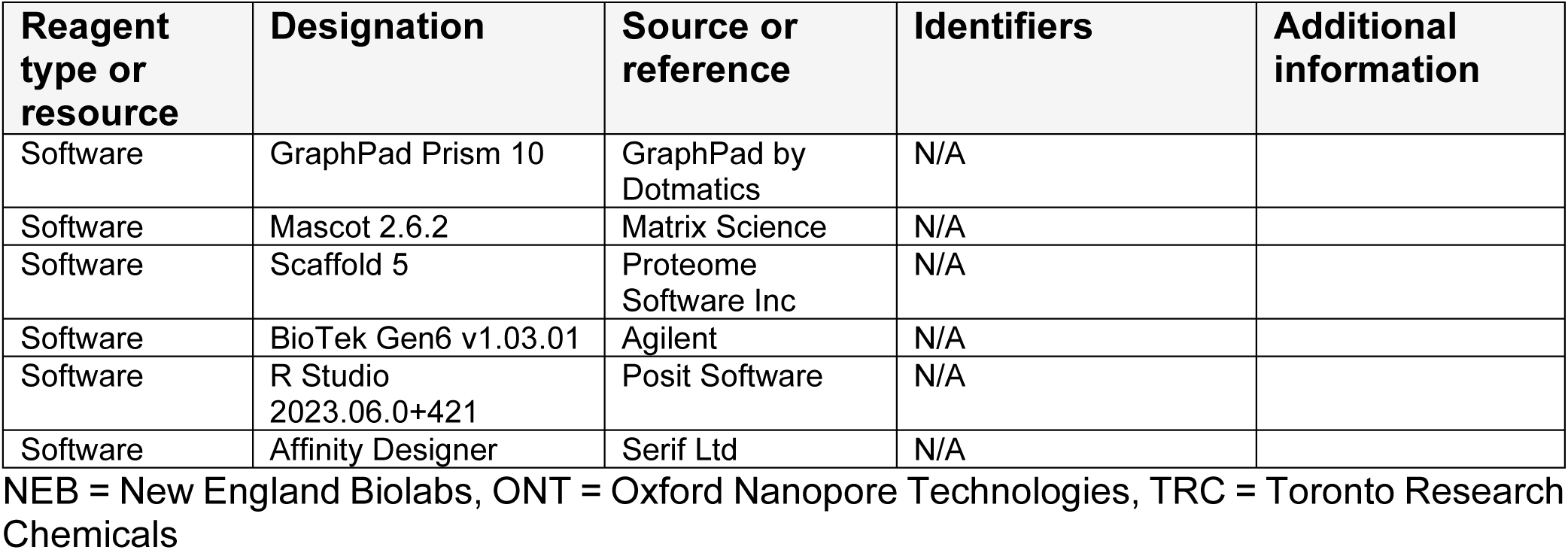

### Primer sequences table

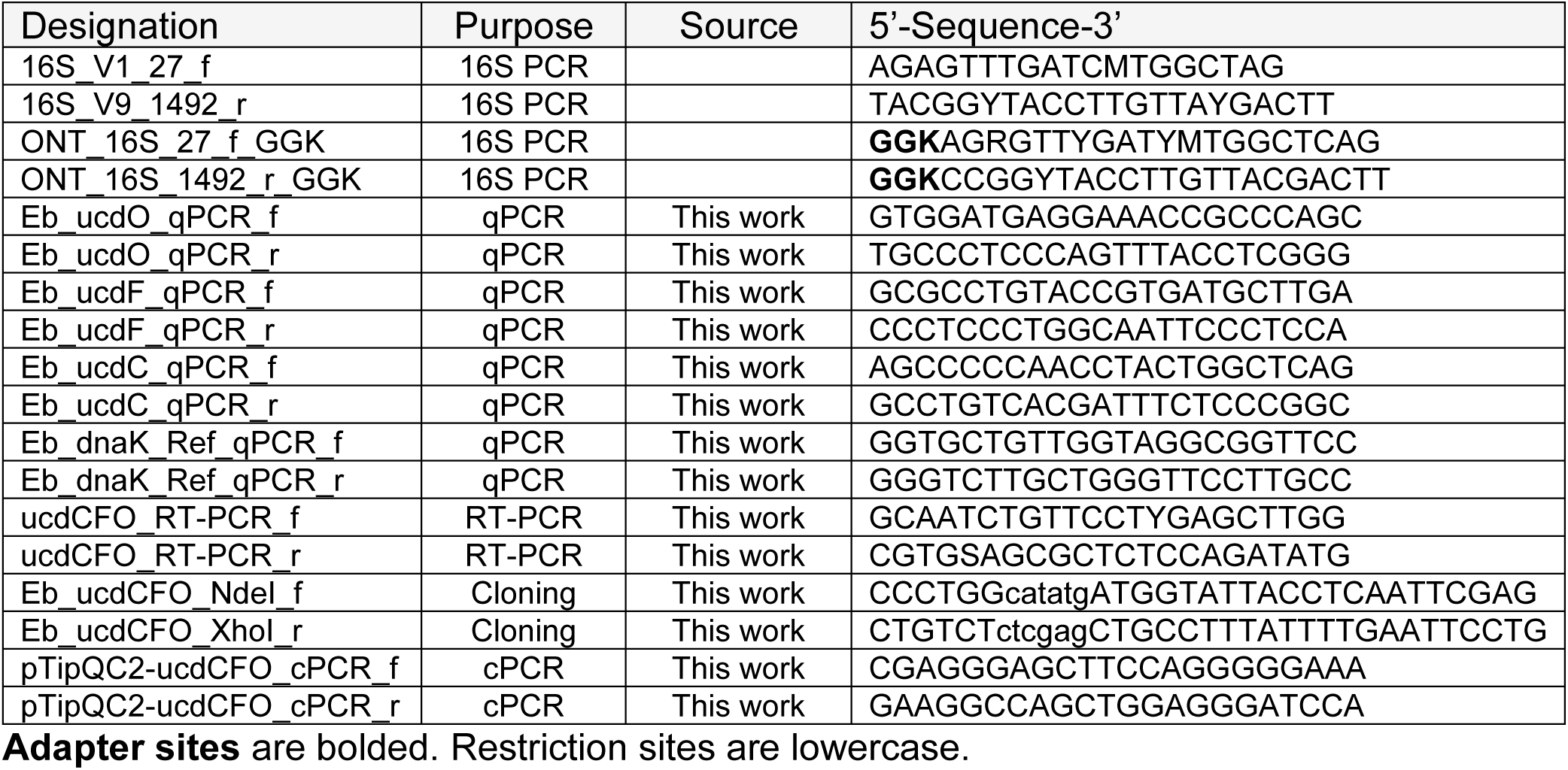

### Anaerobic bacterial strains and culturing conditions

Bacterial strains used in this study are listed in the *Resources table*. All bacterial stains were validated by sequencing the 16S rRNA gene (see Genomic DNA extraction and 16S rRNA sequencing of bacterial isolates). The same culture used for validation was used to make 25% glycerol stocks. Anaerobic strains were grown from glycerol stocks on mABB+H agar plates (recipe below) for 48-72 h at 37 °C in a vinyl anaerobic chamber, which was maintained with a gas mixture of 3% H_2_, 10% CO_2_, 87% N_2_. To make overnight cultures, a single colony was inoculated into 5 mL of liquid mABB or mABB+H medium and incubated at 37 °C between 16-48h, depending on the bacterium (16-24 h for *Enterocloster* spp. and *E. faecium*, and 48 h for *E. lenta* and *Gordonibacter* spp.).

### Human fecal sample collection

Human fecal samples were collected under the approval of protocol A04-M27-15B by the McGill Faculty of Medicine Institutional Review Board. Informed written consent was received from the participants for the use of human samples. Eligibility criteria for the healthy participants were as follows: body mass index between 18.5–30, no diagnosed gastrointestinal disease, no ongoing therapeutic treatment, and no antibiotic usage 3 months prior to the start of the study. Subject information was recorded at the time of sampling. The age of donors ranged from 21–40 years. Fresh fecal samples were collected and placed immediately in an anaerobic chamber, aliquoted, and stored at −70 °C until use.

### Modified anaerobe basal broth (mABB and mABB+H)

For 1 L of modified anaerobe basal broth (mABB), the following components were dissolved in MilliQ water, then autoclaved: 16 g peptone, 7 g yeast extract, 5 g sodium chloride, 1 g starch, 1 g D-glucose monohydrate, 1 g sodium pyruvate, 0.5 g sodium succinate, 1 g sodium thioglycolate, 15 g agar (for plates). The autoclaved solution was allowed to cool, then the following filter-sterilized solutions were added aseptically: 10 mL of 100 mg/mL L-arginine-HCl, 10 mL of 50 mg/mL L-cysteine, 8 mL of 50 mg/mL sodium bicarbonate, 50 μL of 10 mg/mL vitamin K1, 20 mL of 50 mg/mL dithiothreitol, and, for mABB+H, 10 mL of 0.5 mg/mL haemin. The media was then placed in the anaerobic chamber and allowed to reduce for at least 24 h prior to its use in experiments.

### Genomic DNA extraction of isolates and identity validation

The identities of all bacteria in this study were validated by full-length (V1-V9) 16S rRNA sequencing. DNA was first extracted from 0.5-1 mL of overnight culture using the One-4-All Genomic DNA Miniprep Kit (BioBasic) according to the manufacturer’s instructions. The purified genomic DNA (2 μL, ∼ 20 ng) was used as a template for PCR reactions (25 μL reaction volume) using the Q5 High-Fidelity polymerase (NEB). PCR tubes were placed in a thermal cycler and targets (∼1.5 kb) were amplified according to the following cycling conditions: 30 s at 98 °C, 30 cycles (10 s at 98 °C, 20 s at 60 °C, 45 s at 72 °C), 2 min at 72 °C, and hold at 10 °C. 5 μL of the reaction was mixed with 6X loading buffer and loaded onto a 1% agarose gel (made with 1X TAE buffer) containing SafeView Classic (Abm). PCR product sizes were compared to the ExcelBand 100 bp+3K DNA Ladder (SMOBIO).

PCR products (∼1.5 kb) were purified using the Monarch PCR & DNA Cleanup Kit (NEB) according to the manufacturer’s instructions for products < 2 kb. Purified 16S PCR products were eluted in nuclease-free water, quantified using the Qubit dsDNA HS assay kit (Invitrogen), and adjusted to 30 ng/μL. Samples were submitted to Plasmidsaurus for long-read sequencing using Oxford Nanopore Technologies (v14 library preparation chemistry, R10.4.1 flow cells).

### Treatments with urolithins and other catechols

All treatments used in this study (urolithin C, urolithin M6, urolithin A, isourolithin A, 8,9-di-O-methyl-urolithin C, dopamine, m-tyramine, caffeic acid, m-coumaric acid, entacapone, and 2,2’-bipyridyl) were dissolved in DMSO to a concentration of 10 mM.

*Treatment prior to growth (metabolism only)*: Overnight cultures of bacteria were diluted 1/50 into fresh mABB+H. Treatments (10 mM stocks solutions, dissolved in DMSO) were added to the diluted bacterial suspension to a final concentration of 100 μM and samples were incubated for 24 h at 37 °C in an anaerobic chamber.

*Treatment during growth (spike-in)*: Overnight cultures of *Enterocloster* spp. were diluted 1/50 into fresh mABB+H and incubated at 37 °C in an anaerobic chamber. After 5 hours of incubation (∼OD_620_ from a 200 μL sample ∼ 0.4), 10 mM urolithins (or an equivalent volume of DMSO) were added to the growing cultures at a final concentration of 50 or 100 μM for protein expression or RNA expression, respectively. For protein expression analyses and inducibility tests, the cultures were incubated for an additional 4 h. For RNA expression analyses, the cultures were incubated for an additional 2 h.

*Treatment prior to growth (growth curves)*: Overnight cultures of *Enterocloster* spp. were diluted 1/25 into mABB (with or without added 15.4 μM iron source), depending on the experimental design. Separately, treatments (10 mM stocks solutions, dissolved in DMSO) were prepared in mABB (with or without added 15.4 μM iron source) to a final concentration of 200 μM. In each well of a 96-well plate, 100 μL of 1/25 bacteria and 200 μM treatment were combined. These were plated in technical duplicates. The final concentration of treatment was 100 μM (unless otherwise specified in concentration-response experiments) and the final dilution of bacteria was 1/100.

### Urolithin extraction from fecal slurries or bacterial cultures

Frozen (-70 °C) fecal slurries or bacterial cultures were thawed at room temperature. For quantification of urolithin concentrations, urolithin standards (stock 10 mM in DMSO) were spiked into separate media aliquots immediately before extraction.

*Extraction Method A*: This method was used for cultures. Salicylic acid (3 mg/mL in DMSO) was spiked-in as an internal standard at a final concentration of 50 μg/mL. The cultures and standards were then extracted with 3 volumes of ethyl acetate + 1% formic acid (e.g., 600 μL solvent to 200 μL thawed culture). The organic phase (top) was transferred to a new tube and dried in a vacuum concentrator (Heto Lab) connected to a rotary evaporator (Buchi). After solvent removal, samples were redissolved in 0.5 volumes (relative to the starting culture) of 50% MeOH:H_2_O. Samples were centrifuged at 20,000 x g for 5 min to pellet insoluble material, then transferred to LC-MS vials. Urolithins were then analyzed by LC-MS.

*Extraction Method B*: This method was used for cultures and pre-induced cell suspensions. Samples were diluted with an equal volume of MeOH, vortexed briefly, and incubated at room temperature for 10 min. Samples were centrifuged at 20,000 x g for 5 min to pellet insoluble material, then transferred to LC-MS vials. Urolithins were then analyzed by LC-MS.

*Extraction Method C*: This method was used to extract urolithins from crude bacterial lysates. Lysates were diluted with 3 volumes of MeOH, vortexed briefly, and incubated at room temperature for 10 min. Samples were centrifuged at 20,000 x g for 5 min to pellet insoluble material, then transferred to LC-MS vials. Urolithins were then analyzed by LC-MS.

### LC-MS method to quantify urolithins

Samples (10 μL) were injected into a 1260 Infinity II Single Quadrupole LC/MS system (Agilent) fitted with a Poroshell 120 EC-C18 4.6x50 mm, 2.7 μm column (Agilent). The mobile phase was composed of MilliQ water + 0.1% formic acid (solvent A) and acetonitrile + 0.1% formic acid (solvent B). The flow rate was set to 0.7 mL/min. The gradient was as follows: 0-8 min: 10-30 %B, 8-10 min: 30-100 %B, 10-13.5 min: 100 %B isocratic, 13.5-13.6 min: 100-10 %B, then 13.6-15.5 min: 10 %B. The multiple wavelength detector was set to monitor absorbance at 305 nm. The mass spectrometer was run in negative mode in both selected ion monitoring (SIM) and scan (100-1000 m/z) modes to validate peak identities. Peaks were validated based on retention times compared to spike-in standards and mass-to-charge ratios. To quantify urolithins, peak areas for the compounds of interest were compared to spike-in standards of known concentration(s). When standards were not available (urolithin M7), the extracted ion chromatogram was used ([M-H]^-^ = 243).

### Synthesis of di-O-methyl-urolithin C

Di-O-methyl-urolithin C (3-Hydroxy-8,9-dimethoxy-6H-dibenzo[b,d]pyran-6-one, CAS 126438-35-5) was synthesized based on previously described Ullmann-type coupling conditions for urolithin derivatives ^60^. Resorcinol (213 mg, 2 mmol) and 2-bromo-4,5-dimethoxybenzoic acid (261 mg, 1 mmol) were dissolved in 1 mL of 8% w/v NaOH (in MilliQ H_2_O) and heated in a thermo-shaker set to 100 °C for 20 min (in 1.7 mL tube). Then, 200 μL of a 10% w/v Cu(II)SO_4_ pentahydrate solution was added and the reaction was heated at 100 °C for 1 h. The reaction solution (pink-red coloration) contained an insoluble precipitate which was collected by centrifugation (20,000 x g for 30 s). The insoluble pellet was washed 7 times with 1 mL of MilliQ H_2_O until the pH of the wash solution was equal to the pH of MilliQ H_2_O (∼pH 6). The pellet was dried by lyophilization for 16 h (0.0010 mbar, -90 °C) and the product was recovered as a pale pink solid (94 mg, 35% yield). ^1^H NMR (600 MHz, (CD_3_)_2_SO)): δ = 10.22 (s, 1H), 8.21 (d, J = 8.76 Hz, 1H), 7.69 (s, 1H), 7.54 (s, 1H), 6.83 (dd, J = 8.67, 2.37 Hz, 1H), 6.74 (d, J = 2.34 Hz, 1H), 4.02 (s, 3H), 3.89 (s, 3H); HRMS: m/z [M+Na]^+^ calculated for C_15_H_12_NaO_5_: 295.0577, found: 295.0585.

### Phylogenetic tree construction

Phylogenetic trees based on the 16S rRNA gene were constructed using the DSMZ single-gene phylogeny server (https://ggdc.dsmz.de/phylogeny-service.php#)^61^ with GenBank 16S rRNA sequence accessions: *E. aldenensis* (DQ279736), *E. asparagiformis* (AJ582080), *E. bolteae* (AJ508452), *E. citroniae* (HM245936), *E. clostridioformis* (M59089), *E. lavalensis* (EF564277), *L. pacaense* (LT799004), *G. pamelaeae* (AM886059), *G. urolithinfaciens* (HG000667), *E. isourolithinifaciens* (MF322780).

Phylogenetic trees based on whole genomes and proteomes were constructed using the Type (Strain) Genome Server (TYGS, https://tygs.dsmz.de)^40,62^ with the following GenBank genome accessions: *E. aldenensis* (GCA_003467385.1), *E. asparagiformis* (GCA_025149125), *E. bolteae* (GCA_000154365), *E. clostridioformis* (GCA_900113155), *E. lavalensis* (GCA_003024655), *L. pacaense* (GCA_900566185). For *E. citroniae*, the Integrated Microbial Genomes ObjectID was used: *E. citroniae* (2928404274). Further information on nomenclature and taxonomy was obtained from the List of Prokaryotic names with Standing in Nomenclature (LPSN, available at https://lpsn.dsmz.de).

### Cell suspension assay to test inducibility

Bacteria (10 mL growing cultures in mABB+H media) were grown with 50 μM uroC (or an equivalent volume of DMSO) as detailed in the *Treatment during growth (spike-in)* section above and incubated for 4 h at 37 °C. Cultures were then pelleted at 6,500 x g for 3 min and the supernatants were discarded. The cells were washed with 10 mL of pre-reduced PBS (placed in the anaerobic chamber 24 h before), re-pelleted, and resuspended in 2 mL of pre-reduced PBS. For each condition tested, a 200 μL aliquot of the cell suspension was transferred into a sterile 1.5 mL tube, and 10 mM urolithins (uroM6, uroC, isouroA, or DMSO) were added at a final concentration of 100 μM. Cell suspensions were briefly vortexed and incubated at room temperature in the anaerobic chamber for 16h prior to freezing and urolithin extraction using *Extraction Method B*.

### RNA extraction from isolates

A volume of 1.5 mL of treated (100 μM urolithin C for 2 h) *Enterocloster* spp. culture (see *Enterocloster* spp. urolithin C treatments) was pelleted (6,500 g for 3 min) and the supernatant was removed for later LC-MS analysis. The pellet (suspended in 200 μL of media) was then mixed with 800 μL TRI reagent (Zymo Research) and transferred to a ZR BashingBead lysis tube (Zymo Research). Samples were lysed in a Mini Beadbeater 16 (Biospec) according to the following sequence: 1 min ON, 5 min OFF. For RNA-sequencing, bead beating was done for a total of 5 min ON. For RT-(q)PCR, bead beating was done for a total of 2 min ON to preserve longer transcripts. RNA isolation was then performed using the Direct-zol RNA Miniprep Kit (Zymo Research) according to the manufacturer’s instructions (including an on-column DNase digestion). To ensure complete DNA removal, an additional DNA digestion step was performed on the isolated RNA using the Ambion *DNA-free* DNA Removal Kit (Invitrogen) according to the manufacturer’s instructions. The DNA-free RNA was then cleaned up using the *OneStep* PCR Inhibitor Removal Kit (Zymo Research). RNA concentration and quality were initially verified by NanoDrop and 1 % agarose gel electrophoresis. For RNA-sequencing, RNA integrity was assessed by Génome Québec using a Bioanalyzer 2100 (Agilent). RNA integrity (RIN) values ranged between 7.5-7.8 for *E. asparagiformis* DSM 15981 and 7.0-7.3 for *E. bolteae* DSM 15670.

### RNA-sequencing of urolithin C-treated *E. bolteae* and *E. asparagiformis* isolates

Total RNA was sent to Génome Québec for library preparation and RNA-sequencing. Briefly, total RNA was prepared for Illumina sequencing using the NEBNext rRNA Depletion Kit (Bacteria) (NEB) kit to remove rRNA and using the NEBNext Multiplex Oligos for Illumina (NEB) kit (stranded/directional). Prepared libraries were quality checked with a Bioanalyzer 2100 (Agilent) prior to sequencing. Sequencing was performed on a NovaSeq 6000 (Illumina) with the following flow cell/settings: S4 flow cell, 100 bp, 25 M reads, paired end.

Analysis of RNA-seq reads was done using Galaxy bioinformatics cloud computing (https://usegalaxy.org/) hosted by Compute Canada Genetics and Genomics Analysis Platform (GenAP) (https://www.genap.ca/). Genomes and annotations were fetched from the NCBI genome browser: *E. bolteae* (ASM223457v2) (accessed 2022/05/11) and *E. asparagiformis* (ASM2514912v1) (accessed 2023/09/19). Raw reads were first verified for quality using FastQC (v0.73, https://www.bioinformatics.babraham.ac.uk/projects/fastqc/) with default parameters. FastQC reports were aggregated into MultiQC (v1.11) ^63^. The mean sequence quality scored were above 35 for all samples. Raw reads were then trimmed using Cutadapt (v3.7) ^64^ to trim adapter sequences (R1 sequence: AGATCGGAAGAGCACACGTCTGAACTCCAGTCAC, and R2 sequence: AGATCGGAAGAGCGTCGTGTAGGGAAAGAGTGT) that were not removed after sequencing using default parameters for paired end reads. Trimmed reads were then aligned to reference genomes for each bacterium using HISAT2 (v2.2.1) ^65^ with paired-end parameters and reverse strandedness (RF). Aligned read counts were assigned to features in annotation files (.gtf) using featureCounts (v2.0.1) ^66^ with the following parameters: reverse strandedness, count fragments instead of reads, GFF feature type filter = “gene”, multi-mapping and multi-overlapping features included (-M -O), minimum mapping quality per read of 0, and the rest of the parameters were kept as default. Differential gene expression analysis was then performed using DESeq2 (v2.11.40.7) ^67^ using default parameters. Differential expression tables were annotated with the Annotate DESeq2/DEXSeq output tables tool (v1.1.0) in Galaxy to include the following: GFF feature type = “CDS”, GFF feature identifier = “gene_id”, GFF transcript identifier = “transcript_id”, GFF attribute to include = “protein_id, product”. The “protein_id” was used to query the NCBI database and the NCBI Sequence Viewer was used to investigate the genomic context surrounding genes of interest.

### Comparative genomics

The nucleotide sequence for the *Enterocloster bolteae* DSM 15670 *ucd* operon (NCBI NZ_CP022464 REGION: complement(4417875..4421605)) was used as a query for BLASTn (megablast) searches using the refseq_genomes database limited to Bacteria (taxid:2). The NCBI multiple sequence alignment (MSA) viewer was used to download alignment figures.

### RT-PCR analysis of *E. bolteae* to determine *ucd* operon structure

Isolated RNA samples (500 ng) were reverse transcribed using the LunaScript® RT Master Mix Kit (Primer-free) (NEB) in a reaction volume of 10 µL containing the ucdCFO_RT-PCR_r primer at a final concentration of 1 µM. The No-RT Control included in the kit was used as a no-enzyme control for reverse transcription. The reaction mixtures were incubated in a thermal cycler: 10 min at 55°C, 1 min at 95°C. PCR reactions were conducted using the OneTaq 2X Master Mix with Standard Buffer (NEB). The ucdCFO_RT-PCR primer pair was added to the master mix (to a final concentration of 0.2 µM) and 1 µL of template (cDNA, -RT, no template, or gDNA) was added for a total reaction volume of 20 µL. PCR tubes were placed in a thermal cycler and targets were amplified according to the following conditions: 20s at 94°C, 31 cycles (20s at 94°C, 30s at 62°C, 3 min at 68°C), 5 min at 68°C. A volume of 5 μL of reaction was directly loaded onto a 1% agarose gel (made with 1X TAE buffer) containing SafeView Classic (Abm). PCR product sizes were compared to the Quick-Load® Purple 1 kb Plus DNA Ladder (NEB). The rest of the PCR product was then run on a 1% agarose gel and bands corresponding to the desired products were cut out and purified using the Monarch DNA Gel Extraction kit (NEB). DNA was quantified using the Qubit dsDNA HS assay kit (Invitrogen) and submitted to Plasmidsaurus for long-read sequencing using Oxford Nanopore Technologies (Supplementary Sequence 1).

### RT-qPCR analysis of *E. bolteae ucd* operon genes

Isolated RNA samples (500 ng) were reverse transcribed using the iScript Reverse Transcription Supermix (Bio-Rad) in a reaction volume of 10 μL. The iScript No-RT Control Supermix was used as a no enzyme control for reverse transcription (-RT). The reaction mixtures were incubated in a thermal cycler: 5 min at 25 °C, 20 min at 48 °C, and 1 min at 95 °C. Both cDNA and -RT controls were diluted 1/20 in nuclease-free water before use. qPCR reactions were conducted using the Luna Universal qPCR Master Mix kit (NEB). The Eb_ucdO_qPCR, Eb_ucdF_qPCR, Eb_ucdC_qPCR, and Eb_dnaK_Ref_qPCR primer pairs were added to their respective master mixes (final primer concentration of 250 nM) and 6.6 or 4.4 μL of diluted template (cDNA, -RT, no template) were added to 26.4 or 17.6 μL of master mix for triplicates or duplicates, respectively. All cDNA samples were run in technical triplicates, while other sample types were run in technical duplicates. Replicate mixes were pipetted (10 μL/well) into a MicroAmp Fast 96-Well Reaction Plate (Applied Biosystems) and the plates were sealed, then spun down for 2 min to eliminate air bubbles. The qPCR detection parameters were as follows: SYBR Green detection, ROX reference dye, 10 μL reaction volume. The thermal cycling conditions were: 1 min at 95 °C, 40 cycles (15 s at 95 °C, 30 s at 60 °C), then melt analysis (60-95 °C). Data were analyzed according to the 2^-ΔΔCt^ method ^68^ with the dnaK gene serving as the reference gene (*E. bolteae* dnaK RNA-seq log_2_FC = 0.122).

### Protein extraction from *Enterocloster* spp

All steps other than sonication were carried out under anaerobic conditions. To extract proteins, 10 mL of treated (50 μM urolithin C for 4 h) *Enterocloster* spp. culture (see *Enterocloster* spp. urolithin C treatments) were pelleted (6,500 g for 3 min) and the supernatant was discarded. The pellet was washed with 10 mL of pre-reduced PBS, pelleted again, and resuspended in 0.5 mL of pre-reduced lysis buffer (20 mM Tris, pH 7.5, 500 mM NaCl, 10 mM MgSO_4_, 10 mM CaCl_2_, and 1 tablet/100 mL SIGMAFAST protease inhibitor (EDTA-free)). The resuspended pellet was then sonicated on ice using a Misonix Sonicator 3000 set to power level 2/10 according to the following sequence (aerobically, in a cold room): 20 s ON, 40 s OFF, for a total of 2 min ON. Tubes were centrifuged at 20,000 x g for 2 min to pellet insoluble particles and 0.4 mL of lysate was transferred to a new 1.5 mL tube (kept on ice). Lysates used in metabolism assays were transported to the anaerobic chamber in a sealed plastic bag containing an anaerobic gas generating system to minimize loss in activity.

### Urolithin metabolism assays using crude lysates from uroC-induced *E. bolteae*

Protein lysates (described above) were aliquoted (50 μL aliquots) into 1.5 mL tubes, then treated with DMSO or urolithin C (10 mM stock) at a final concentration of 350 μM. Cofactors (NADPH, NADH, and FAD, each dissolved to a final concentration of 30 mM (in lysis buffer immediately before the assay was run) were added individually to the lysates at a final concentration of 1 mM. The lysates were incubated at room temperature in an anaerobic chamber for 20 h prior to freezing at -70 °C. Samples were then extracted using *Extraction Method C*.

To assess the oxygen sensitivity of crude lysates from uroC-induced *E. bolteae*, samples were prepared as described above. After adding DMSO or uroC and NADH (under anaerobic conditions), tubes were either incubated at room temperature inside the anaerobic chamber or just outside of the chamber for 20 h. Afterwards, samples were frozen at -70 °C and then extracted using *Extraction Method C*.

### Proteomics analysis of uroC-treated *E. bolteae*

Extracted proteins were submitted for proteomic analysis at the RI-MUHC. For each sample, protein lysates were loaded onto a single stacking gel band to remove lipids, detergents, and salts. The gel band was reduced with DTT, alkylated with iodoacetic acid, and digested with trypsin. Extracted peptides were re-solubilized in 0.1% aqueous formic acid and loaded onto a Thermo Acclaim Pepmap (Thermo, 75 um ID X 2 cm C18 3 um beads) precolumn and then onto an Acclaim Pepmap Easyspray (Thermo, 75 um ID X 15 cm with 2 um C18 beads) analytical column separation using a Dionex Ultimate 3000 uHPLC at 250 nL/min with a gradient of 2-35% organic (0.1% formic acid in acetonitrile) over 3 hours. Peptides were analyzed using a Thermo Orbitrap Fusion mass spectrometer operating at 120,000 resolution (FWHM in MS1) with HCD sequencing (15,000 resolution) at top speed for all peptides with a charge of 2+ or greater. The raw data were converted into *.mgf format (Mascot generic format) for searching using the Mascot 2.6.2 search engine (Matrix Science) against *Enterocloster bolteae* DSM 15670 proteins (NCBI assembly GCF_002234575.2) and a database of common contaminant proteins. Mascot was searched with a fragment ion mass tolerance of 0.100 Da and a parent ion tolerance of 5.0 ppm. O-63 of pyrrolysine, carboxymethyl of cysteine and j+66 of leucine/isoleucine indecision were specified in Mascot as fixed modifications. Deamidation of asparagine and glutamine and oxidation of methionine were specified in Mascot as variable modifications.

The database search results were loaded into Scaffold Q+ Scaffold_5.0.1 (Proteome Sciences) for statistical treatment and data visualization. Scaffold (v5.3.0) was used to validate MS/MS based peptide and protein identifications. Peptide identifications were accepted if they could be established at greater than 95.0% probability by the Peptide Prophet algorithm ^69^ with Scaffold delta-mass correction. Protein identifications were accepted if they could be established at greater than 99.0% probability and contained at least 2 identified peptides. Protein probabilities were assigned by the Protein Prophet algorithm ^70^. Proteins that contained similar peptides and could not be differentiated based on MS/MS analysis alone were grouped to satisfy the principles of parsimony. Proteins sharing significant peptide evidence were grouped into clusters. Protein quantification and differential expression were determined in Scaffold using the following parameters: Quantitative method was set to total spectra, the minimum value was set to 0.5 in case proteins were not detected in one condition, and statistical tests were performed using Fisher’s exact test with the Benjamini-Hochberg multiple test correction at a significance level set to 0.05.

### Protein structures

The protein FASTA sequences (NCBI RefSeq accessions for UcdC: WP_002569575.1, UcdF: WP_002569574.1, UcdO: WP_002569573.1) for the *Enterocloster bolteae* DSM 15670 *ucd* operon were used as a query for BLASTp searches against the UniProtKB reference proteomes + Swiss-Prot databases. The AlphaFold2 protein structures for matches (UniProt accessions for UcdC: A8RZR5, UcdF: A8RZR3, UcdO: A8RZR2) of each protein were downloaded and imported into PyMOL (v2.4.1). Foldseek (in 3Di/AA mode) was used to generate a list of proteins with similar structures from solved crystal structures in the Protein Data Bank (PDB) ^48^.

Hits of published X-ray crystal structures (PDB: 1ZXI (from *Afipia carboxidovorans* OM5) and PDB: 3UNI (from *Bos taurus*)) were fetched from the PDB and imported into PyMOL. The AlphaFold2 structures for each Ucd protein were aligned to the following chains in the published PDBs using the “super” command: PDB 1ZXI: UcdC to chain C, UcdF to chain A, UcdO to chain B; PDB 3UNI: UcdC, UcdF, and UcdO to chain A.

### pTipQC2-*ucdCFO* Plasmid Construction, Purification, and Transformation

*Plasmid construction in E. coli* NEB10β: Primers flanking the *E. bolteae ucd* operon (NCBI NZ_CP022464 REGION: complement(4417875..4421605)) were designed in Benchling using the Primer3 tool. Tails including 6 random bases, followed by restriction sites for NdeI and XhoI were included on the forward and reverse primers, respectively (Eb_ucdCFO_NdeI_f and Eb_ucdCFO_XhoI_r). PCR was performed using the Q5 High-Fidelity polymerase (NEB) with *E. bolteae* DSM 15670 genomic DNA as a template. The target was amplified according to the following cycling conditions: 30 s at 98 °C, 30 cycles (10 s at 98 °C, 20 s at 60 °C, 80 s at 72 °C), 2 min at 72 °C. The *ucdCFO* PCR product was purified using the Monarch PCR & DNA Cleanup Kit (NEB) according to the manufacturer’s instructions for products ≥ 2 kb. The resulting purified PCR product and the pTipQC2 plasmid (Hokkaido Systems Science Co.) were digested overnight (16 h) with NdeI and XhoI (both from NEB) in rCutSmart buffer according to the manufacturer’s instructions (∼600-1000 ng DNA per 50 μL reaction). Double digested DNA was migrated on a 0.6% agarose gel and bands corresponding to the desired products were cut out and purified using the Monarch DNA Gel Extraction kit (NEB). The purified products were ligated using the Hi-T4 DNA Ligase (NEB): a ∼3:1 insert:plasmid molar ratio ligation reaction was set up on ice, then incubated at room temperature for 2 h. The ligation mixture (2 μL) was electroporated (1.8 kV, 25 μF, 200 Ω) into 40 μL electrocompetent *E. coli* NEB10β cells (according to the Quick-n’-Dirty Electrocompetent *E. coli* protocol (dx.doi.org/10.17504/protocols.io.bjpykmpw) using 0.1 cm gap cuvettes (Bio-Rad). The cuvette was immediately filled with 1 mL pre-warmed LB post-shock and cells were allowed to recover at 37 °C for 30 min before plating on LB + 100 μg/mL ampicillin. After an overnight incubation at 37 °C, colonies were picked and grown in selective LB + 100 μg/mL ampicillin. Plasmids were purified using the Plasmid DNA Miniprep Kit (BioBasic) and size was confirmed with a diagnostic restriction digest (10 μL reactions). The final plasmid construct (pTipQC2-*ucdCFO*) was submitted to Plasmidsaurus for long-read sequencing using Oxford Nanopore Technologies (v14 library preparation chemistry, R10.4.1 flow cells) (Supplementary Sequence 2).

*pTipQC2-ucdh transformation into Rhodococcus erythropolis DSM 43066*: Electrocompetent *R. erythropolis* DSM 43066 were prepared according to a modified protocol from <u>P. Lessard 2002</u>. Briefly, 50 mL LB were inoculated with 1 mL of a stationary phase (48-72 h growth from a single colony) *R. erythropolis* DSM 43066 culture and grown aerobically for 16 h at 30 °C with shaking at 200 RPM. The next day, cells were pelleted at 5,000 x g for 10 min at 4 °C and washed according to the following sequence: 2 washes of (10 mL of ice cold sterile MilliQ water), 10 mL of ice cold sterile 10% glycerol. The final pellet was then resuspended in 5 mL of ice cold sterile 10% glycerol. The resuspended electrocompetent *R. erythropolis* DSM 43066 were aliquoted (50 μL/aliquot), then 3 μL (∼0.5-1 μg) of pTipQC2-*ucdCFO* plasmid was added to appropriate tubes and incubated for 30 min on ice. Cells with plasmid were transferred to 0.1 cm gap cuvettes (Bio-Rad) and electroporated (1.8 kV, 25 μF, 200 Ω). Time constants were between 4.3-4.6 ms. The cuvette was immediately filled with 1 mL LB post shock and cells were allowed to recover at 30 °C for 2.5 h before plating 100 μL dilutions (1/10 dilution, undiluted, and concentrated recovery culture) on LB + 30 μg/mL chloramphenicol at 30 °C. After 2-3 days of incubation, colonies were picked and grown in selective liquid LB + 30 μg/mL chloramphenicol at 30 °C with shaking at 200 RPM. Plasmid-positive colonies were identified by colony PCR using the pTipQC2-*ucdCFO*_cPCR primer set and validated by diagnostic restriction digests and whole-plasmid sequencing.

### Heterologous expression of UcdCFO enzymes in *Rhodococcus erythropolis*

All growth steps below were performed in selective media (LB + 30 μg/mL chloramphenicol) in aerobic conditions at 30 °C with shaking at 200 RPM, unless otherwise specified. Single colonies of *R. erythropolis* DSM 43066 harboring the pTipQC2 (empty plasmid) or pTipQC2-*ucdCFO* were inoculated into 5 mL selective media and grown for 72 h to produce overnight cultures. Overnight cultures were then thoroughly resuspended and diluted 1:10 into 25 mL fresh selective media and grown for ∼8 h until OD_600_ values reached ∼0.6. Thiostrepton (5 mg/mL in DMSO) was added to a final concentration of 1 μg/mL and cultures were incubated aerobically for 16 h at 25 °C to induce protein expression. The next morning, cultures were pelleted and resuspended in 0.2 volumes of lysis buffer (20 mM Tris, pH 7.5, 500 mM NaCl, 10 mM MgSO_4_, 10 mM CaCl_2_, 2 mM DTT, 1% Triton X-100, 2 mg/mL lysozyme, and 1 tablet/100 mL SIGMAFAST protease inhibitor (EDTA-free)). The resuspended cells in lysis buffer were incubated on ice for 1 h with shaking, then sonicated on ice using a Misonix Sonicator 3000 set to power level 2/10 according to the following sequence (aerobically, in a cold room): 20 s ON, 40 s OFF, for a total of 4 min ON. Crude lysates were transported to the anaerobic chamber in a sealed plastic bag containing an anaerobic gas generating system to minimize loss in activity and treated in the same manner detailed in *Urolithin metabolism assays using crude lysates from uroC-induced E. bolteae*.

### SDS-PAGE analysis of UcdCFO proteins in crude lysates

Crude lysates described above (*Heterologous expression of UcdCFO enzymes in Rhodococcus erythropolis*) were centrifuged for 2 min at 20,000 x g. The insoluble pellet was separated from the soluble supernatant. The insoluble pellet (from 100 μL of crude lysate) was resuspended in 100 μL of 1X reducing loading dye (62.5 mM Tris-HCl (pH 6.8), 2% (w/v) SDS, 10% glycerol, 0.01% (w/v) bromophenol blue, 38 mM DTT). The soluble fraction was diluted with 3X reducing loading dye to a final concentration of 1X. All samples were heated at 95 °C for 5 min, then 10 μL were loaded onto a 10% bis-tris polyacrylamide protein gel. Gels were fixed and stained with GelCode Blue Stain Reagent (Fisher) according to the manufacturer’s instructions.

### Growth curves with catechols in different iron-containing media conditions

Overnight cultures of *Enterocloster* spp. were treated as described in the *Treatment prior to growth (growth curves)* sub-section of *Treatments with urolithins and other catechols*.

Once plated, 96-well plates were sealed with a Breathe-Easy membrane and placed in a pre-warmed plate reader inside the anaerobic chamber (BioTek Epoch 2). The optical density at 620 nm was recorded every 30 min for 48 h. Kinetic analysis was performed in BioTek Gen6 Software using the built-in kinetic analysis.

### curatedMetagenomicData meta-analysis of *Enterocloster ucd* operon in human fecal metagenome datasets

All 93 metagenomic studies (22,588 samples and their metadata) available in the curatedMetagenomicData R package ^53^ (v3.8.0) were downloaded locally (ExperimentHub snapshotDate(): 2023-04-24, accessed on 2023-06-06) and transferred to the Narval cluster hosted by the Digital Research Alliance of Canada. Metagenomic data for urolithin C-metabolizing *Enterocloster* spp. were obtained by querying the "relative_abundance" (pre-processed using MetaPhlAn3 and "gene_families" (pre-processed using HUMAnN3) entries in individual study datasets ^71^. For individual taxa (containing partial strings “bolteae”, “citroniae”, “asparagiformis”, “asparagiforme”, “Enterocloster”, or “47FAA” (corresponding to *L. pacaense*)), relative abundance (%) was extracted from the rows of the "relative_abundance" datasets using the stringr R package (v1.5.0, https://github.com/tidyverse/stringr). Prevalence (relative abundance in sample > 0) was then calculated for each sample.

For specific genes, the NCBI protein accessions for each gene of the *ucd* operon (*ucdO*, *ucdF*, *ucdC*) was used to search the UniProt database. UniRef90 accession numbers corresponding to hits (C5EGQ4, G5HFF3, A8RZR5, respectively) were then extracted from the rows of the "gene_families" datasets using the stringr R package. Prevalence (abundance in sample > 0) was then calculated for each sample. R scripts and RData files are available in Zenodo (see Data Availability).

### Fecal slurry preparation and treatment

Frozen (-70 °C) fecal samples were brought into the anaerobic chamber and allowed to thaw. The samples were suspended in 1 mL mABB medium per 0.1 g feces and homogenized by breaking apart large pieces with a sterile loop and by vortexing. Large particles were pelleted by centrifuging the tubes at 700 x g for 3 min. The supernatants (containing bacteria) were transferred to new tubes and centrifuged at 6,500 x g for 5 min to pellet the cells. The supernatants were discarded, and the cell pellets were washed with 5 mL of fresh media. The cell suspensions were once again centrifuged at 6,500 x g for 5 min and the resulting cell pellets were resuspended in 600 μL media per 0.1 g feces. Resuspended cells were treated with either 100 µM urolithin C or an equivalent volume of DMSO and incubated at 37°C anaerobically for 48 h. 200-300 µL volumes were removed from the batch cultures and immediately frozen at - 70°C for later extraction of urolithins (using *Extraction Method A*), DNA, and RNA.

### Genomic DNA extraction from fecal slurries

A 300 μL fecal slurry aliquot (between 50-100 mg wet weight) was pelleted (10,000 g for 5 min) and the supernatant was removed for later LC-MS analysis. The pellet was then mixed with 750 μL of ZymoBIOMICS lysis solution (Zymo Research) and transferred to a ZR BashingBead lysis tube (Zymo Research). Samples were lysed in a Mini Beadbeater 16 (Biospec) according to the following sequence: 1 min ON, 5 min OFF for a total of 5 min ON. DNA was then purified using the ZymoBIOMICS DNA Miniprep Kit (Zymo Research) according to the manufacturer’s instructions (including the *OneStep* PCR Inhibitor Removal step). Purified DNA samples were quantified using the Qubit dsDNA HS assay kit (Invitrogen).

### Long read 16S rRNA sequencing of microbial communities in fecal slurries

Long read 16S PCR reactions were conducted using the Platinum SuperFi II Green PCR Master Mix (Invitrogen). The ONT_16S_27F_GGK and ONT_16S_1492R_GGK primer pairs (see *Primer sequences table*) were added to their respective master mixes (final primer concentration of 0.2 μM) and 1 μL of template (∼10 ng) was added (for a total reaction volume of 25 μL). PCR tubes were placed in a thermal cycler and targets were amplified according to the following cycling conditions: 30 s at 98 °C, 30 cycles (10 s at 98 °C, 10 s at 60 °C, 30 s at 72 °C), 5 min at 72 °C, and hold at 4 °C. Amplicons were quantified using the Qubit dsDNA HS assay kit (Invitrogen) to verify that amplicon concentrations were reasonably balanced (range = 18.36-24.00 ng/μL). Barcoding of amplicons was performed with 2 μL of PCR reaction according to the manufacturer’s instructions (for ONT kit SQK-AMB111-24). Barcoding reactions were incubated in a thermal cycler for 10 min at 65 °C, then for 2 min at 80 °C. 10 μL of each barcoding reaction were pooled and proteins were digested using heat-labile proteinase K (NEB) by incubating the pooled library for 15 min at 37 °C, followed by heat inactivation for 10 min at 55 °C. Amplicons were purified using Agencourt AMPure XP beads (Beckman Coulter) using 0.7 volumes of beads-to-library. Following 70% EtOH washes and drying steps, the library was eluted using 15 μL of the provided elution buffer (EB), yielding a library with a concentration of 30 ng/μL using the Qubit dsDNA HS assay kit (Invitrogen). 11 μL of the eluted

DNA library were transferred to a new tube and combined with 1 μL of Rapid Adapter T (RAP T). This mixture was incubated at room temperature for 10 min. Since the library was concentrated, it was diluted 1:2 in EB before combining with SB II and LB II, then loaded into a primed Flongle Flow Cell (R9.4.1) in a MinION device following the manufacturer’s instructions. Sequencing was allowed to proceed for ∼20 h until pore exhaustion or enough reads were obtained. Base calling & demultiplexing was performed using Guppy (v6.4.6) using the “SUP” super high accuracy model for R9.4.1 flow cells. The raw reads were filtered for a length between 1500 ± 200 bp. Filtered reads were assigned to taxa using Emu ^72^ (v3.4.4, GitLab Project ID: 19618062) by mapping 16S rRNA sequences to the emu_database database (based on the NCBI 16S RefSeq with the entry for *E. asparagiformis* changed to the sequence obtained by ONT sequencing (GenBank accession PP280819) since the RefSeq sequence for this bacterium contained multiple N nucleotides that biased the assignment of *E. asparagiformis* to *E. lavalensis*). Data were not rarefied or scaled. Count tables were then used to create a phyloseq (v1.44.0, https://github.com/joey711/phyloseq) object in R ^73^. Stacked bar plots were generated using ggnested (v0.1.0, https://github.com/gmteunisse/ggnested) and fantaxtic (v0.2.0, https://github.com/gmteunisse/Fantaxtic). Diversity analyses were performed using Microbiome Analyst (https://www.microbiomeanalyst.ca/) ^74^.

### Total RNA extraction from fecal slurries

A 300 μL fecal slurry aliquot (treated with either 100 µM urolithin C or an equivalent volume DMSO for 48 h) was thawed and pelleted (6,500 g for 3 min). The pellet (in 200 µL media) was then mixed with 800 µL TRI reagent (Zymo Research). Samples were lysed in a Mini Beadbeater 16 (Biospec) according to the following sequence: 1 min ON, 5 min OFF for a total of 5 min ON. RNA isolation was then performed using the Direct-zol RNA Miniprep Kit (Zymo Research) according to the manufacturer’s instructions (including an on-column DNase digestion). To ensure complete DNA removal, an additional DNA digestion step was performed on the isolated RNA using the Ambion DNA-free DNA Removal Kit (Invitrogen) according to the manufacturer’s instructions. The DNA-free RNA was then cleaned up using the OneStep PCR Inhibitor Removal Kit (Zymo Research). RNA concentration and quality were verified by Qubit RNA BR assay kit (Invitrogen) and 1% agarose gel electrophoresis.

### RT-PCR analysis of the *ucd* operon in fecal slurries

Total RNA was extracted from frozen fecal slurries as previously described (see Total RNA extraction from microbial communities), and subsequently reverse transcribed as described above (see *RT-PCR analysis of E. bolteae ucd operon structure*) in a reaction volume of 5 µL. PCR reactions were conducted using the OneTaq 2X Master Mix with Standard Buffer (NEB). The ucdCFO_RT-PCR primer pair was added to the master mix (to a final concentration of 0.2 µM) and 1 µL of template (cDNA, -RT, or no template) was added for a total reaction volume of 20 µL. PCR tubes were placed in a thermal cycler and targets were amplified according to the following conditions: 30s at 94°C, 45 cycles (30s at 94°C, 1 min at 61°C, 4 min at 68°C), 5 min at 68°C. A volume of 10 μL of reaction was directly loaded onto a 1% agarose gel (made with 1X TAE buffer) containing SafeView Classic (Abm). PCR product sizes were compared to the Quick-Load® Purple 1 kb Plus DNA Ladder (NEB).

### PCR analysis of the *ucd* operon prevalence in fecal slurries

Genomic DNA (gDNA) was extracted from frozen fecal slurries as previously described (see Genomic DNA extraction from microbial communities). PCR reactions and product visualization was conducted on the gDNA as described above (see RT-PCR analysis of the *ucd* operon in microbial communities). In this case, 5 µL of PCR product was loaded onto the gels instead of 10 µL.

### Statistical analyses and graphing

Statistical methods were not used to determine sample sizes, experiments were not randomized, and the investigators were not blinded. Data points related to uroC metabolism and RT-qPCR were assumed to be normally distributed, though this was not formally tested. Correlation analyses were performed using the non-parametric Spearman rank correlation (*ρ*). Statistical tests on bacterial relative abundances were performed using the Kruskal-Wallis test on untransformed relative abundance values, which are skewed towards 0. Statistical analyses for large datasets are detailed in the relevant methods sections. Details related to each test performed are supplied in the figure legends. In all cases, α = 0.05 and tests were two-tailed. Data were plotted in GraphPad Prism (v10.0.0) or using the ggplot2 (v3.4.2) R package. Figures were assembled in Affinity Designer (v1.10.6.1665).

## Data Availability

RNA-seq reads were deposited in the NCBI SRA BioProject ID PRJNA996126 under BioSample accession codes SAMN36514640 (*Enterocloster bolteae* DSM 15670) and SAMN36514641 (*Enterocloster asparagiformis* DSM 15981). Reviewer link: https://dataview.ncbi.nlm.nih.gov/object/PRJNA996126?reviewer=fpbuj6eeuebv6mij3pme3mp8ep. Untargeted proteomics data have been deposited to the ProteomeXchange Consortium via the PRIDE partner repository with the dataset identifier PXD048514 and 10.6019/PXD048514 ^75^. Oxford Nanopore 16S rRNA sequencing reads of healthy human fecal slurries were deposited in the NCBI SRA BioProject ID PRJNA1073957. Reviewer link: https://dataview.ncbi.nlm.nih.gov/object/PRJNA1073957?reviewer=nh1a04lg59enlti318qs23s6ot. The 16S rRNA sequence for *E. asparagiformis* DSM 15670 used in the Emu database search was deposited in GenBank under accession PP280819. All original code, tables, and RData files obtained from the analysis of curatedMetagenomicData were deposited in Zenodo (https://doi.org/10.5281/zenodo.8302320).

## Supplementary Figures

**Supplementary Figure 1.**
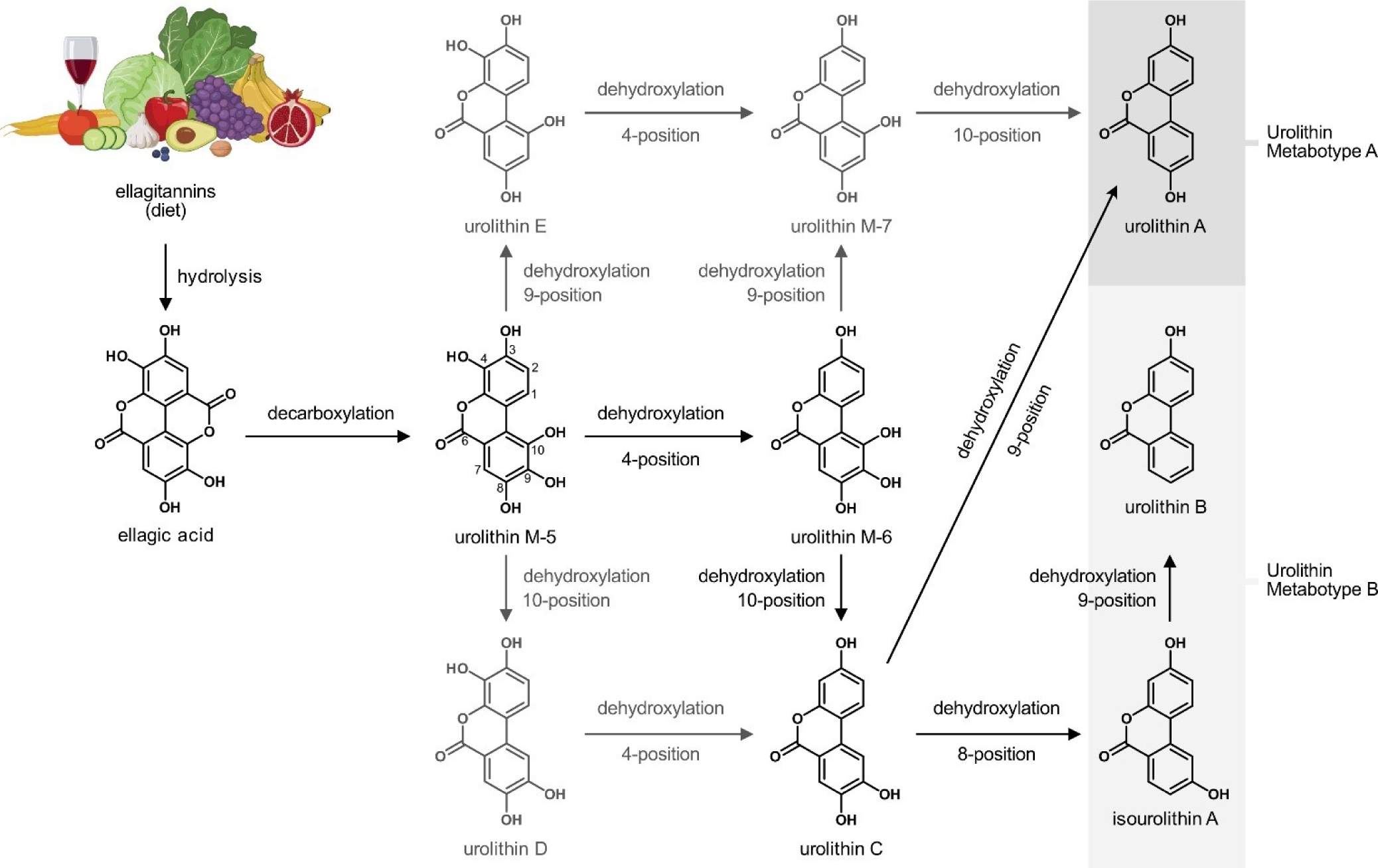
Ellagitannins are metabolized by gut bacteria. Reaction scheme of dietary ellagitannin metabolism by the human gut microbiota. Larger ellagitannin structures are hydrolyzed during gut transit, releasing hexahydroxydiphenic acid, which spontaneously lactonizes into ellagic acid. Once in the gut lumen, members of the *Gordonibacter* spp. and *Ellagibacter isourolithinifaciens* can decarboxylate ellagic acid, forming urolithin M-5. The resulting urolithin M-5 can be further dehydroxylated (at the 4,10- or 4,8,10- positions) to uroC or isourolithin A by *Gordonibacter* spp. or *Ellagibacter isourolithinifaciens*, respectively. Compounds colored in light gray are urolithin metabolites that are rarely observed during *ex vivo* metabolism assays on ellagitannins. Once uroC or isouroA are produced, *Enterocloster* spp. can further dehydroxylate the 9-position, yielding uroA or uroB, respectively. The cartoon was generated in BioRender.

**Supplementary Figure 2.**
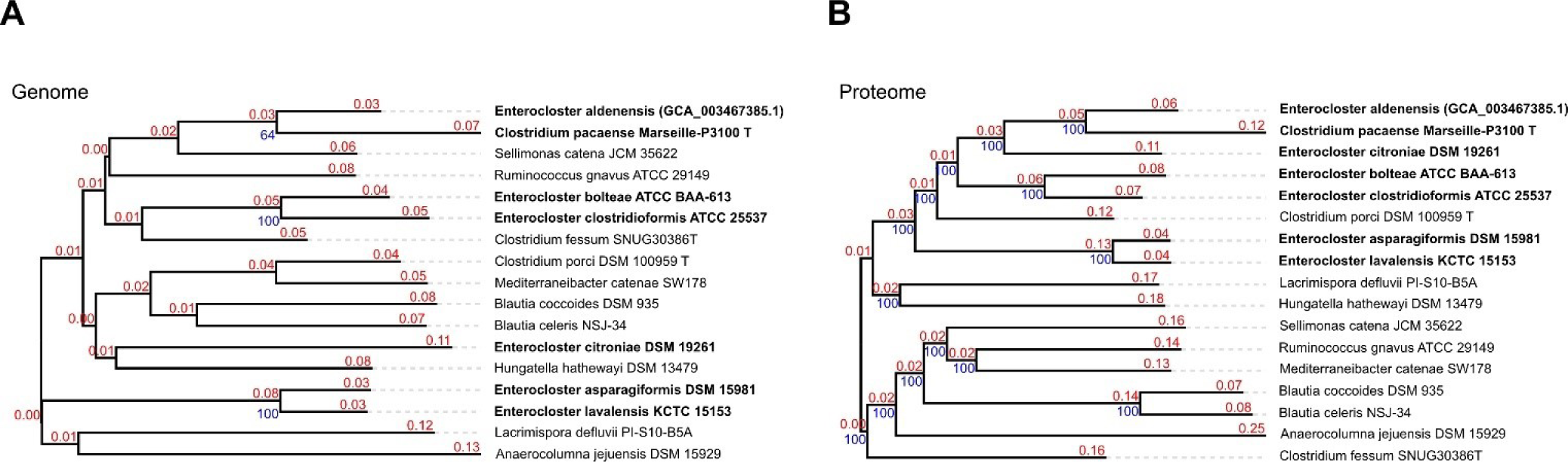
*Enterocloster* spp. whole genome and proteome phylogenetic trees. **A)** Whole genome phylogenetic tree of *Enterocloster* spp. The tree was inferred with FastME 2.1.6.1 from Genome BLAST Distance Phylogeny (GBDP) distances calculated from genome sequences. The branch lengths are scaled in terms of GBDP distance formula d5. The numbers above branches are GBDP pseudo-bootstrap support values > 60 % from 100 replications, with an average branch support of 36.8 %. The tree was rooted at the midpoint. **B)** Whole proteome phylogenetic tree of *Enterocloster* spp. The tree was inferred with FastME 2.1.6.1 from whole- proteome-based GBDP distances. The branch lengths are scaled via GBDP distance formula d5. Branch values are GBDP pseudo-bootstrap support values > 60 % from 100 replications, with an average branch support of 100.0 %. The tree was rooted at the midpoint. Source data are provided as a Source data file.

**Supplementary Figure 3.**
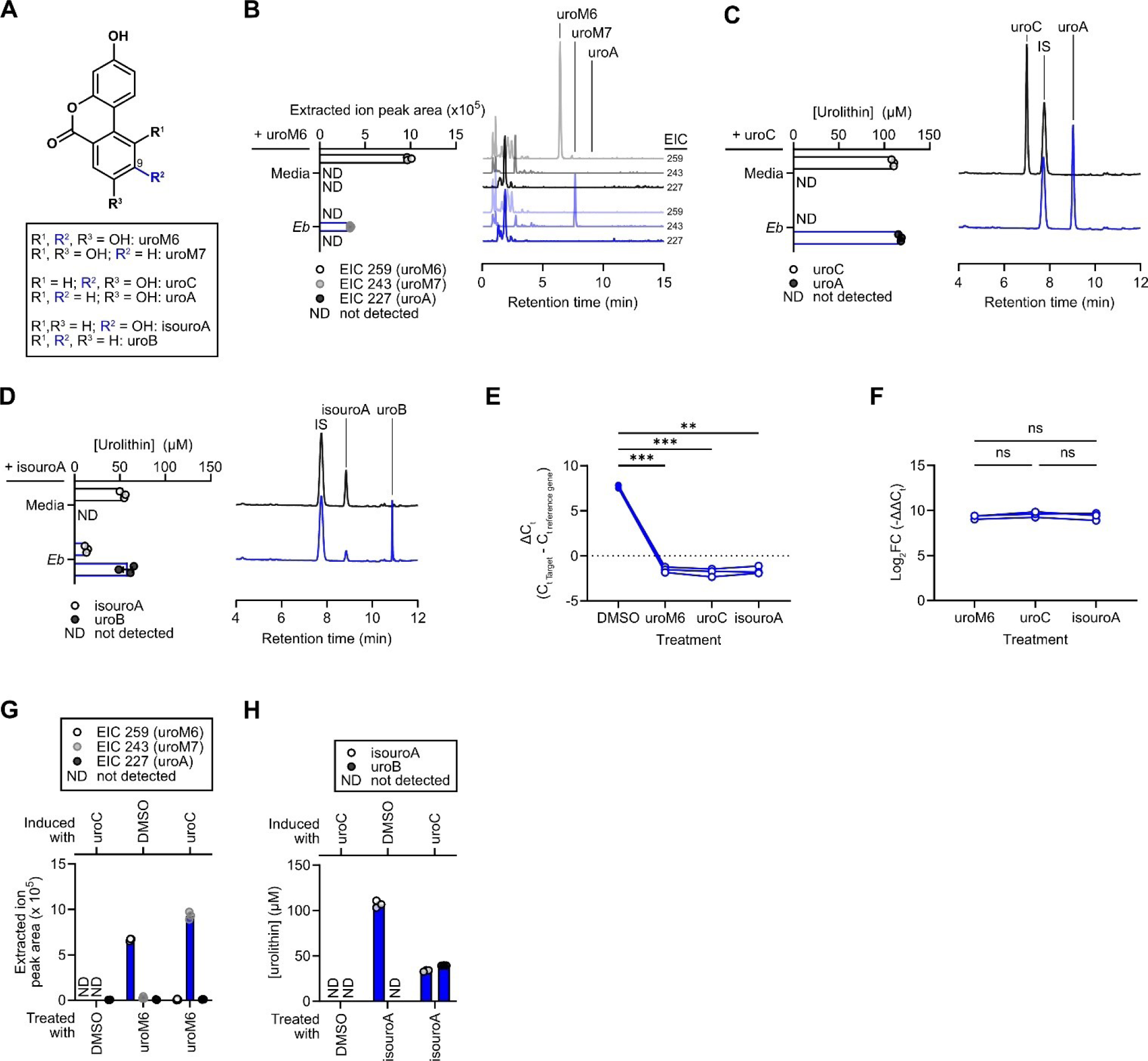
*E. bolteae* dehydroxylates urolithins at the 9-position. **A)** Chemical structures of urolithin M-6, urolithin C, and isourolithin A along with their dehydroxylated counterparts urolithin M7, urolithin A, and urolithin B. **B)** Quantification of extracted ion chromatogram (EIC) peak areas from *Eb* cultures sampled after 24 h of growth with 100 μM uroM6 (n = 3 biological replicates) with representative extracted ion chromatograms (EIC) to the right (from one representative biological replicate). The same scale was used for each chromatogram. **C)** Quantification of urolithin peak areas from *Eb* cultures sampled after 24 h of growth with 100 μM uroC (n = 3 biological replicates) with representative chromatograms (λ = 305 nm) to the right (from one biological replicate). The same scale was used for each chromatogram. **D)** Quantification of urolithin peak areas from *Eb* cultures sampled after 24 h of growth with 100 μM isouroA (n = 3 biological replicates) with representative chromatograms (λ = 305 nm) to the right (from one biological replicate). The same scale was used for each chromatogram. **E,F)** RT-qPCR expression of the *Eb ucdO* gene. Growing *Eb* cultures were treated with DMSO, uroM6, uroC, or isouroA (100 μM) for 2 h before RNA isolation and reverse transcription (n = 3 biological replicates). **E)** Differential *Eb ucdO* gene expression comparing DMSO, uroM6, uroC, and isouroA is displayed as target-specific ΔCt (C_t MoO Gene_ - C_t dnaK Reference Gene_) values. Data are presented as individual ΔCt values with lines connecting paired biological replicates (from the same pre-spike culture); repeated- measures one-way ANOVA with Dunnett’s multiple comparisons test; **, p < 0.01; ***, p < 0.001. **F)** Gene expression profile of the *Eb ucdO* gene in different urolithin treatment groups displayed as log_2_FC (equivalent to -ΔΔC_t_, where ΔΔC_t_ = ΔC_t urolithin_ - ΔC_t DMSO_). Data are presented as individual log_2_FC values with lines connecting paired biological replicates; repeated-measures one-way ANOVA with Tukey’s multiple comparisons test. **G)** Quantification of extracted ion chromatogram (EIC) peak areas in DMSO- or uroC-treated *Eb* cell suspensions. Cell suspensions were prepared from *Eb* cells grown with either DMSO or 50 μM uroC. The cells were washed and resuspended in PBS to halt the production of new enzymes, then treated with 100 μM uroM6 (n = 3 biological replicates). **H)** Quantification of urolithin concentrations in DMSO- or uroC-treated *Eb* cell suspensions (n = 3 biological replicates). For B-D and G-H, data are represented as mean ± SEM. ND, not detected; ns, not significant; FC, fold change; FDR, false discovery rate. Source data and statistical details are provided as a Source data file.

**Supplementary Figure 4.**
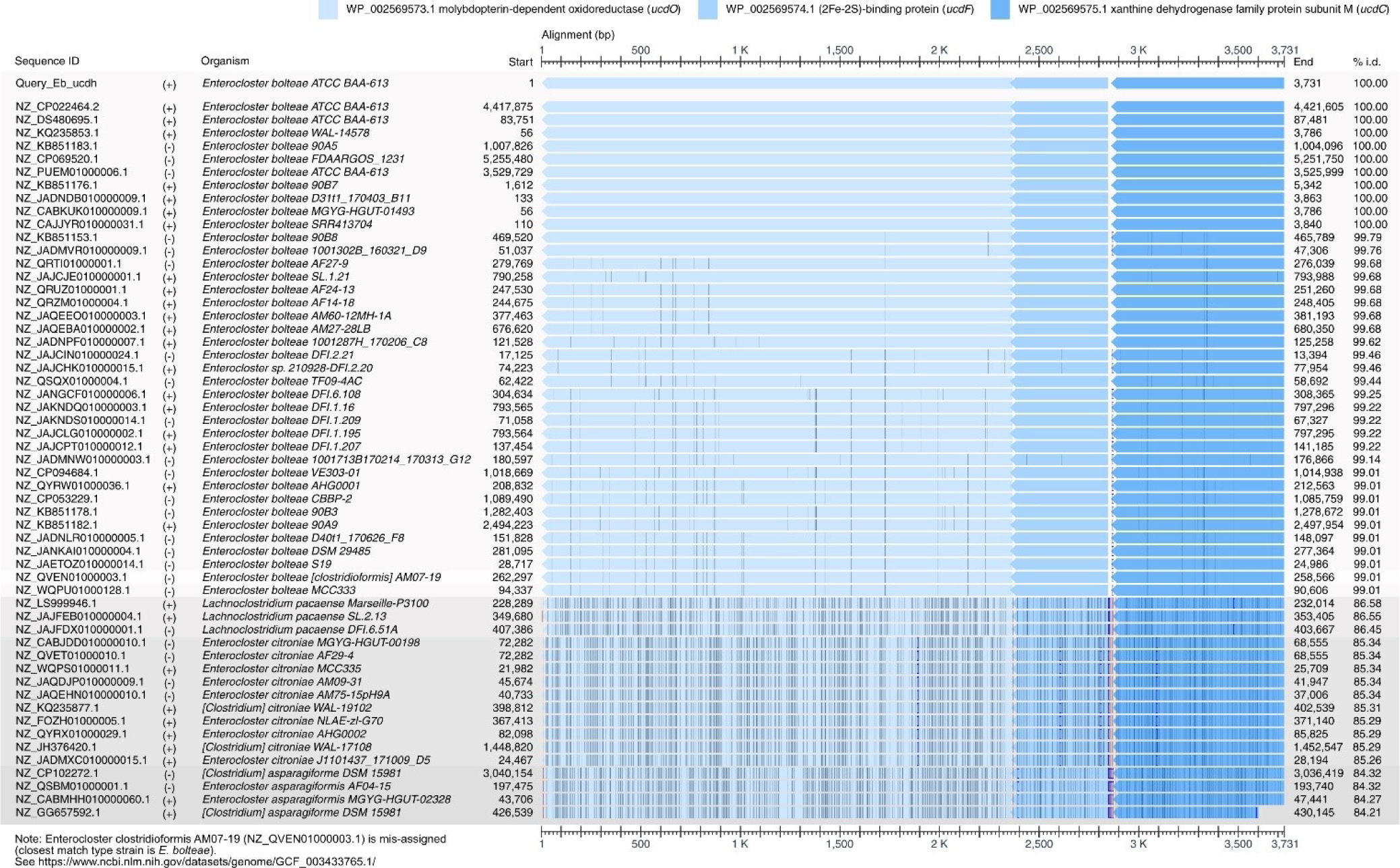
BLASTn searches using the *E. bolteae ucd* operon genomic sequence identifies homologues in gut bacteria. NBCI Multiple Sequence Aligner viewer hits for BLASTn searches using the *E. bolteae* DSM 15670 *ucd* operon nucleotide sequence as a query against the NCBI refseq_genomes database (limited to Bacteria). Vertical lines in the sequence alignment represent nucleotide differences (show differences option) and insertions relative to the query sequence. Source data are provided as a Source data file.

**Supplementary Figure 5.**
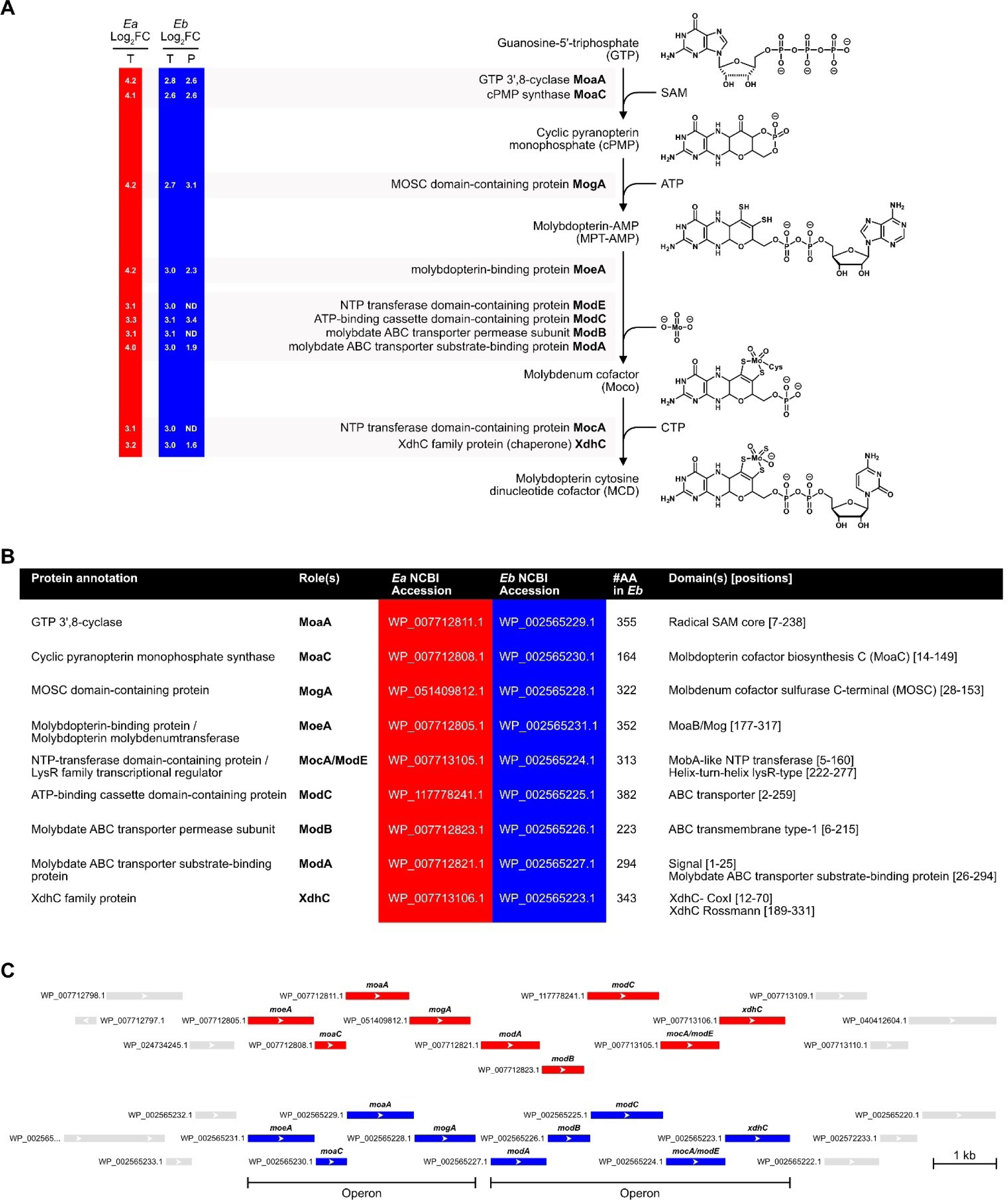
Urolithin C treatment upregulates molybdopterin cytosine dinucleotide cofactor biosynthetic gene clusters. **A)** Molybdopterin cytosine dinucleotide (MCD) cofactor biosynthetic pathway. The log_2_FC values of MCD cofactor biosynthetic genes upregulated by uroC in both transcriptomics (T) and proteomics (P) datasets (when available) are provided to the left of the figure for both *Ea* and *Eb*. **B)** Table of molybdenum cofactor biosynthetic genes found in the genomes of *Ea* and *Eb*. Annotations are based on NCBI (GTF files) and UniProt gene/protein names. Roles are assigned based on required proteins for molybdenum cofactors in the xanthine oxidase/dehydrogenase family of enzymes. Accessions, primary sequence length, and annotated domains (with positions within the primary sequence) are also provided. **C)** Genomic organization of the MCD cofactor biosynthetic genes for *Ea* and *Eb* (generated from the NCBI Sequence Viewer). Source data are provided as a Source data file.

**Supplementary Figure 6.**
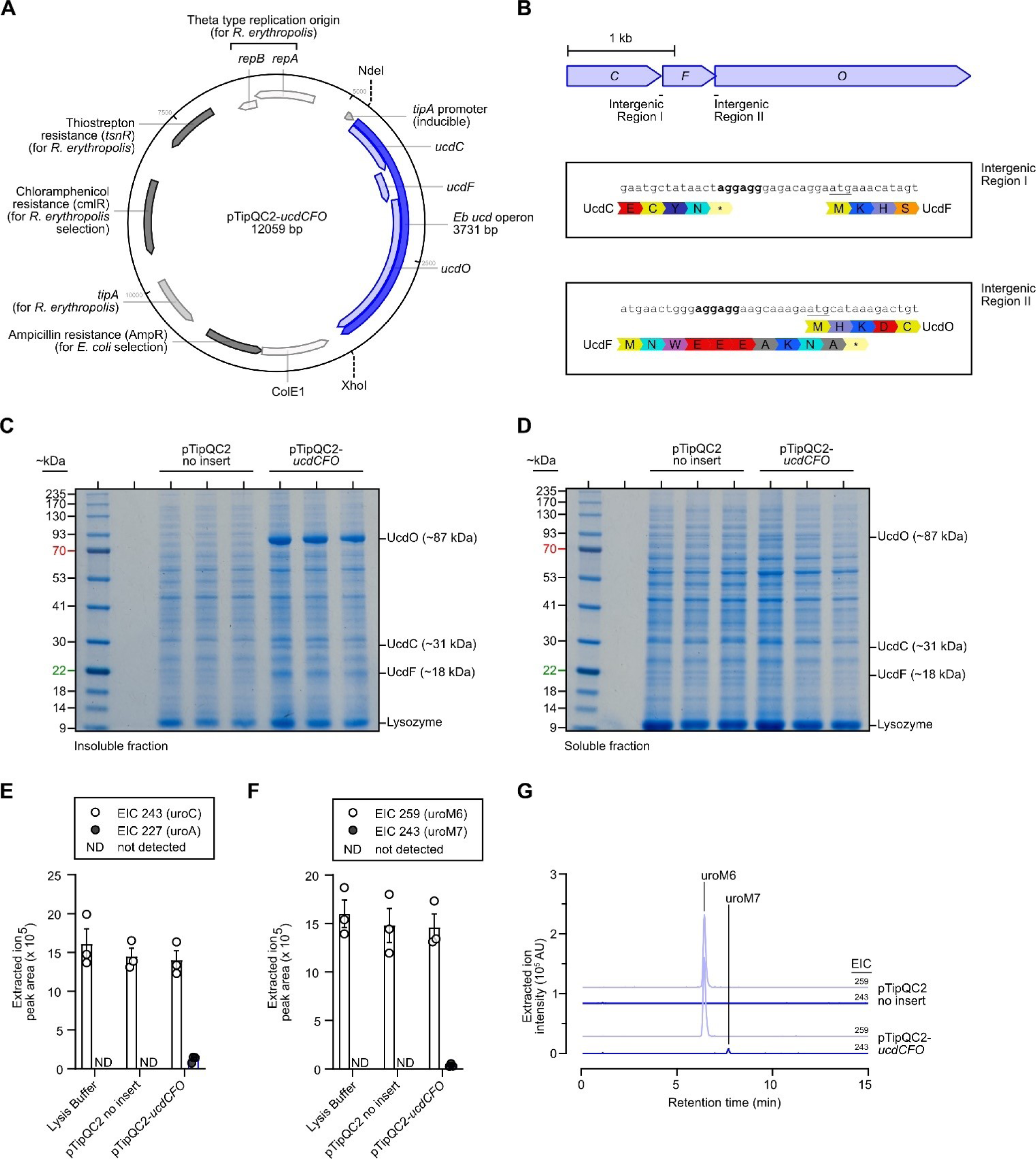
Heterologous expression of *E. bolteae ucdCFO* genes in *R. erythropolis*. **A)** Map of the *E. coli* – *R. erythropolis* pTipQC2-*ucdCFO* shuttle plasmid generated in Benchling. **B)** Genomic organization of the wild-type *E. bolteae ucd* operon with intergenic regions highlighted. Shine-Dalgarno consensus sequences (bold) are denoted between translational stop and start (underlined) sites for *ucdC* and *ucdF* (Intergenic Region I) and for *ucdF* and *ucdO* (Intergenic Region II). **C,D)** SDS-PAGE gels (10% bis-tris) stained with colloidal Coomassie dye of the insoluble (C) and soluble (D) fractions from thiostrepton-induced (1 μg/mL) *R. erythropolis* harboring pTipQC2 no insert or pTipQC2-*ucdCFO* plasmids (n = 3 biological replicates). UcdCFO complex proteins are labeled on the right side of each gel image. **E,F)** Quantification of extracted ion chromatogram (EIC) peak areas from crude lysates of thiostrepton-induced *R. erythropolis* harboring pTipQC2 no insert or pTipQC2-*ucdCFO* plasmids (n = 3 biological replicates). Crude lysates were incubated anaerobically with 2 mM NADH and 357 μM uroC (E) or 357 μM uroM6 (F) for 72 h before extraction and analysis by LC-MS. Data are represented as mean ± SEM. **G)** LC-MS extracted ion chromatograms (EIC) of uroM6 ([M-H]^-^ = 259) and uroM7 ([M-H]^-^ = 243) from a representative anaerobic uroC dehydroxylation assay using crude lysates of *R. erythropolis* harboring either pTipQC2 (no insert) or pTipQC2-*ucdCFO* plasmids. Source data are provided as a Source data file.

**Supplementary Figure 7.**
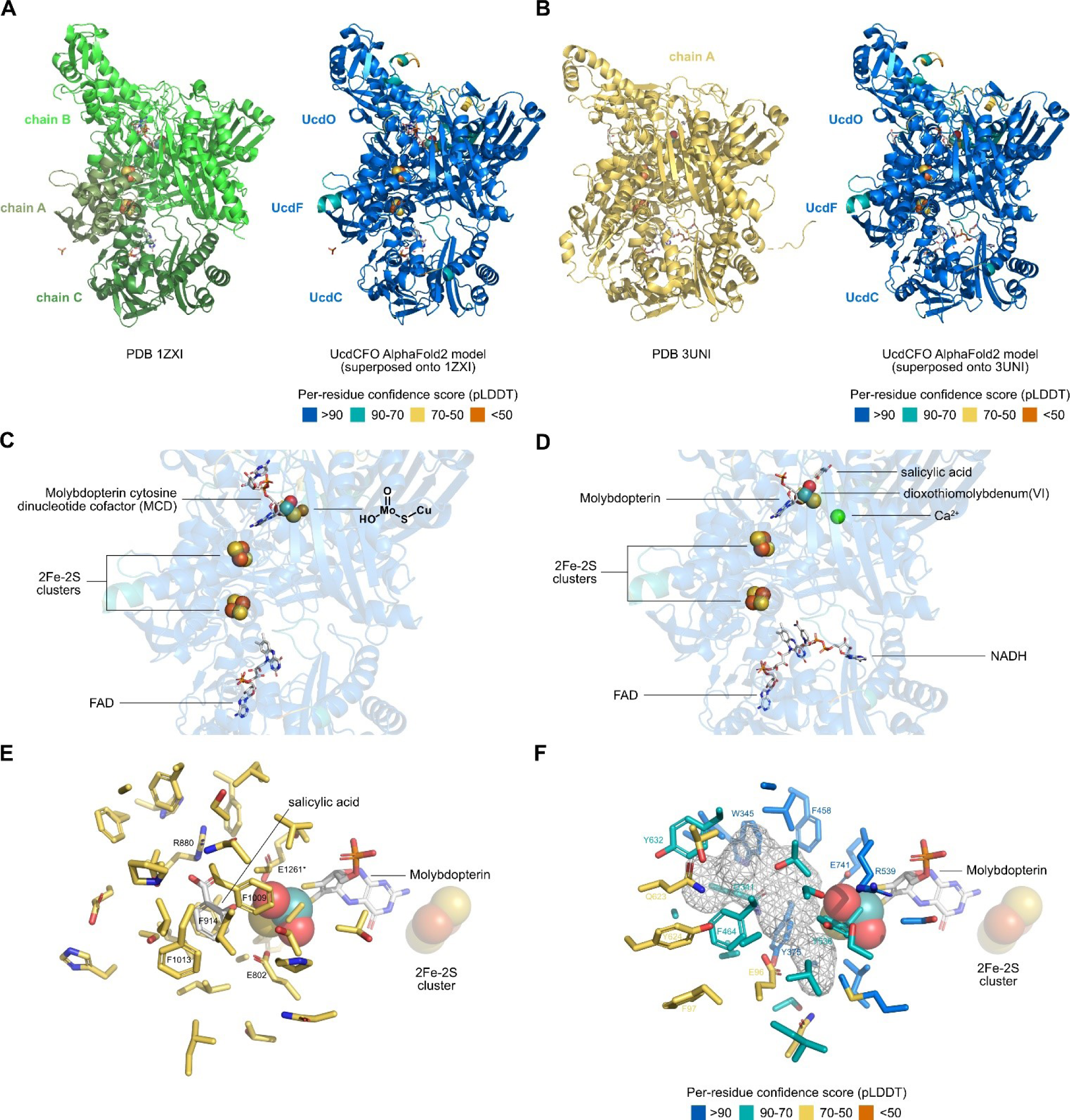
The AlphaFold2 model of the *E. bolteae* UcdCFO enzyme complex has a similar quaternary structure to xanthine dehydrogenase superfamily crystal structures. A,B) Structural superposition of AlphaFold2-predicted *Eb* Ucd proteins (right) onto the X-ray crystal structures of **A)** PDB 1ZXI (carbon monoxide dehydrogenase from *Oligotropha carboxidovorans* OM5) X-ray crystal structure (left) and **B)** PDB 3UNI (bovine milk xanthine dehydrogenase with NADH bound). PDB 1ZXI is colored in shades of green (according to chain ID), PDB 3UNI is colored in gold (single chain A shown), and the *Eb* Ucd enzyme complex is colored according to its per-residue confidence score as indicated in the legend. The various ligands (cofactors, coenzymes, ions, small molecules) of PDB 1ZXI and PDB 3UNI are included in the respective *Eb* Ucd enzyme models. **C,D)** Cofactors from X-ray crystal structures **C)** PDB 1ZXI and **D)** PDB 3UNI modeled into the AlphaFold2 *Eb* Ucd enzyme complex, showing a complete electron transport chain from a bound FAD molecule to a molybdopterin cofactor *via* two 2Fe-2S clusters. **E)** Xanthine dehydrogenase active site from PDB 3UNI. Side chains within 8 Å of the salicylic acid ligand are colored in gold. Residues important for substrate (purine) binding and catalysis are labeled with their one letter amino acid code and sequence position. Position E1261* is catalytically important (acts as a general base) and is conserved in XDH/xanthine oxidase enzymes ^59^. **F)** *Eb* Ucd enzyme complex active site modeling. Side chains within 8 Å of the salicylic acid ligand from PDB 3UNI are colored on the superposed *Eb* Ucd enzyme complex according to their per-residue confidence score as indicated in the legend. Residues surrounding the predicted active site are labeled with their one letter amino acid code and sequence position (in the *Eb* MoO protein). The predicted urolithin binding site is depicted by the surface (mesh) created by the active site residues. The surface was rendered using the cavities and pockets only (culled) setting with a cavity detection cutoff of 5 solvent radii in PyMOL. Source data are provided as a Source data file.

**Supplementary Figure 8.**
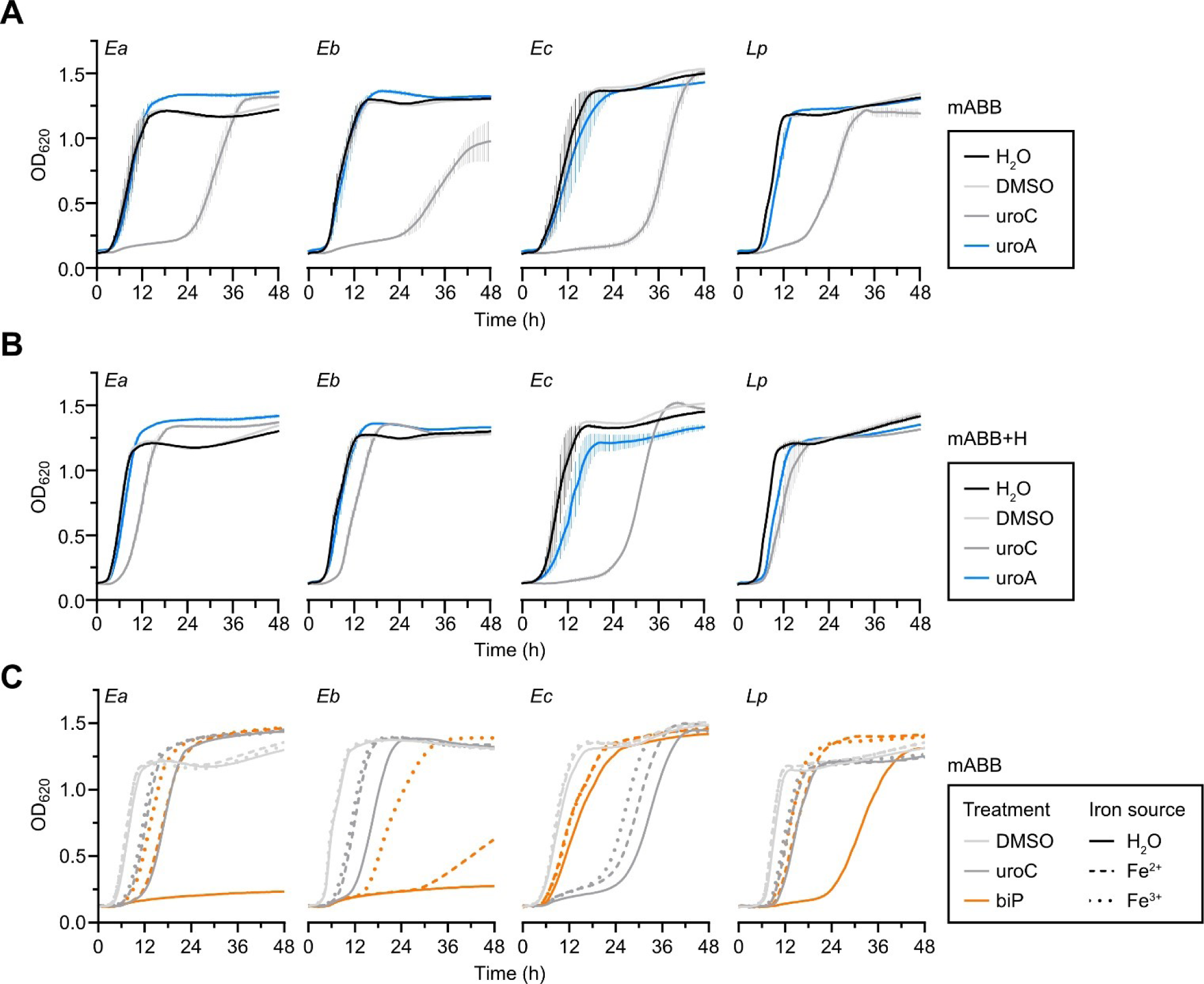
Iron supplementation rescues lag time extension by uroC and 2,2’-bipyridyl. A,B) Growth curves (optical density (OD) at 620 nm) of uroC-metabolizing *Enterocloster* spp. and *L. pacaense* treated with H_2_O, DMSO (vehicle), or 100 μM of uroC or uroA in mABB medium (lacking added iron) (A) or mABB+H (B) medium (containing 7.7 μM hemin) (n = 3 biological replicates). Data are represented as mean ± SEM. **C)** Growth curves (optical density (OD) at 620 nm) of uroC-metabolizing *Enterocloster* spp. and *L. pacaense* treated with DMSO (vehicle), 100 μM of uroC, or 2,2’-bipyridyl (biP) in mABB media (lacking added iron) supplemented with solutions containing no iron (H_2_O), 7.7 μM Fe^2+^ (Fe(II)SO_4_), or 7.7 μM Fe^3+^ (Fe(III) pyrophosphate) (n = 3 biological replicates). Data are represented as means without error bars for clarity. Source data are provided as a Source data file.

**Supplementary Figure 9.**
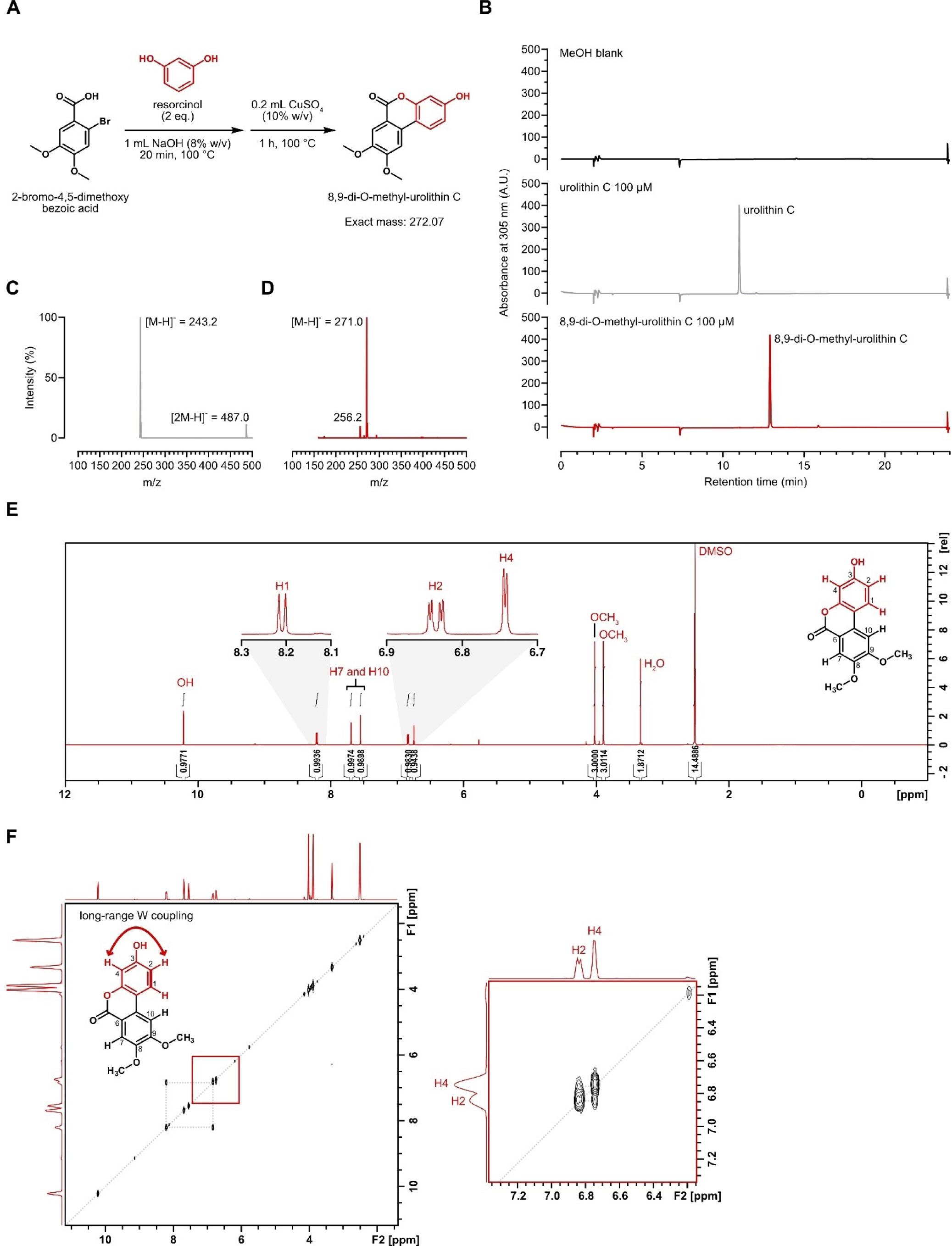
Synthesis and characterization of 8,9-di-O-methyl-urolithin C A) Reaction scheme for the synthesis of 8,9-di-O-methy-urolithin C. **B)** Aligned reversed-phase (C18) HPLC chromatograms (λ = 305 nm) of 10 μL injections of the following solutions: MeOH blank, urolithin C 100 μM, 8,9-di-O-methy-urolithin C 100 μM. **C,D)** Negative ESI-MS spectra of urolithin C (C) and 8,9-di-O-methy-urolithin C (D) following chromatographic separation (B). **E)** ^1^H NMR spectrum (600 MHz, (CD_3_)_2_SO)) of 8,9-di-O-methy-urolithin C. **F)** COSY NMR spectrum (600 MHz, (CD_3_)_2_SO)) of 8,9-di-O-methy-urolithin C. The grey diagonal line denotes self correlation between protons. The right panel corresponds to the area in the red box. Coupling between protons is shown on the 8,9-di-O-methy-urolithin C using bold lines and bold arrows (for long range coupling).

**Supplementary Figure 10.**
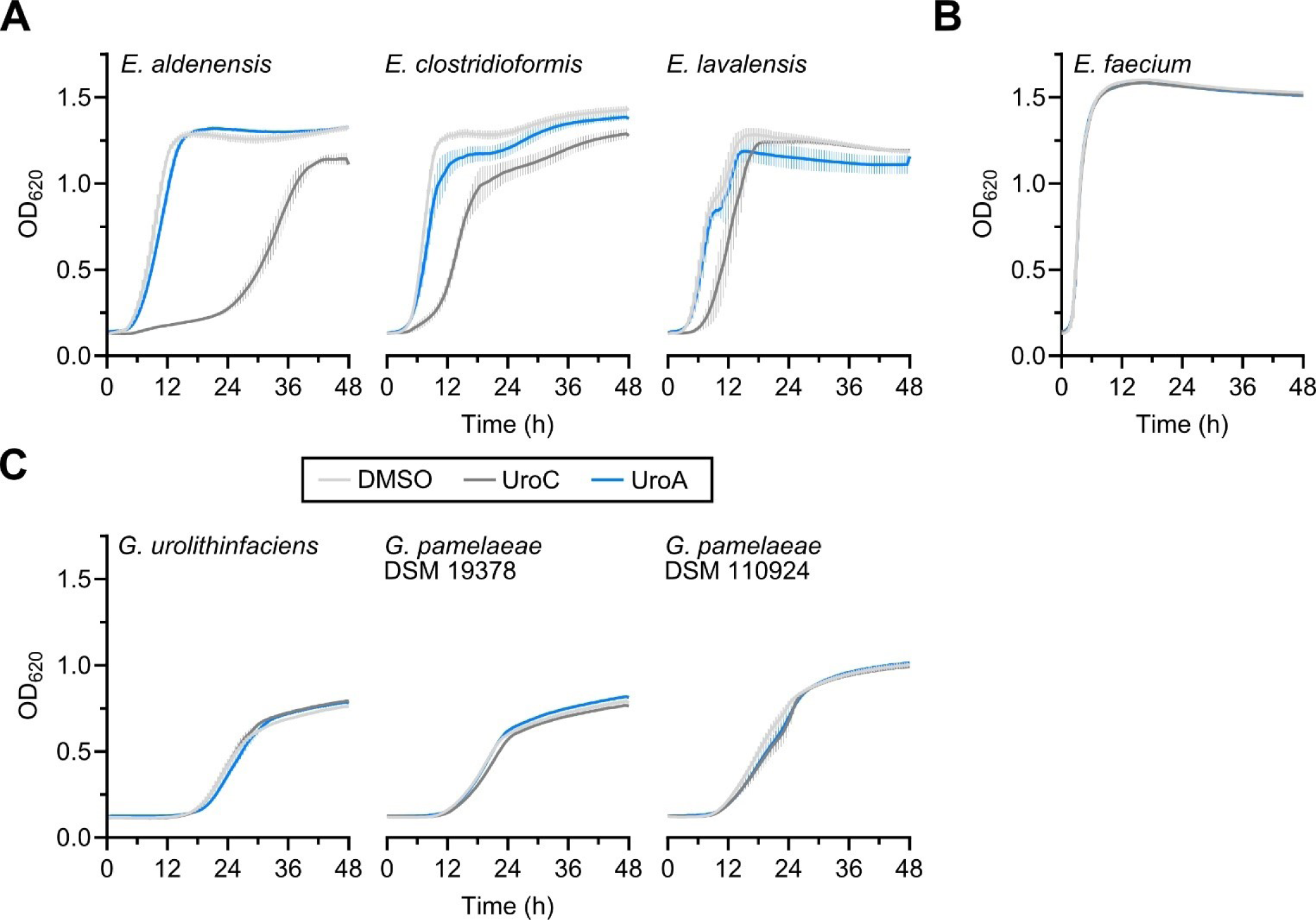
Urolithin C differentially affects the growth of gut bacteria *in vitro*. A,B,C) Growth curves (optical density (OD) at 620 nm) of non-uroC metabolizing *Enterocloster* spp. (A), *Enterococcus faecium* (B) and *Gordonibacter* spp. (C) treated with DMSO (vehicle), or 100 μM of uroC or uroA in mABB medium (n = 3 biological replicates for *Enterocloster* spp. and *E. faecium*, and n = 4 biological replicates for *Gordonibacter* spp.). Data are represented as mean ± SEM.

**Supplementary Figure 11.**
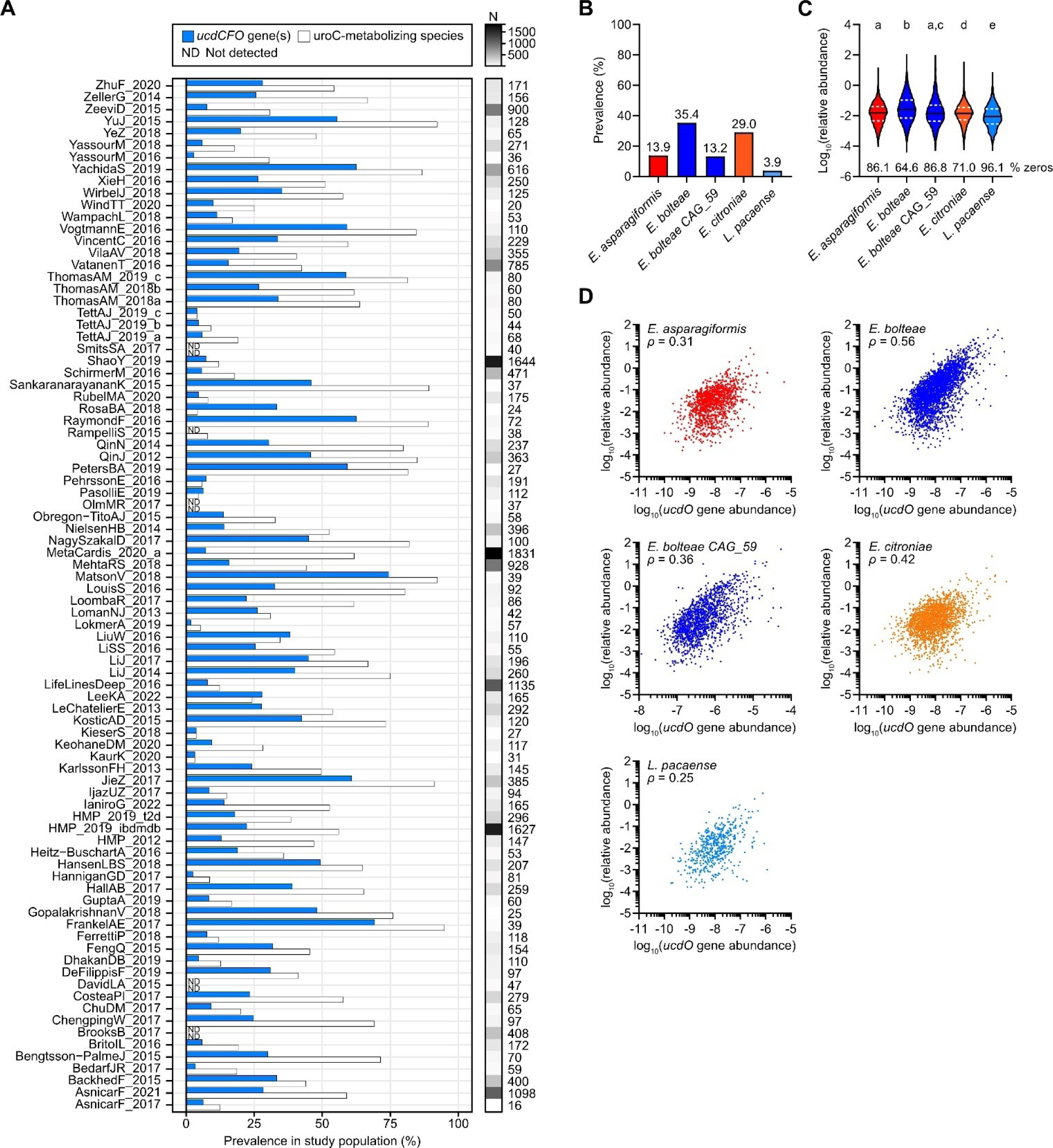
Urolithin C-metabolizing species and genes are prevalent and correlate with *ucdO* gene abundance in human gut metagenomes. **A)** Related to Fig. 6A. Prevalence of at least one *ucd* operon gene (blue bars) and uroC- metabolizing species (white bars) in fecal metagenomes across all 86 studies (in reverse alphabetical order). The number of participants in each study are represented to the right of the prevalence plot as a heatmap. Details on the study populations (from the curatedMetagenomicData R package) can be found in the Source Data file. **B)** Prevalence of uroC-metabolizing species in human fecal metagenomes from the curatedMetagenomicData R package. Data are reported for 86 studies (N=21,030 individuals) and are colored according to the species. **C)** Violin plot of the log_10_(relative abundance) of uroC-metabolizing species in human fecal metagenomes. The solid horizontal line corresponds to the median and the dashed white lines correspond to the first and third quartiles. The percentage of zeroes are denoted below the plotted distributions. Differences between groups were determined using the Kruskal- Wallis test on untransformed relative abundance values. Significant differences between groups are denoted by a different lowercase letter above each plot. **D)** Correlation between the *ucdO* gene abundance in reads per kilobase per million mapped reads (RPKM) and the relative abundance of each uroC-metabolizing species in fecal metagenomes. Both values are illustrated on a log_10_ scale. Spearman rho (*ρ*) values are denoted above the scatter plots. All correlations were significant, P < 0.0001. Source data are provided as a Source data file.

**Supplementary Figure 12.**
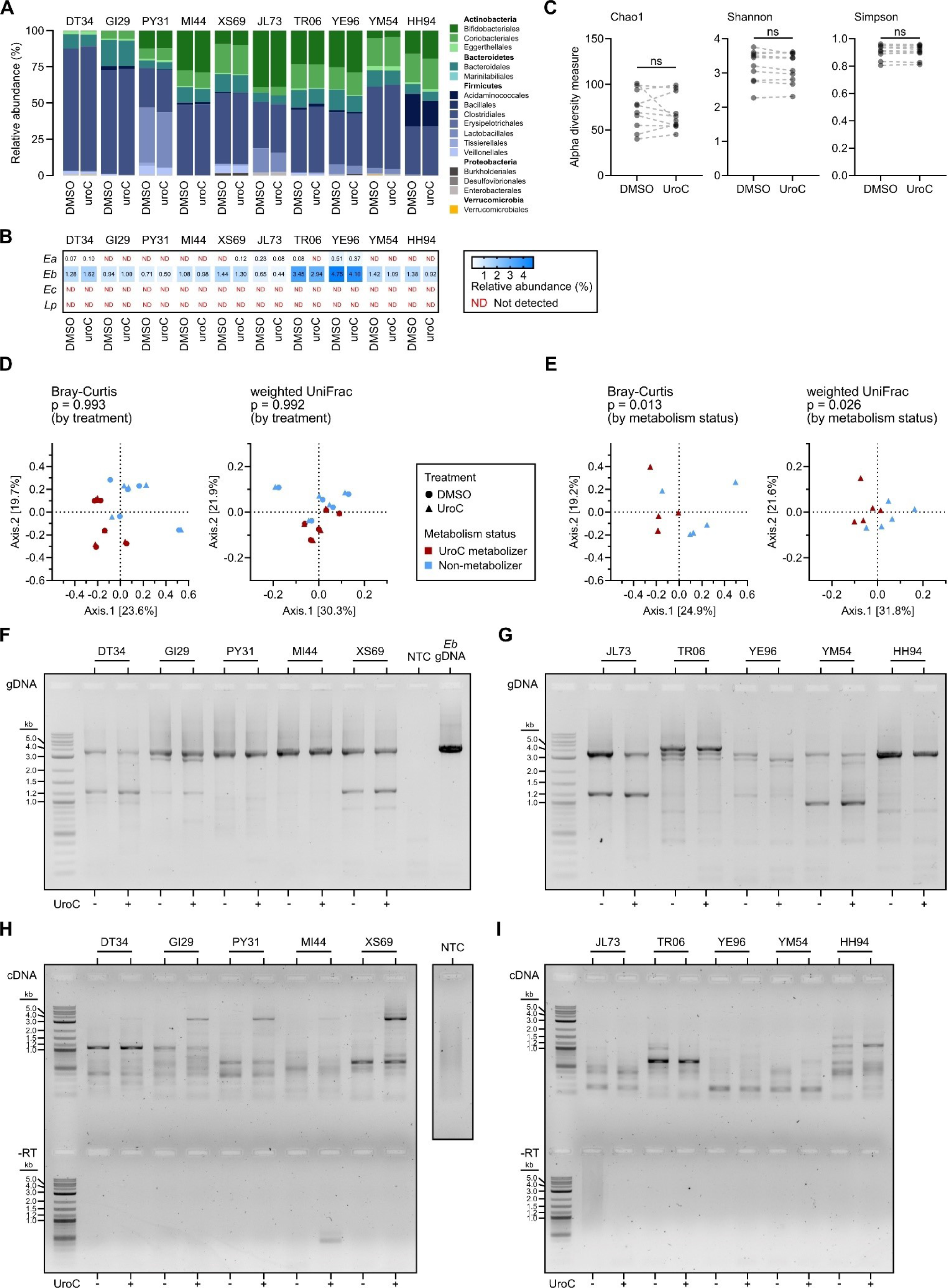
Urolithin C-metabolizing species and the *ucd* operon are prevalent in fecal slurries, but only *ucd* transcription correlates with urolithin C metabolism. **A)** Stacked bar plot of bacterial percent relative abundance (based on V1-V9 16S rRNA gene sequencing) in DMSO- or uroC-treated fecal slurries from 10 healthy donors (from one experimental replicate). Bars are colored according to the phylum (bold) and order. **B)** Heatmap of the percent relative abundance of uroC-metabolizing *Enterocloster* spp. in A). **C)** Alpha diversity plots between DMSO- and uroC-treated fecal slurries according to Chao1, Shannon, and Simpson diversity metrics. Lines between data points connect paired biological replicates; Wilcoxon test; ns, not significant. **D,E)** Principal coordinate analyses of dissimilarities between 16S rRNA compositions based on the Bray-Curtis and weighted UniFrac distance methods. Data points are colored according to the treatment used and the metabolism status of the fecal slurry; PERMANOVA test according to the treatment (D) for all fecal slurries or uroC metabolism status (E) for uroC-treated slurries. **F,G)** PCR with *ucd*-specific primers on gDNA extracted from an *E. bolteae* isolate and fecal microbiota communities from 10 healthy donors (from one experimental replicate). 1% agarose gel of amplicons derived from uroC-metabolizing (F) and non-metabolizing (G) fecal slurries. Samples are matched to the urolithin metabolism data in Fig. 6C and 16S rRNA sequencing in A) and B). The no template control (NTC) and *E. bolteae* positive control are the same for both gels. **H,I)** 1% agarose gel of amplicons from Fig. 6D,E including the no reverse transcriptase (-RT) in uroC-metabolizing (H) and non-metabolizing (I) fecal slurries. The NTC is derived from the final lane of the gel in H). Source data are provided as a Source data file.

**Supplementary Sequence 1.**
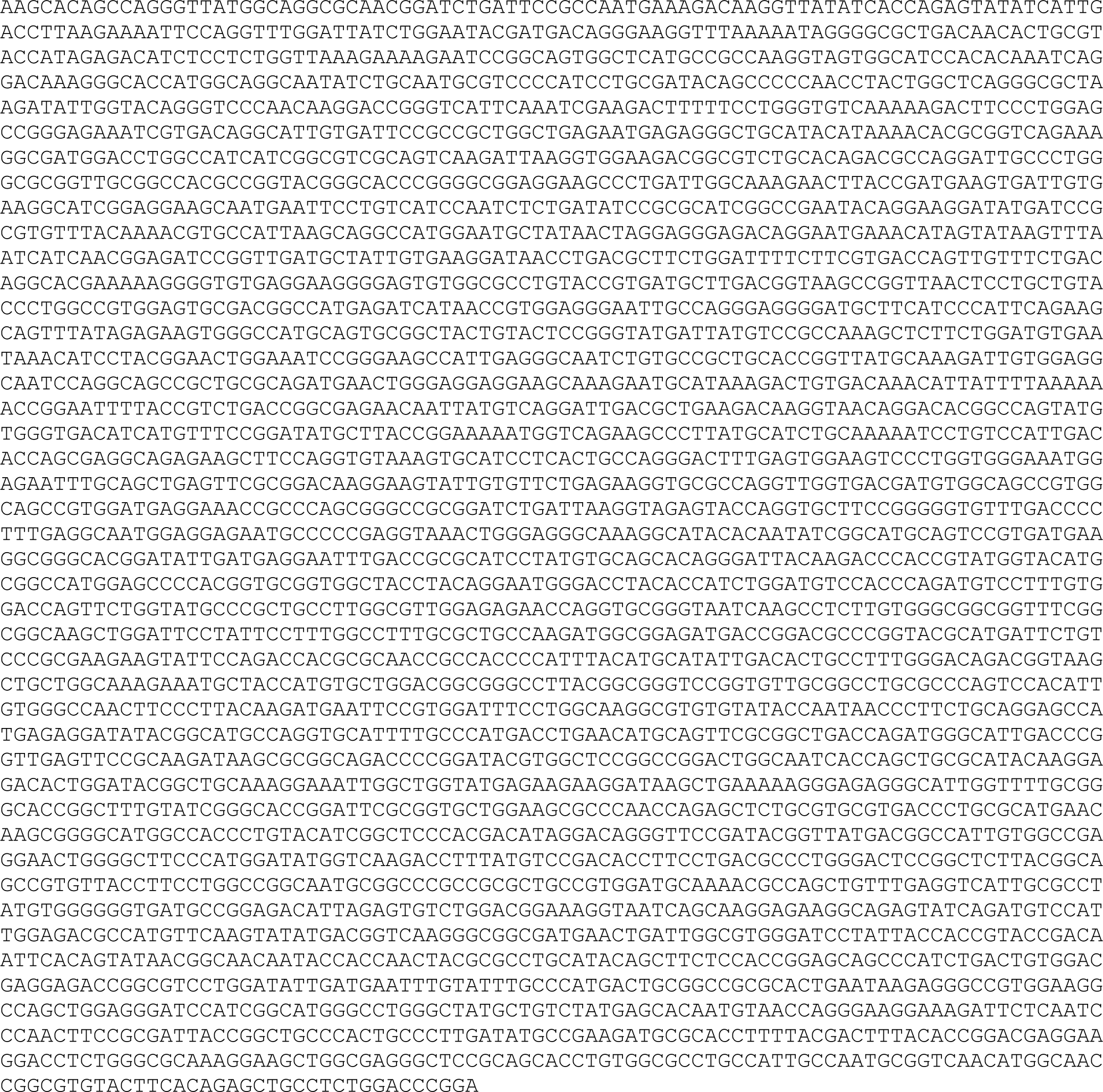
cDNA sequence of *E. bolteae ucdCFO* transcript, 3605 bp (band in Fig. 2I) >Eb_ucd_RT-PCR_band coverage: 4.94e+03x

**Supplementary Sequence 2.**
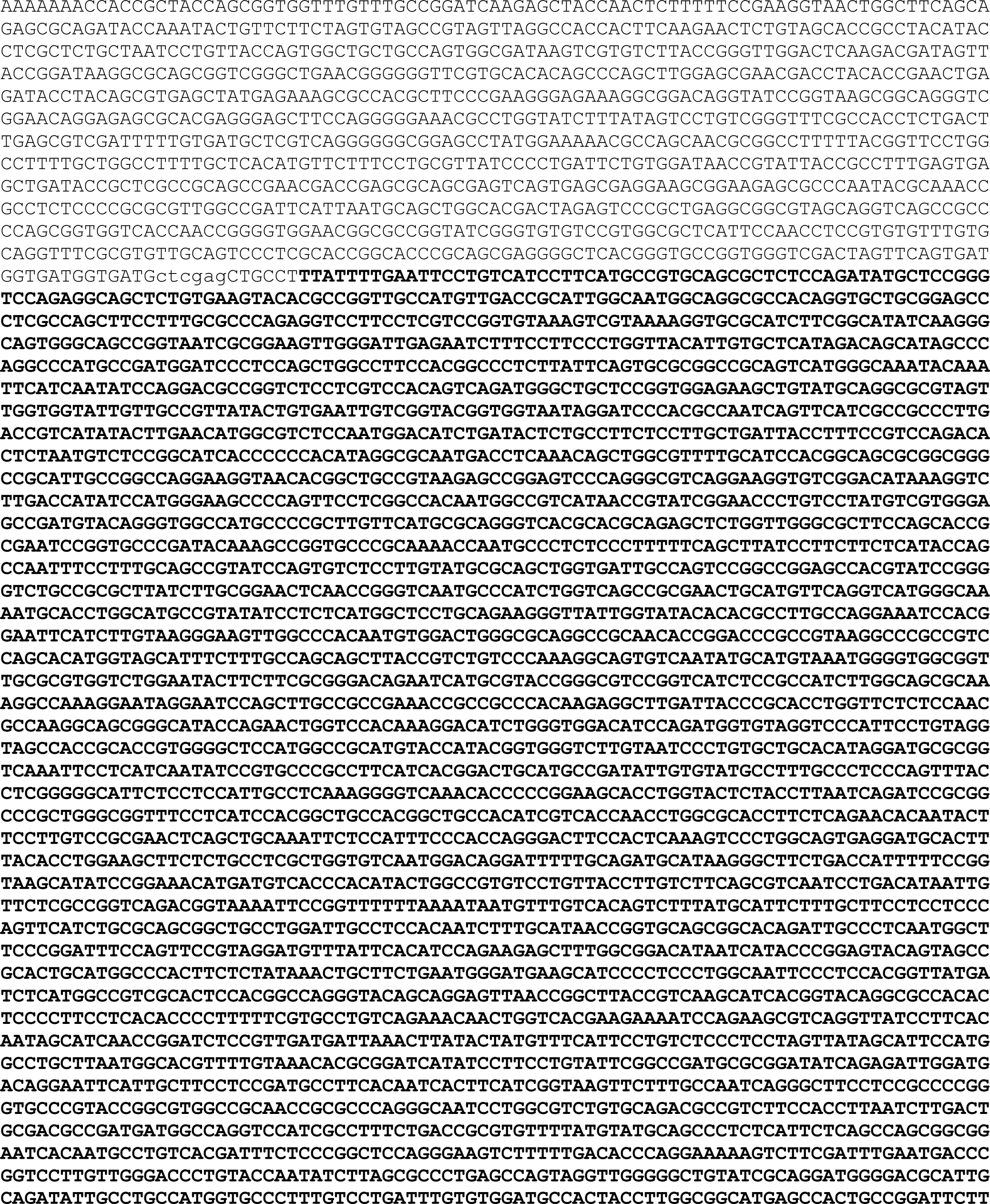
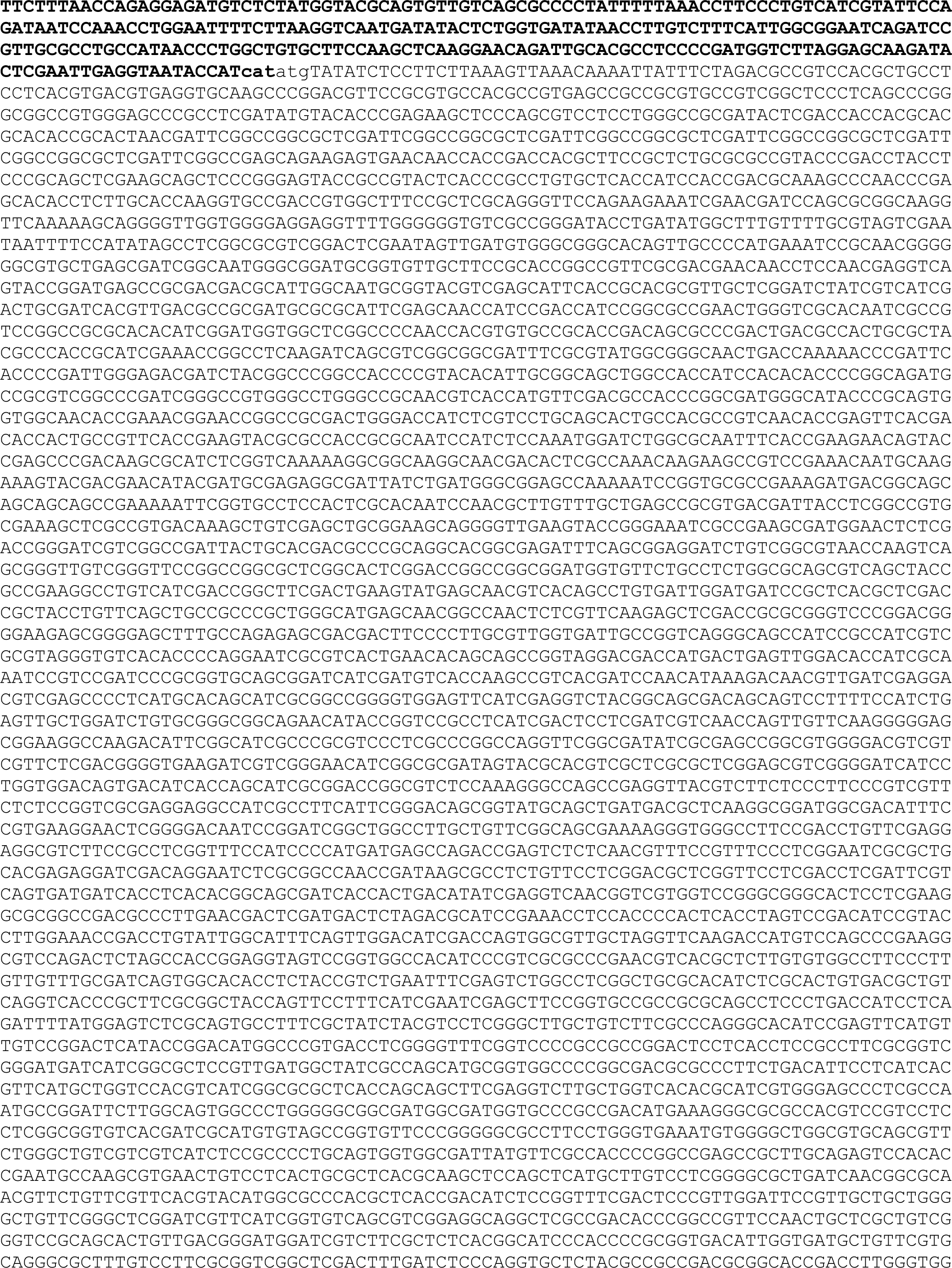
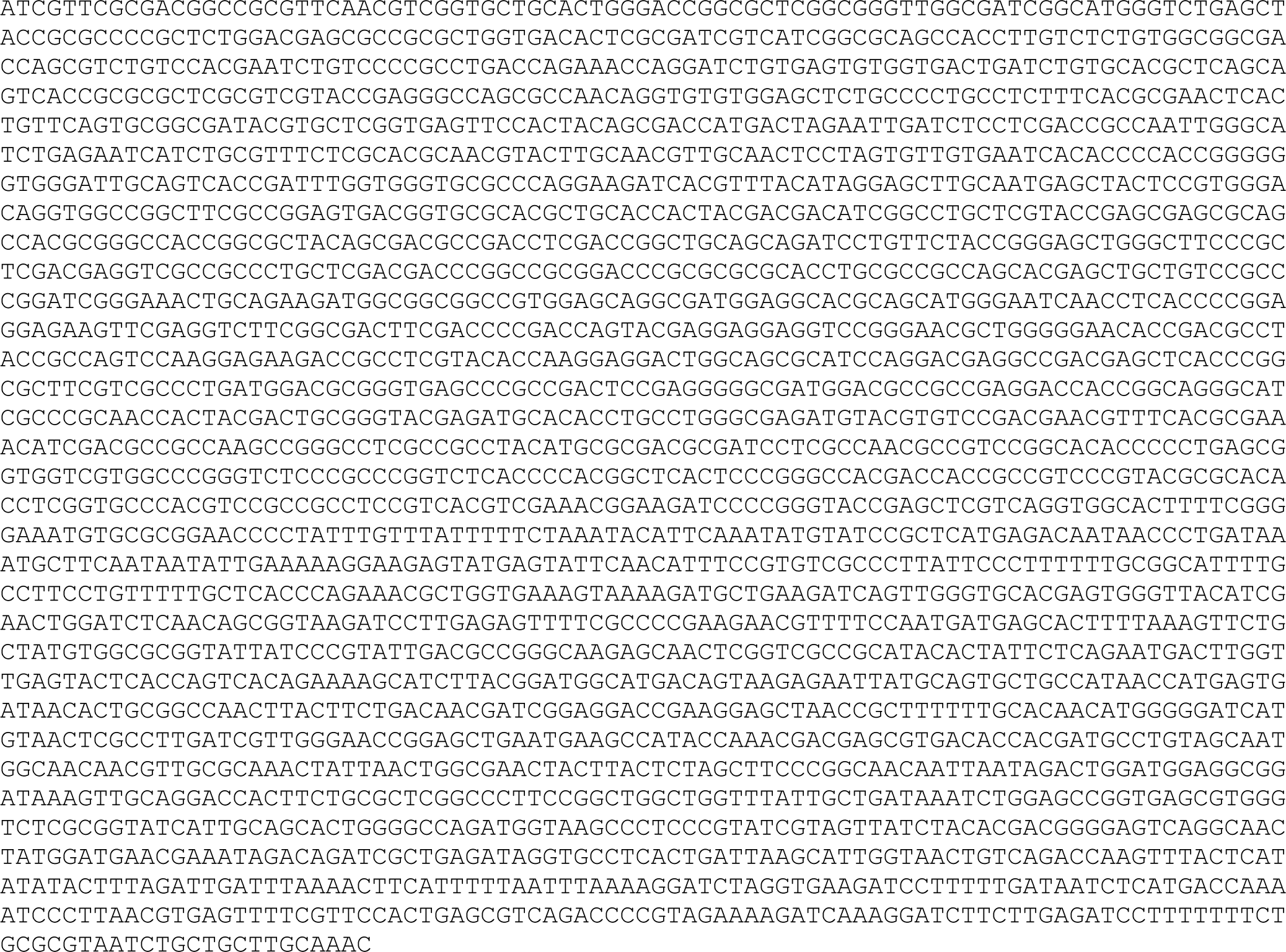
pTipQC2-ucdCFO. >pTipQC2-ucdCFO coverage: 717x NdeI and XhoI restriction sites are lowercase and the *ucdCFO* insert is in bold.

## Notes

https://doi.org/10.5281/zenodo.8302320

### Competing Interest Statement

The authors have declared no competing interest.

### Summary of Updates

Added data related to heterologous expression and growth kinetics in the presence of metabolites.

